# Signatures of brain-wide functional maturation in larval zebrafish

**DOI:** 10.64898/2026.10.14.759119

**Authors:** Antoine Légaré, Mohamed Amine El Barkat, Marc Lebordais, Hugo Poulin, Mado Lemieux, Patrick Desrosiers, Paul De Koninck

## Abstract

Brain development involves the coordinated organization of cellular activity into distributed functional networks, yet how this organization emerges across an entire vertebrate brain remains poorly understood. Here, we combined brain-wide structural and functional imaging to characterize brain maturation in larval zebrafish from 4 to 8 days post-fertilization. Brain growth was regionally heterogeneous and accompanied by progressive remodeling of mesoscale functional networks, with functional connectivity strengthening or weakening within distinct subnetworks. Spontaneous activity became more correlated in regions exhibiting pronounced neurogenesis, as revealed by HaloTag-based neuronal birthdating. Functional interactions became progressively less constrained by physical distance, while network organization became more modular, suggesting that long-range functional integration develops alongside increasing segregation of large-scale brain systems. At the cellular level, neuronal activity consolidated into larger, more modular functional ensembles whose combinations increasingly differentiated mesoscale modules. Visually responsive neurons redistributed along visual processing pathways, becoming more prevalent in regions further removed from retinal input, while motor-related activity declined broadly across the brain. The forebrain emerged as a major locus of maturation, exhibiting substantial anatomical expansion and a growing population of neurons displaying intrinsic activity fluctuations distinct from sensory and motor processes. Together, these findings reveal coordinated anatomical, cellular, and network-level signatures of early vertebrate brain maturation.

## Introduction

Developing nervous systems must become functional long before their construction is complete. Early brain development is a period of exceptionally rapid growth and emerging function, during which disruption can have lasting consequences and contribute to neurodevelopmental disorders ^1^. Classically, neurodevelopment has been framed as a sequence of cellular programs: neurogenesis, neuronal migration and differentiation, axon guidance, and synaptogenesis^2,3^. The resulting brain scaffold then expands and reorganizes as neurons are added and connections form, while activity-dependent refinement and pruning tune existing circuits ^4,5^. Decades of research have uncovered many of the genetic, molecular, and cellular mechanisms governing these processes, establishing fundamental principles by which neurons acquire their identities and form circuits ^6^. Yet how their combined action rapidly yields a coherent functional organization across the brain remains unclear.

A central task of neurodevelopment is to assemble a connectome capable of supporting the functions required at each stage of life. The most direct view of this process comes from electron microscopy reconstructions of the *C. elegans* nervous system, which revealed extensive remodeling of synaptic wiring across development ^7^. Neuronal activity provides a complementary window: its organization bears the imprint of underlying connectivity ^8^, while directly capturing the dynamic interactions associated with emerging brain functions ^9^. Human neuroimaging has provided the broadest view of functional brain development, revealing large-scale functional networks before birth ^10,11^ and their continued reorganization through infancy, childhood, and adolescence ^12,13^. Across these periods, functional maturation is characterized by increasingly anatomically distributed interactions, the consolidation and differentiation of intrinsic networks, and selective reorganization of cross-network coupling ^13,14^. These changes unfold heterochronically, with primary sensory and motor systems maturing before higher-order association networks ^15,16,17^. Recent structural and functional brain charts have consolidated these observations into consensus trajectories of macroscale maturation ^18,14^, portraying neurodevelopment as a structured reorganization with shifting developmental priorities rather than uniform growth. Despite these conceptual advances, fMRI remains an indirect and spatially coarse measure of neuronal activity whose interpretation is complicated by age-dependent neurovascular coupling and head motion ^19,20^. How these macroscale trajectories arise from individual neurons and cellular populations, particularly during early development, therefore remains largely unknown.

Invasive studies in animal models have complemented this macroscopic view by resolving functional maturation at the level of individual neurons and circuits. In the developing mammalian cortex, neuronal activity shifts from widespread corecruitment during synchronized bursts toward sparser, desynchronized, and increasingly differentiated dynamics ^9,21,22^. At the mesoscale, wide-field imaging has revealed coherent cortical networks early in mouse postnatal life and their progressive segregation into specialized subnetworks ^23,24^. In ferrets, modular spontaneous activity emerges across cortical areas before sensory experience and subsequently undergoes area-specific refinement, while visual experience stabilizes sensory representations ^25,26,27,28^. Sparse neuronal populations can also coordinate patterned activity throughout the developing *Drosophila* brain, illustrating how cellular mechanisms can organize dynamics across much broader spatial scales ^29^. These findings broadly echo human neuroimaging by suggesting that coordinated early activity gives way to a more modular architecture with increasingly selective and reliable representations. However, functional imaging in animal models, often in rodents, remains largely restricted to accessible cortical regions, with many studies focusing on primary sensory systems, leaving associative regions and coordinated maturation across an entire brain comparatively unexplored ^30,22^.

Rapid external development, optical transparency, and genetic tractability make larval zebrafish uniquely suited to bridge developmental biology and systems neuroscience. These properties enable cellular-resolution imaging across the brain, yielding detailed accounts of brain-wide cellular dynamics underlying several behaviors ^31,32^. Despite the rapid development of zebrafish larvae, most brain-wide imaging studies have examined a single larval stage or pooled nearby ages, effectively approaching the developing brain as a fixed circuit ^32^. Developmental studies have instead charted specific systems, including retinotectal ^33,34^ and spinal locomotor circuits ^35,36^, but a large-scale account of functional maturation across the larval brain remains lacking. Because zebrafish larvae already exhibit brain-wide functional networks with organizational features shared with mammalian brains ^37,38^, their first week of life offers an exceptionally accessible window into the emergence and maturation of vertebrate brain networks. Here, we provide a comprehensive five-day structural and functional imaging survey of zebrafish brain development coinciding with the emergence and refinement of larval behavior. We identify coordinated signatures of brain maturation across anatomy, cellular activity, and systems-wide functional interactions, revealing how an early functional scaffold is progressively reorganized.

## Results

### Whole-brain calcium imaging across early neurodevelopment

We used whole-brain two-photon calcium imaging, visual stimulation, and tail monitoring to probe the functional architecture of the larval zebrafish brain across early neurodevelopment (Fig. 1a, Methods). Experiments were conducted cross-sectionally from 4 to 8 days post-fertilization (dpf) in Tg(*elavl3* :H2B-GCaMP6s) larvae expressing a pan-neuronal, nuclear-localized calcium sensor ^39^ (Fig. 1b). To contextualize this developmental window, we performed high-throughput behavioral experiments from 3 to 9 dpf, which showed the emergence of core behaviors from 4 dpf onward, including spontaneous swimming, light-driven modulation of swimming, and acoustic startle responses (Supp. Fig. S1, Methods). The magnitudes of most behavioral parameters, such as swimming levels during light or dark periods, varied approximately linearly with age, recapitulating some established patterns of locomotor maturation during the first week of zebrafish development ^40,41^. These observations reinforce 4–8 dpf as a developmental window in which a core behavioral repertoire is already present, but its expression is progressively modulated as the nervous system undergoes maturation. Existing zebrafish brain atlases are also defined at 6 dpf, providing a common anatomical reference for comparisons across ages ^42,43^.

**Figure 1:**
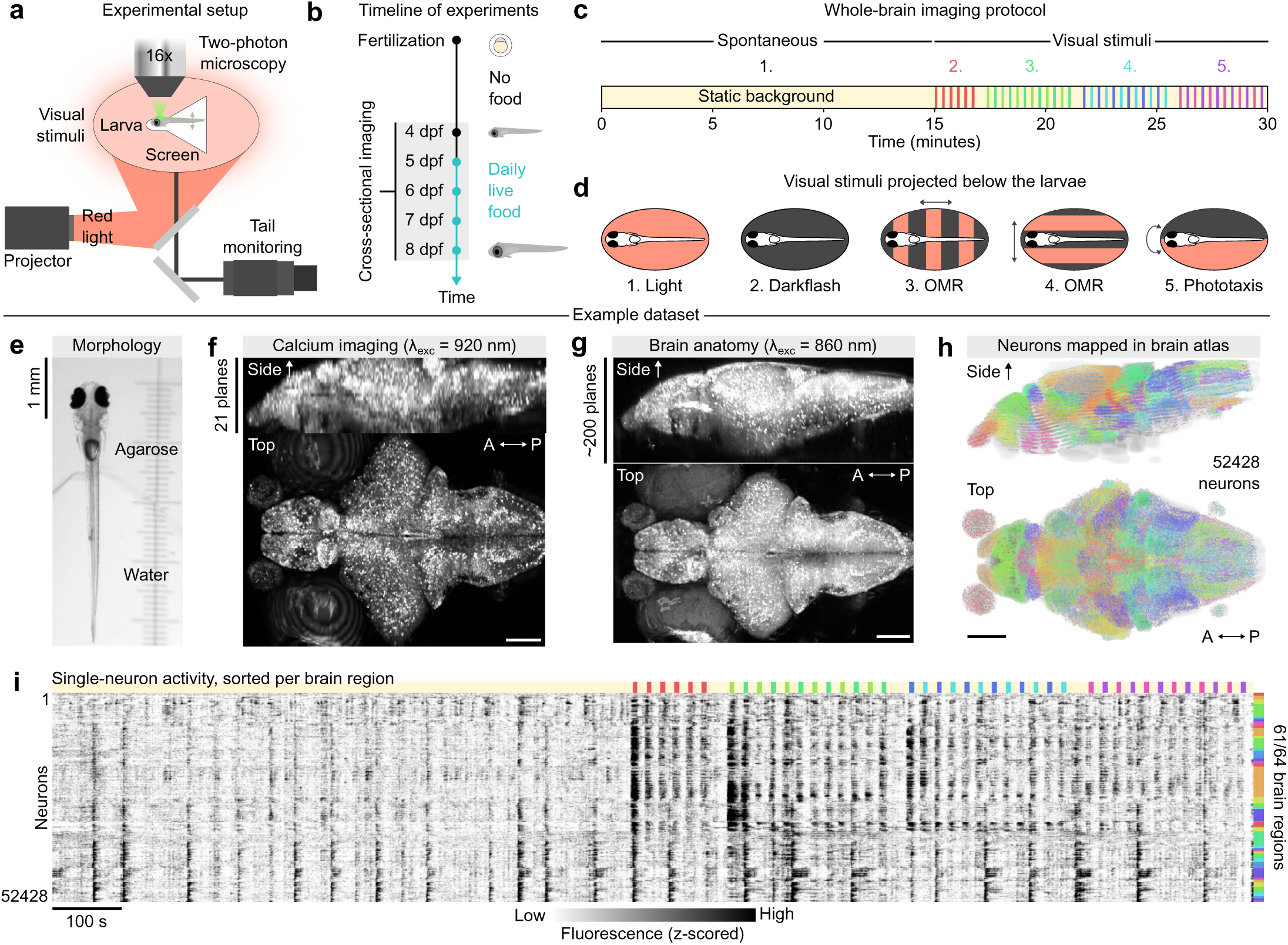
Whole-brain calcium imaging across early neurodevelopment. (**a**) Schematic of the experimental setup. (**b**) Timeline of the cross- sectional imaging experiments. (**c**) 30-minute sequence of visual stimuli used during brain-wide imaging. (**d**) Schematic of the different stimulus types. (**e**) Photograph of one representative head-restrained larva prior to brain imaging. (**f**) Maximum intensity projections of 21 calcium imaging planes, averaged over time; *λ*exc indicates the laser excitation wavelength. (**g**) Maximum intensity projections of the subsequent anatomical brain scan from the same animal as the previous panel; *λ*exc indicates the laser excitation wavelength. (**h**) Neuron centroids from one representative animal, projected into a 3D atlas coordinate system; centroids are pseudocolored by anatomical brain regions, whose masks are shown in the background. (**i**) Calcium fluorescence time series from 52,428 single neurons, sorted by brain region using the same colors as the previous panel; visual stimuli in the second half of the recordings are indicated above. A, anterior; P, posterior.

To comprehensively characterize brain function across development, we first recorded 15 minutes of spontaneous wholebrain activity under static background illumination, followed by 15 minutes of ethologically relevant visual stimuli projected below the animals ^44^ (Fig. 1c–d, Supp. Video 1; Methods). The resulting dataset comprised 79 larvae spanning five developmental stages (*n* = 13 at 4 dpf; *n* = 16 at 5 dpf; *n* = 18 at 6 dpf; *n* = 17 at 7 dpf; *n* = 15 at 8 dpf). Prior to imaging, each larva was photographed to quantify morphological features (Fig. 1e). Whole-brain calcium imaging was then performed at 1 Hz with single-cell resolution (Fig. 1f, 21 imaging planes), after which a dense anatomical scan was collected to characterize brain morphology and register each animal to the 6 dpf reference atlas (Fig. 1g, Methods). Following motion correction, nuclear segmentation, and atlas registration, neuronal activity traces were extracted and individual neurons assigned to one of 64 anatomical regions from mapZebrain ^42^ (Fig. 1h, Methods). Figure 1i shows activity from over 50,000 neurons in a representative 6 dpf subject.

### Heterogeneous brain expansion during development

Before examining brain function, we first characterized anatomical growth across development. Larvae grew significantly in length from 4.0 0.1 mm at 4 dpf to 4.7 0.1 mm at 8 dpf, corresponding to a 18.2% growth (Supp. Fig. S2a,b). Other morphological measures such as head width also grew steadily over this period and correlated positively with each other (Supp. Fig. S2b-d). To visualize brain growth, we constructed unbiased anatomical templates for each developmental stage from high-resolution structural scans. To ensure accurate morphological measurements, we corrected image distortions introduced by resonant scanning mirrors ^45^ (Supp. Fig. S3a–d; 1.28% morphological error after correction; Methods), then generated template brains using iterative multivariate ANTs registration ^46^, revealing developmental expansion across multiple brain structures (Fig. 2a; Supp. Fig. S3e–h; Supp. Video 2; Methods).

**Figure 2:**
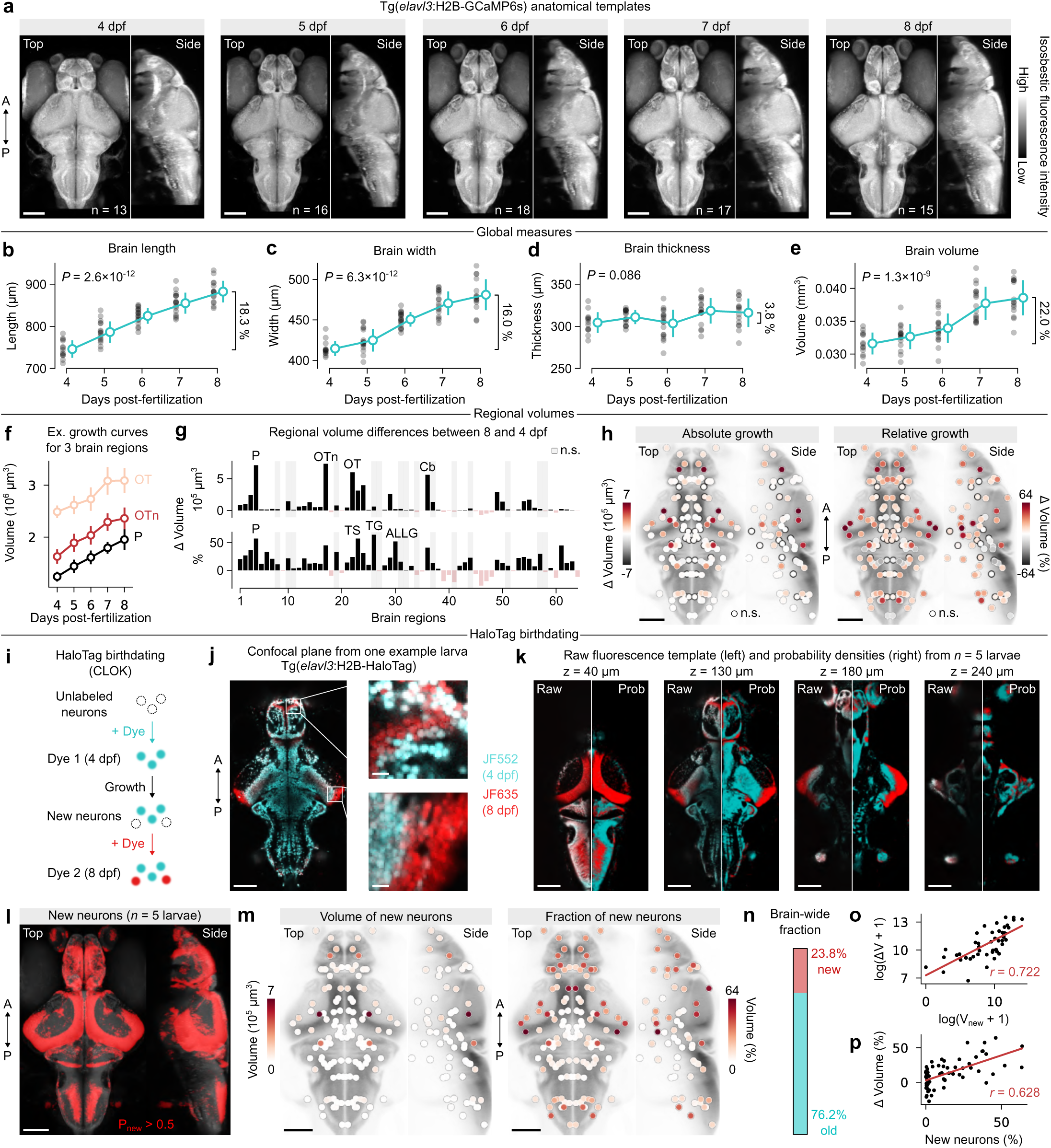
Brain anatomy changes over neurodevelopment. (**a**) Maximum intensity projections of Tg(*elavl3* :H2B-GCaMP6s) template brains at each developmental time point; the number of animals is indicated on each plot. (**b**-**e**) Global morphological measures (brain length, width, thickness, and volume); FDR-corrected *P* -values for K-W tests are indicated on each plot; average relative growth values are indicated on the right of each plot. (**f**) Example growth curves for three brain regions; error bars indicate standard deviation. (**g**) Bar charts of absolute (top) and relative (bottom) growth, measured between 4 and 8 dpf, for each brain region; shaded vertical bars indicate brain regions where no significant change is observed (Mann-Whitney U, FDR-corrected *P ≥* 0.05); key brain regions are annotated in each plot (see acronyms in Supp. Table 1). (**h**) Anatomical visualization of absolute (left) and relative (right) growth in each brain region; nodes are aligned with brain region centroids and color-coded by growth values. (**i**) Schematic of the HaloTag birthdating protocol. (**j**) Example dual-color confocal imaging plane in a Tg(*elavl3* :H2B-HaloTag) larva; blue corresponds to older neuron nuclei, while red corresponds to nuclei of newborn neurons; two small crops are shown to highlight the cellular resolution of the labeling (crop scale bars: 10 *µ*m). (**k**) Four example z-planes of the 8 dpf brain template; left halves, raw fluorescence images averaged across 5 animals; right halves, corresponding segmentation probability maps. (**l**) Spatial probability map of newborn neurons highlighted in red; probabilities are thresholded at 0.5 to accentuate regions of overlap across animals. (**m**) Anatomical visualization of absolute and fractional volumes of new neurons per brain region. (**n**) Brain-wide fraction of old and new neurons. (**o**) Linear relationship between log-transformed changes in volume of new neurons and brain region masks (Pearson *r* = 0.722). (**p**) Linear relationship between fractional volumes of new neurons and relative volume growth of each brain region (Pearson *r* = 0.628). A, anterior; P, posterior. Scale bars: 100 *µ*m.

Quantification from individual anatomical scans revealed steady growth along the anterior–posterior and medio–lateral axes over development (Supp. Video 3). Brain length increased from 746 *±* 21 to 882 *±* 26 µm (+18.3%, Fig. 2b, *P* = 2.6 *×* 10^−12^, Kruskal–Wallis (K-W) test with FDR correction), while brain width increased from 415 *±* 9 to 481 *±* 19 µm (+16.0%, Fig. 2c, FDR-corrected *P* = 6.3 *×* 10^−12^). In contrast, brain thickness showed higher variability and small but non-significant growth (+3.8%, Fig. 2d, FDR-corrected *P* = 0.086). We projected a common 3D brain mask onto individual brain scans to estimate total brain volume, observing a monotonic increase from 0.032 to 0.038 mm^3^ (+22.0%, Fig. 2e, FDR-corrected *P* = 1.3 10*^−^*^9^; Methods). Relative to the adult brain ^47^, larval brains represented only 1.08% and 1.32% of the adult brain volume at 4 and 8 dpf, respectively (Supp. Fig. S4a). Assuming a constant daily growth rate of 5.1% per day (inferred from the observed 22.0% increase over 4 days), a simple extrapolation predicts that adult brain volume would be reached in ~95 days, roughly matching the age at which the adult atlas was established ^47^ (105–112 days, Supp. Fig. S4b, Methods). Hence, this simple extrapolation yields a plausible timescale for reaching adult brain size, although the growth rate likely increases further at the juvenile stage before reaching a plateau toward adulthood ^48^.

Next, we projected 3D brain region masks from the 6 dpf atlas onto individual brain scans to quantify regional growth. Most regions exhibited gradual, monotonic growth over the 4-day period (Fig. 2f, three example regions), although some regions exhibited progressive volume decreases. Overall, 39/64 regions showed significant increases in volume between 4 and 8 dpf, whereas 10/64 exhibited significant decreases (Fig. 2g, FDR-corrected *P <* 0.05, Mann-Whitney U). In absolute volume, the pallium, optic tectum, and cerebellum contributed most to total brain growth, together accounting for ~ 47% of the overall volume increase (Fig. 2g, top). Relative growth peaked in a different subset of regions, with the torus semicircularis, trigeminal ganglion, and anterior lateral line ganglion showing the largest proportional increases (Fig. 2g, bottom). The pallium also ranked among the top regions in relative growth. Regions exhibiting volume decreases were primarily located in anterior hindbrain segments, consistent with a progressive narrowing of the hindbrain over this developmental period (Supp. Fig. S3g,h). Comparison with adult anatomy revealed that larval regional volumes are correlated with adult proportions (Supp. Fig. S4c, Pearson’s *r* = 0.739), and that the relative sizes of the 10 largest regions become increasingly similar to their adult proportions over time (Supp. Fig. S4d). These results indicate that brain growth in zebrafish is regionally heterogeneous and progressively reshapes the larval brain toward its adult proportions ^47^, with the pallium standing out in both absolute and relative expansion.

To complement our volumetric analyses, we used a recently established HaloTag birthdating approach (CLOK ^49^) to label neurons in vivo at 4 and 8 dpf using Janelia Fluor dyes in a separate cohort of Tg(*elavl3* :H2B-HaloTag) larvae (Fig. 2i), enabling brain-wide visualization of neurons born during this developmental interval (Fig. 2j). Whole-brain confocal image volumes were acquired, and labeled nuclei were segmented using a U-Net model trained on sparse manual annotations from the same dataset (Methods). Binary segmentations were then registered to the 8 dpf template brain and averaged across animals to generate continuous probability maps describing the spatial distributions of early-born and late-born (newborn) neuronal populations (Fig. 2k, Supp. Video 4). Newborn neurons integrated into many stereotyped domains, including the forebrain periventricular zone ^50^, tectal proliferation zone ^51^, and dorsal hindbrain (Fig. 2l). Proliferative zones were mostly segregated from early-born neuronal populations, with occasional mixing in transition zones. Integrating newborn probability densities within each brain region revealed that newly generated neurons occupied nearly one quarter of the 8 dpf brain volume (23.8%; Fig. 2m,n), closely matching the independently measured 22.0% increase in total brain volume (Fig. 2e). Moreover, regional growth estimates derived from birthdating correlated with atlas-based volumetric measurements (Pearson’s *r* = 0.722 and *r* = 0.628 for absolute and fractional growth, respectively; Fig. 2o,p). Notably, the pallium, optic tectum, and cerebellum exhibited both the largest volumetric expansion and the highest densities of newborn neurons.

We next performed complementary imaging experiments at 4 and 8 dpf in transgenic larvae expressing cytosolic red fluorescent proteins in either *vglut2a*-or *gad1b*-positive neurons, corresponding to glutamatergic excitatory (E) and GABAergic inhibitory (I) populations, respectively, while co-expressing nuclear GCaMP for anatomical registration (Methods). Labeled neurons were segmented in individual brains (*n* = 4 per line and developmental stage), registered to age-matched template brains, and averaged to generate whole-brain probability maps of E/I neuronal spatial distributions using the same approach as the birthdating analysis (Supp. Fig. S5a). Large-scale neurotransmitter domains were similar between 4 and 8 dpf, indicating that the brain-wide organization of E/I populations is largely established by 4 dpf (Supp. Fig. S5b, Supp. Video 5). Consistent with this observation, regional E/I volumes were highly correlated across development (Supp. Fig. S5c,d, *R*^2^ = 0.967 and *R*^2^ = 0.916 for E and I, respectively), while the global E/I ratio remained stable at slightly below 1.5:1 (Supp. Fig. S5e, *P* = 0.55, paired t-test). This estimate is lower than recent measurements at 6 dpf ^52,53^ (1.8:1), likely reflecting our use of spatial probability densities rather than direct cell counts. We finally leveraged cytosolic E/I labeling to segment neuropil (NP) and cell body (CB) compartments in 4 and 8 dpf template brains (Supp. Fig. S5f,g) before projecting these segmentations onto individual animals for quantification. Both compartments increased significantly across development, with CB volume increasing by 14.7% and NP volume by 30.6% (Supp. Fig. S5h,i). Consequently, the CB/NP volume ratio decreased from 1.55 to 1.39 ( −12.1%; Supp. Fig. S5j), indicating that neuropil expansion outpaces the volumetric contribution of neurogenesis, consistent with the progressive elaboration of dendritic and axonal processes.

Altogether, these analyses reveal widespread yet spatially heterogeneous neuroanatomical growth across the larval zebrafish brain, with region-specific growth progressively reshaping the brain toward adult proportions while large-scale excitatory–inhibitory organization remains broadly stable.

### Gradual reorganization of brain-wide functional connectivity

To characterize the brain-wide organization of neural activity across development, we next measured functional connectivity (FC), which captures functional interactions among distributed brain regions and reflects underlying anatomical connections ^38^. Individual neurons were assigned to 278 nodes (139 per hemisphere) corresponding to subdivided brain atlas regions, providing a more homogeneous spatial coverage of the brain (Supp. Fig. S6a,b, Methods). The total number of segmented neurons remained approximately constant across development, although their spatial distribution shifted toward anterior brain regions, reflecting the skewed anatomical growth observed previously (Supp. Fig. S6c,d). Average activity traces were extracted for each node, and pairwise correlations were computed to generate FC matrices for each animal. We denote FC_spont_ and FC_stim_ as the connectivity matrices measured during spontaneous and visually stimulated activity, respectively. FC matrices computed for individual stimulus epochs are shown in Supp. Fig. S6e, although subsequent analyses focus on connectivity aggregated over 15-minute recording periods.

Group-averaged FC matrices revealed broadly similar large-scale functional networks across development during both spontaneous and visually evoked activity (Fig. 3A,B). Despite their overall similarity, group-averaged FC matrices became progressively less correlated with each other with increasing developmental separation, with the lowest similarity observed between 4 and 8 dpf (Fig. 3c,d), indicating incremental changes in brain-wide functional interactions. To localize these developmental differences, we next applied network-based statistics ^54^ (NBS; Methods). Both FC_spont_ and FC_stim_ exhibited significantly larger subnetworks of strengthened and weakened connections than expected by chance (Fig. 3e, both *P <* 0.001, 1000 permutations). During spontaneous activity, increases and decreases in FC were approximately balanced (Fig. 3f; 52.6% positive changes), whereas visually evoked activity exhibited a bias toward stronger connectivity at 8 dpf (68.6% positive changes). Strengthened spontaneous connections were broadly distributed across the brain (Fig. 3g), whereas weakened connections were enriched in ventral and hindbrain regions (Fig. 3h). Mapping node-wise mean FC changes onto anatomy further revealed strengthening throughout the anterior brain and caudal hindbrain, while much of the remaining hindbrain exhibited weak or negative changes (Fig. 3i). At the regional level, the pallium showed the largest net increase in FC, whereas the intermediate hypothalamus exhibited the strongest decrease (Fig. 3j). Comparable analyses of FC_stim_ revealed both shared and distinct developmental patterns, including more pronounced connectivity strengthening in hindbrain regions (Supp. Fig. S7a-f). Intersecting spontaneous and stimulus-driven FC changes identified the pallium as the region exhibiting the greatest convergence across conditions, defined as connections undergoing significant changes of the same sign in both cases (Fig. 3k). The number of convergent edges greatly exceeded chance levels, suggesting that many developmental changes in FC persist under different sensorimotor conditions (Supp. Fig. S7h,i, *P <* 0.0001, permutation test).

**Figure 3:**
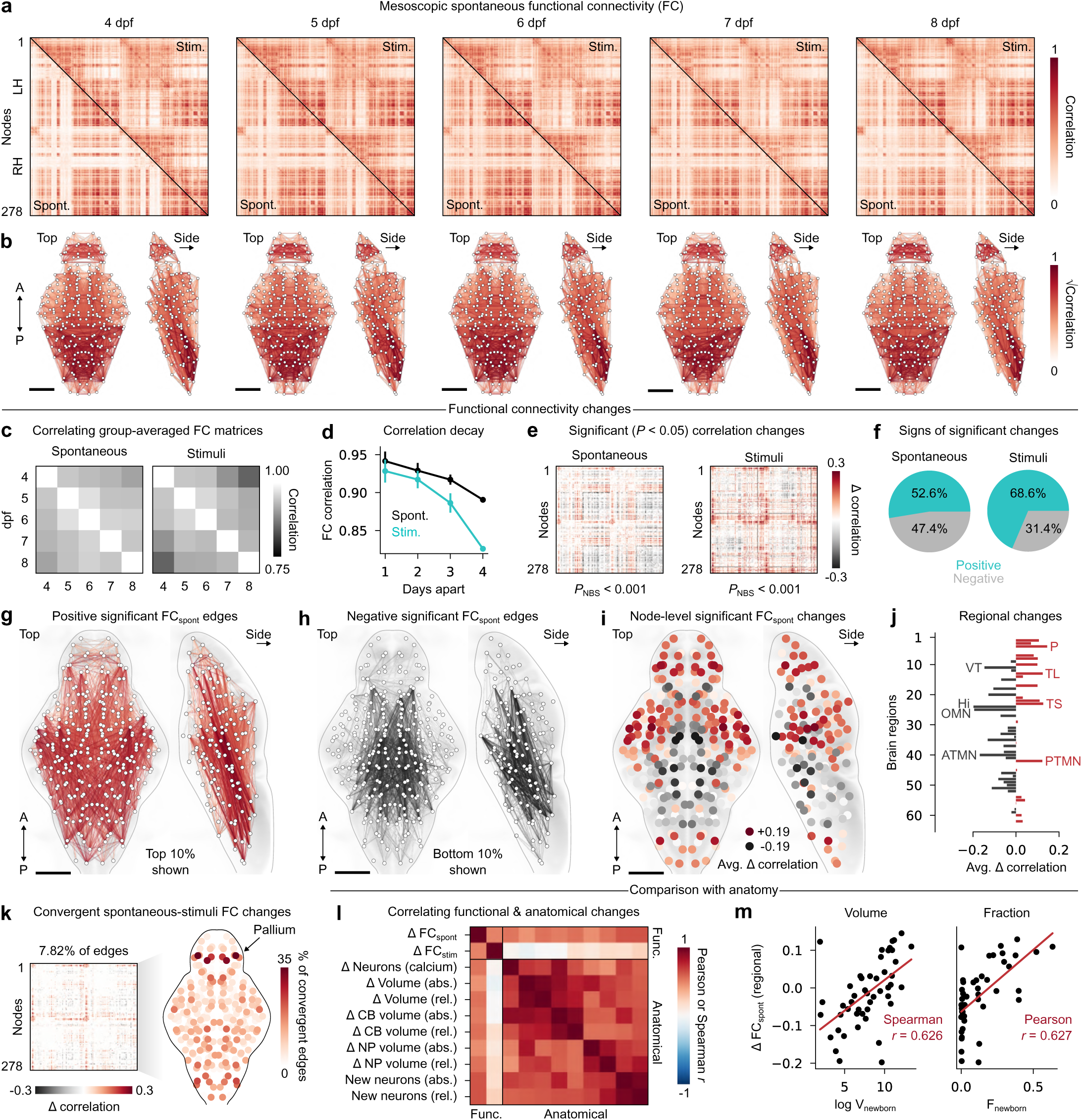
Brain-wide functional networks exhibit gradual changes. (**a**) Group-averaged FC matrices at each developmental time point; lower triangles correspond to spontaneous correlations, and upper triangles correspond to visually stimulated correlations; matrices are sorted in antero-posterior order, and from left to right hemisphere. (**b**) Network visualization of the previous FC matrices (spontaneous only); the top 10% edges are shown for each node and correlations square-rooted to aid visualization. (**c**) Correlations between group-averaged FC matrices for spontaneous and visually stimulated epochs. (**d**) Correlations averaged per time distance bin derived from the previous panel. (**e**) Significant FC changes between 4 and 8 dpf; non-significant changes are displayed in white (**f**) Signs of significant edges. (**g**) Network visualization of the top 10% of positive changes for FCspont. (**h**) Similar visualization for the bottom 10% of negative changes. (**i**) Average significant FCspont changes per network node. (**j**) Chart of average significant spontaneous FC changes per anatomical brain region. (**k**) Left, convergent FC changes across spontaneous and stimulus epochs; right, fraction of convergent edges per network node. (**l**) Correlations between region averaged FC changes and various anatomical changes measured in the previous section; the maximum value between Pearson and Spearman correlation coefficients is chosen for each comparison. (**m**) Correlations between the regional volume (left) and fraction (right) of newborn neurons and significant changes in FCspont; correlation coefficients are annotated on each plot. A, anterior; P, posterior; func., functional. Scale bars: 100 *µ*m.

The anatomical distribution of significant ΔFC_spont_ changes closely resembled the spatial distribution of newborn neurons observed in Fig. 2l, suggesting that anatomical growth supports emerging functional interactions. Consistent with this interpretation, ΔFC_spont_ correlated positively with all measured neuroanatomical changes (Fig. 3L). In contrast, ΔFC_stim_ was markedly less coupled to anatomy than ΔFC_spont_ (Fig. 3l). The strongest association was observed between ΔFC_spont_ and both the absolute and fractional volumes of newborn neurons (Spearman *r* = 0.626 and Pearson *r* = 0.627, Fig. 3m). These correlations were significant under spatial permutations which approximately preserve spatial autocorrelation (both *P <* 0.001, 1000 permutations, Methods), indicating that spontaneous FC strengthening preferentially occurs in regions undergoing high neurogenesis. Stimulus-driven FC changes instead concentrate in hindbrain regions exhibiting limited growth, potentially reflecting changes in the organization or efficacy of existing connections.

We next assessed the robustness of these observations. First, developmental FC changes could not be explained by alterations in signal quality, including noise level and signal-to-noise ratio (Supp. Fig. S8; maximum explained variance of 25.1% for noise level), indicating that they primarily reflect changes in neural activity. Second, principal component analysis (PCA) of individual FC matrices revealed multiple network components whose expression varied monotonically with age (Supp. Fig. S7j-o, spontaneous PCs 1, 3 and 4; stimulus-driven PCs 3 and 4; all FDR-corrected *P <* 0.05), further supporting a progressive brain-wide reorganization of functional connectivity on a day-to-day basis.

Together, these findings reveal widespread and gradual remodeling of brain-wide functional connectivity in zebrafish from 4 to 8 dpf. In the following sections, we show that this reorganization reflects fundamental principles of brain network organization.

### Emergence of long-range functional interactions

Functional brain networks are strongly constrained by physical distance, as nearby regions are more likely to be connected anatomically and therefore tend to exhibit correlated activity ^55^. As the brain matures, however, the elaboration and strengthening of long-range projections are expected to progressively relax this spatial constraint ^56^. We therefore asked how the relationship between FC and physical distance evolves throughout larval development.

Both spontaneous and visually evoked FC decayed with distance, consistent with the predominance of local connectivity in the larval brain ^42,38,57^ (Fig. 4a,b). This dependence was stronger during visual stimulation, reflecting enhanced local synchronization among visually responsive nodes. Across development, however, the FC-distance relationship progressively weakened for both FC_spont_ and FC_stim_ (Fig. 4c; *R*^2^ = 0.84 and 0.89, respectively), indicating that functional interactions became increasingly decoupled from spatial proximity. Consistent with this interpretation, the characteristic FC length, corresponding to the mean anatomical distance between nodes weighted by their FC strength, increased significantly from 4 to 8 dpf (Fig. 4d, both FDR-corrected *P <* 0.01, t-test). Although characteristic length was initially shorter during visual stimulation than during spontaneous activity, this difference was no longer significant by 8 dpf, suggesting that visually evoked activity increasingly recruits anatomically distributed circuits (Fig. 4d, FDR-corrected *P* = 0.077, paired t-test). Because distances were measured in a common 6 dpf atlas space, these effects are not explained by overall brain expansion.

**Figure 4:**
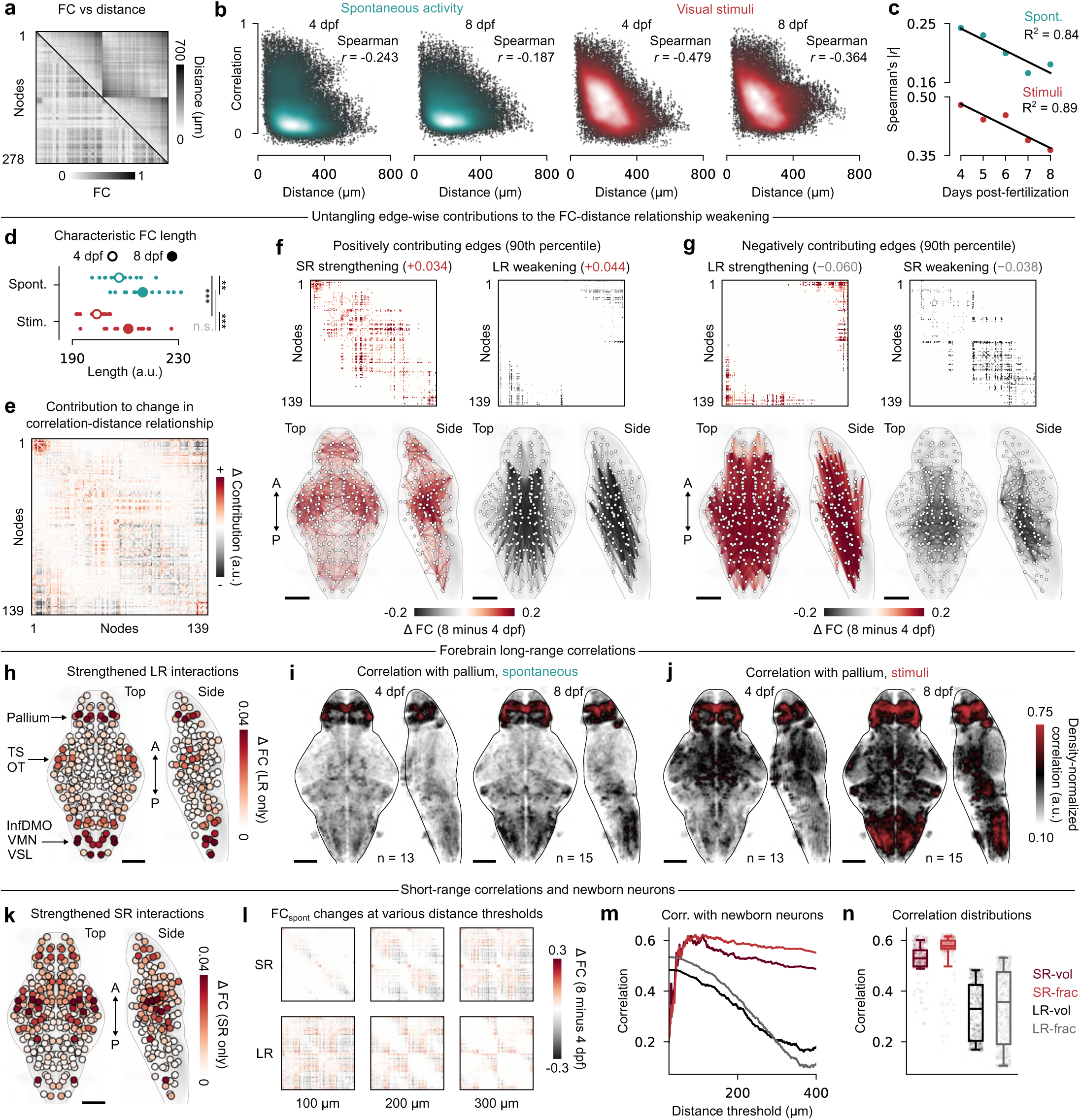
Emergence of long-range functional interactions. (**a**) Comparison between spontaneous functional connectivity (FC; lower triangle) and Euclidean distance between network nodes (upper triangle). (**b**) Example FC-distance scatter plots during spontaneous (left) and stimulus- evoked (right) activity at 4 and 8 dpf; brighter colors indicate higher point densities. (**c**) Absolute Spearman correlation between FC and distance across development for spontaneous (top) and stimulus-evoked (bottom) activity; linear regression *R*^2^ values are indicated. (**d**) Characteristic FC length measured in individual animals during spontaneous and stimulus-evoked epochs; all FDR-corrected *P* -values (unpaired or paired t-tests) are significant with the exception of the spont-stim comparison at 8 dpf. (**e**) Edge-wise contributions to the developmental change in the spontaneous FC-distance relationship (8 minus 4 dpf). Positive (negative) values denote edges that strengthen (weaken) this relationship with development. (**f**) Top 10% positive edges from **e**, separated into short-range (SR; left) and long-range (LR; right) connections; numbers in parentheses correspond to the effect magnitude. (**g**) Bottom 10% negative edges from **e**, separated into LR (left) and SR (right) connections. (**h**) Node-wise sum of FC changes associated with strengthened LR edges; see Supp. Table 1 for brain region acronyms. (**i**) Single-cell seed-based correlation maps using pallial activity during spontaneous epochs, compiled across animals at 4 (left) and 8 (right) dpf. (**j**) Same as **i**, but during stimulus-evoked activity. (**k**) Node-wise sum of FC changes associated with strengthened SR edges. (**l**) Example significant ΔFCspont matrices with different edge length thresholds. Upper row, shorter edges are preserved (SR case); bottom row, longer edges are preserved (LR case). Threshold values are indicated below. (**m**) Correlation between region-averaged FCspont changes and regional volumes or fractions of newborn neurons, at varying edge length thresholds for the SR and LR cases. Color legend is indicated on the next panel. (**n**) Compiled correlation distributions from the four previous curves. A, anterior; P, posterior; SR, short-range; LR, long-range.

To determine how this spatial decoupling emerges, we decomposed the developmental change in the FC-distance relationship into edge-wise contributions (Fig. 4e, Methods). Positive contributions corresponded to strengthened short-range (SR) or weakened long-range (LR) interactions that reinforced the spatial correlation (Fig. 4f), whereas negative contributions reflected strengthened LR and weakened SR interactions that weakened the relationship (Fig. 4g). The four edge categories contributed to the FC–distance relationship with comparable absolute magnitudes but acted in opposing directions (Fig. 4f,g). Thus, development simultaneously strengthened local circuitry and distributed interactions, but the latter effect predominated to produce an overall weakening of the FC-distance relationship. Comparable results were obtained during visually evoked activity (Supp. Fig. S9a-c).

Summing strengthened LR connections across nodes identified three major anatomical domains of emerging long-range connectivity: the pallium, the torus semicircularis together with the ventral optic tectum, and multiple caudal hindbrain regions including the inferior dorsal medulla oblongata and vagal sensory-motor nuclei (Fig. 4h). Similar domains were observed during visual stimulation, particularly in the forebrain and caudal hindbrain (Supp. Fig. S9d). To visualize these emerging interactions at cellular resolution, we performed seed-based correlation analyses using the pallium. Correlation maps revealed a striking developmental increase in brain-wide coupling with the forebrain during spontaneous activity, including several newly coupled cell clusters in hindbrain regions (Fig. 4i). Under visual stimulation, these longrange interactions became even more pronounced, particularly within caudal hindbrain regions, consistent with node-level analyses (Fig. 4j). In both conditions, increased local coupling within the pallium and neighboring subpallium further suggested the strengthening of internal forebrain connectivity.

Because newborn neurons have been shown to initially integrate circuits through predominantly local connectivity ^58,59^, we hypothesized that the association between neurogenesis and changes in spontaneous FC observed in Fig. 3m was primarily driven by SR interactions. Summing strengthened SR edges across nodes yielded an anatomical distribution that indeed resembled the previously observed maps (Fig. 4k). To avoid relying on a single arbitrary distinction between short- and long-range connections, we calculated significant regional FC changes across a range of distance-thresholded matrices, excluding edges longer than a given threshold for the SR case or shorter than that threshold for the LR case (Fig. 4l). We then correlated these thresholded FC changes with both the regional volume and fraction of newborn neurons, observing substantially stronger correlations for SR interactions (Fig. 4m,n). For SR matrices, correlations peaked at thresholds of 81 and 112 *µ*m for volume and fraction, suggesting that strengthened functional interactions within a radius of approximately 100 *µ*m correlate most strongly with neurogenesis (Fig. 4m). By contrast, LR matrices were weakly correlated, particularly at thresholds of 200–400 *µ*m, where only the longest-range interactions were retained (Fig. 4m). Thus, the association between FC_spont_ and newborn neurons is driven predominantly by strengthened local correlations.

Finally, to determine whether spatial decoupling through LR strengthening reflected motor-related activity, we regressed motor signals from all node time series before recomputing FC (Methods). Although this substantially reduced brain-wide correlations, the developmental weakening of the FC-distance relationship persisted (Supp. Fig. S9e,f, *R*^2^ = 0.702), and multiple long-range forebrain-hindbrain interactions remained positive (Supp. Fig. S9g,h), suggesting that these changes extend beyond motor-related activity.

Together, these findings show that larval brain development progressively relaxes the spatial constraints on functional connectivity through strengthening of long-range interactions, promoting increasingly distributed and integrated brain networks. In parallel, short-range correlations strengthen preferentially within regions undergoing high neurogenesis, suggesting that local circuit refinement proceeds alongside the maturation of long-range pathways.

### Developmental increase of functional network modularity

A defining feature of brain networks is their organization into mesoscale modules, whereby groups of brain regions exhibit preferential interactions with one another ^60^. As neural circuits mature and become functionally specialized, both anatomical wiring ^7,61^ and functional interactions ^62,24^ are expected to progressively segregate into mesoscale modules.

To characterize the modular architecture of larval FC, we used a hierarchical consensus modularity approach previously applied to zebrafish ^38^ (Methods). Although raw correlations exhibit significant modular organization ^38^, we used global signal regression (GSR), following previous zebrafish network analyses ^37^, to enhance the block structure of FC matrices by removing widespread signals that span multiple modules and obscure functional boundaries (Supp. Fig. S10a,b). Because the interpretation of negative edge weights following regression can be ambiguous, we focused on strictly positive correlations, although the following conclusions remain robust when using signed modularity (Supp. Fig. S10c).

To identify connectivity modules across spatial scales, we generated consensus node partitions from group-averaged FC matrices by using the Louvain community detection algorithm while varying its resolution parameter across 1000 runs ^63^ (*γ* = 0.25–10; Methods). We then hierarchically clustered node coassignment probabilities computed across all generated partitions, yielding nested consensus node partitions spanning 2 to 100 modules (Fig. 5a, examples shown at 8 dpf). To estimate uncertainty, we performed a leave-one-out analysis in which each animal was sequentially excluded from its age group, generating 79 group-average FC matrices. We benchmarked empirical modules against 79 geometric surrogate networks generated from spatially decaying exponential correlation profiles matched to the data (Fig. 5b, Methods). These surrogates reproduced the broad structure of empirical FC (Supp. Fig. S10d,e, average *r* = 0.480), particularly within single hemispheres (Supp. Fig. S10f, average *r* = 0.585), providing a benchmark for modular organization expected from spatial embedding alone ^64^.

**Figure 5:**
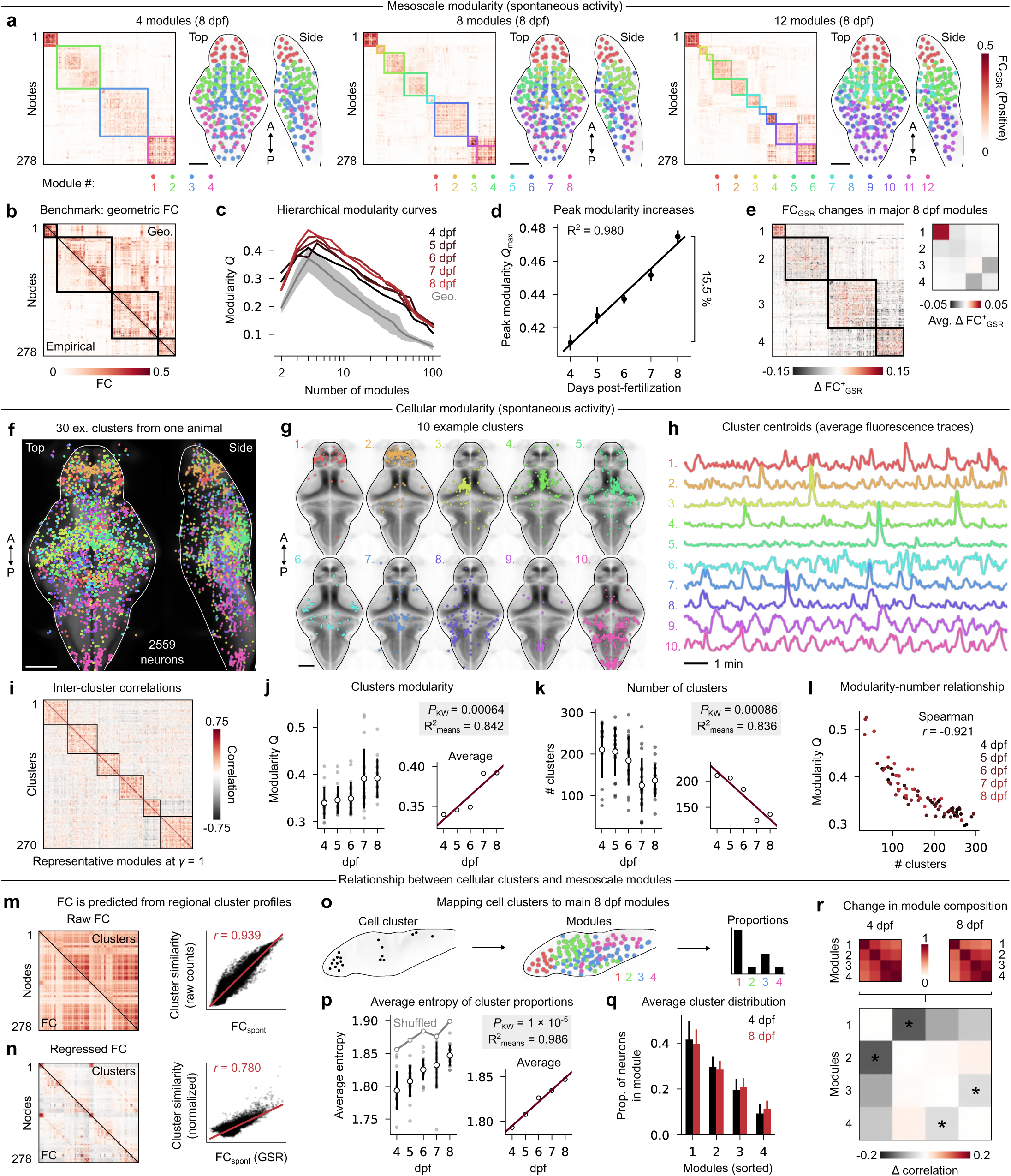
Developmental increase of modularity in mesoscale and cellular functional networks. (**a**) Group-averaged 8 dpf FC matrix (left) sorted by connectivity modules (right); 4, 8 and 12 modules are displayed. (**b**) Comparison between 4 dpf FC (bottom left) and an exemplar geometric surrogate FC matrix (top right). (**c**) Average hierarchical modularity curves for each age group, compared with modularity of geometric surrogates (gray line); the shaded region indicates minimum and maximum values. (**d**) Linear regression of peak modularity values for each developmental time point; error bars indicate standard deviation estimated using the leave-one-out procedure. (**e**) Left, subtraction between 8 dpf and 4 dpf FC^+^ matrices, sorted with 8 dpf modules; right, average within each block of the previous matrix representing within- and between- module connectivity changes. (**f**) Centroids of neurons from 30 different pseudocolored functional clusters, overlaid on brain anatomy. (**g**) 10 example clusters, individually separated from the previous panel. (**h**) Average fluorescence traces of the 10 previous clusters. (**i**) Representative inter-cluster correlation matrix, sorted by detected connectivity modules. (**j**) Left, distributions of average inter- cluster modularity per age group, with error bars indicating standard deviation; right, linear regression of distribution averages; *P* -value for K-W test and linear regression coefficient indicated above. (**k**) Similar plot for the average number of detected clusters per animal. (**l**) Relationship between the number of clusters and inter-cluster functional modularity; a Spearman correlation coefficient is indicated on the plot. (**m**) Left, comparison between FC (bottom left) and cluster profile similarity, averaged over all 79 animals; right, correlation between the two matrix triangles. (**n**) Similar plot comparing FC after global signal regression with normalized cluster profiles. (**o**) Schematization of the cluster mapping in mesoscale modules analysis. (**p**) Similar plot to **j**,**k** displaying the average Shannon entropy of cluster distributions onto the four 8 dpf modules for each animal; a gray curve indicates the average entropy measured from randomly shuffled cluster labels. (**q**) Average cluster distribution profile for 4 and 8 dpf age groups, with modules sorted for each cluster in order of most to least populated. (**r**) Top, correlations of average cluster similarity profiles between the four 8 dpf modules for 4 and 8 dpf animals; bottom, difference between the two above matrices, with asterisks denoting significant correlation differences (FDR-corrected *P <* 0.05, Mann-Whitney U test). A, anterior; P, posterior. Scale bars: 100 *µ*m.

Across all hierarchical levels, empirical networks exhibited consistently higher modularity than geometric surrogates (Fig. 5c), indicating that functional subdivisions cannot be explained solely by distance-dependent connectivity. Despite appearing anatomically compact, empirical modules were significantly more spatially distributed than those generated by geometric FC (Supp. Fig. S10g), further indicating that functional modules are organized beyond simple nearest-neighbor relationships. Peak modularity increased progressively over development, rising linearly from 4 to 8 dpf (Fig. 5d; *R*^2^ = 0.980, 15.5% increase). Notably, 4 dpf networks exhibited peak modularity values comparable to those of geometric surrogates, suggesting that early functional subdivisions are only marginally more modular than those expected from simple distance-based constraints.

We repeated the same analysis for stimulus-evoked FC. Although stimulus-evoked networks consistently exceeded geometric null expectations (Supp. Fig. S10i), their modularity increased only weakly across development (*R*^2^ = 0.214). By comparison, spontaneous networks were initially less modular, but progressively reached similar modularity levels by 8 dpf (Supp. Fig. S10j). Moreover, stimulus-evoked modules exhibited spatial extents comparable to spontaneous modules (Supp. Fig. S10k), yet the overlap between spontaneous and stimulus-evoked partitions remained modest and peaked only deeper in the hierarchy at 8 modules (Supp. Fig. S10l), indicating that visual stimulation recruits a distinct modular organization that undergoes comparatively little developmental refinement.

We next asked whether spontaneous functional modules reorganize spatially throughout development. Peak modularity consistently occurred near 4 modules, suggesting that the number of major functional subdivisions remains stable across ages (Fig. 5c). Nevertheless, module composition gradually evolved over development. Hierarchical partition overlap progressively decreased with increasing separation between age groups (Supp. Fig. S10m,n, Pearson *r* = *−*0.557), indicating gradual changes in module composition. Consistent with this observation, changes in node coassignment probabilities between 4 and 8 dpf identified several regions with shifting module allegiance, including the cerebellum and torus longitudinalis (Supp. Fig. S10o). For example, cerebellar nodes were preferentially associated with tectal modules at 4 dpf but became incorporated into hindbrain modules by 8 dpf (Supp. Fig. S10p). Despite these refinements, module overlap remained non-negligible across age groups and peaked at intermediate hierarchical levels around 10–15 modules (Supp. Fig. S10q), indicating that developmental correspondence emerges deeper in the modular hierarchy.

Lastly, we used the four principal modules identified at 8 dpf, corresponding roughly to forebrain, midbrain, and hindbrain subdivisions, as a developmental reference to determine how connectivity changes contribute to the observed increase in modularity. In principle, higher modularity may arise from stronger connectivity within modules, weaker connectivity between modules, or both. Within-module strengthening was dominated by the forebrain, whereas the remaining modules exhibited mixed positive and negative FC changes that largely canceled out (Fig. 5e). In contrast, between-module connectivity generally weakened, particularly between anterior (modules 1 and 2; forebrain-midbrain) and posterior (modules 3 and 4; dorsal-ventral hindbrain) subdivisions. Thus, increasing modularity arises through both the strengthening of internal forebrain interactions and the progressive segregation of anterior and posterior brain systems. Despite their net cancellation, within-module changes were larger in absolute magnitude than between-module changes, indicating that developmental remodeling preferentially occurs within pre-existing modular boundaries (Fig. 5e).

Together, these findings show that spontaneous functional networks become progressively more modular during larval development, reflecting the functional specialization and segregation of large-scale brain subnetworks.

### Reorganization of neuronal ensembles within mesoscale module boundaries

A major unresolved question is how mesoscale functional modules emerge from the activity of individual neurons and how developmental changes in cellular activity patterns give rise to large-scale network organization ^65^.

To examine the cellular basis of increasing modularity, we identified functional neuronal ensembles ^66^, defined as groups of neurons sharing similar activity patterns regardless of their anatomical location. We adapted an iterative functional clustering approach in which the number of clusters is not specified a priori, but emerges through the successive merging of functionally similar neuronal populations ^67^ (Methods). Five randomly initialized clustering runs were performed for each animal during spontaneous activity, yielding reproducible partitions across runs (Supp. Fig. S11a-c). The resulting clusters collectively spanned the brain (Fig. 5f), although many formed anatomically compact, lateralized or bilaterally symmetric populations (Fig. 5g). Neuronal clusters captured several recurring activity motifs, including highly active forebrain populations, sparsely bursting tectal ensembles, hindbrain clusters dominated by large-amplitude motor events, and slowly oscillating hindbrain populations (Fig. 5h).

Because these ensembles were defined functionally rather than anatomically, inter-cluster correlations naturally segregated into positively and negatively correlated modules (Fig. 5i; example partition at *γ* = 1). The modularity of inter-cluster correlations, averaged across clustering runs, increased progressively from 4 to 8 dpf, recapitulating the developmental trajectory observed in anatomically defined networks 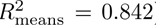. In contrast, the number of detected clusters declined from 211 *±* 68 at 4 dpf to 136 *±* 41 at 8 dpf (Fig. 5k; *P* = 0.00086, K-W test, 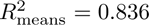. Network modularity was strongly negatively correlated with the number of clusters (Fig. 5l; Spearman *r* = *−*0.921), while mean cluster size increased with age (Supp. Fig. S11d-h). These trends suggest that small, weakly correlated ensembles progressively merge into larger and more coherent functional units to promote modularity. The number of neurons excluded by the clustering algorithm also increased from 2086 2068 to 4428 3009, reflecting a growing population of neurons with noisy, presumably reduced activity (Supp. Fig. S11i,j).

We next examined how these cellular ensembles relate to mesoscale network organization. Brain-wide FC was largely recapitulated by correlating regional cluster profiles—vectors describing the number of neurons per cluster within each node—rather than neuronal activity itself (Fig. 5m; Pearson *r* = 0.939). Similar results were obtained for FC_GSR_ after normalizing cluster profiles to reduce the disproportionate influence of large clusters (Fig. 5n; Pearson *r* = 0.780). These findings indicate that regional functional connectivity is largely determined by the relative composition of neuronal ensembles within each node. To understand how ensemble composition shapes mesoscale modules, we mapped each cluster onto the four principal modules identified at 8 dpf (Fig. 5o). The average entropy of cluster distributions increased linearly with age 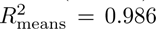, indicating more even cluster distributions across modules.

Consistent with this observation, ranked module occupancy profiles progressively flattened from 4 to 8 dpf, with average occupancy in the two dominant modules decreasing in favour of the third and fourth modules (Fig. 5q). Shuffled cluster labels produced a similar developmental trend, indicating that this can largely be explained by increasing cluster sizes and more homogeneous distributions of segmented neurons over the four modules at later ages (Fig. 5p). At first glance, this result appears incompatible with the concurrent increase in network modularity, which might instead predict that functional ensembles become increasingly confined within individual modules. However, we found that pairwise correlations between the average cluster profiles of each module decreased significantly over development, particularly between modules 1 and 2 and between modules 3 and 4, mirroring the module segregation observed earlier (Fig. 5r; Mann–Whitney tests, FDR-corrected *P <* 0.05). Thus, although neuronal ensembles become increasingly distributed across the brain, the mesoscale modules they populate become progressively more functionally differentiated. This observation suggests that increasing long-range interactions and increasing segregation in anatomically compact modules are not opposing processes, but rather complementary features of network maturation.

Together, these findings suggest that developmental increases in mesoscale modularity arise through the consolidation of neuronal activity into larger functional units and the progressive differentiation of their representation across brain modules. This compositional reorganization provides a cellular basis for the emergence of increasingly segregated largescale functional networks.

### Shift of visual responses along a retinal depth gradient

Visual processing undergoes extensive maturation during early development, especially in primary visual structures ^33^, yet how these changes are distributed across other brain circuits remains unclear. To map visual responses across the brain, we identified stimulus-responsive neurons (SRNs) showing a significant increase in fluorescence during the 5 s following stimulus onset relative to pre-stimulus activity (Methods). Significance was assessed separately for each neuron and each of the eight stimulus categories using randomized stimulus-onset times (500 permutations per neuron; FDR-corrected *P <* 0.05). Across all animals, we identified 543,442 SRNs distributed mostly throughout the anterior brain, with the notable exception of olfactory regions (Fig. 6a,b, Supp. Video 6). The inferior olive also stood out as a hindbrain region with prominent visual responses. Example recordings show robust stimulus-locked calcium transients (Fig. 6c,d).

**Figure 6:**
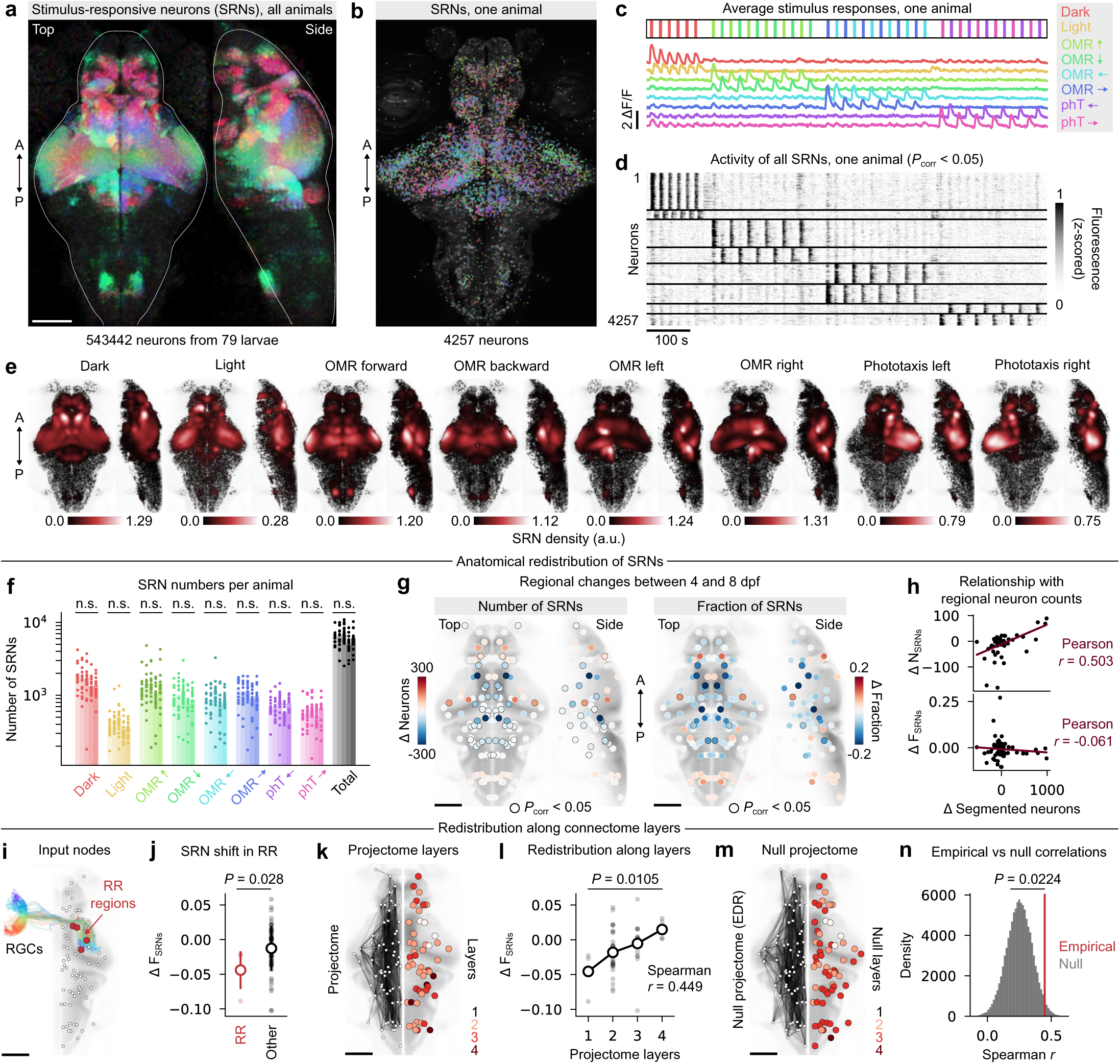
Shift of visual responses along a retinal depth gradient. (**a**) Spatial density of SRNs combining all 79 animals; pseudocolors represent different stimuli and are consistent with other plots (see annotation on the right of the figure). (**b**) SRNs from one representative animal overlaid on its brain projection. (**c**) Overlaid average traces of SRNs for each of the 8 stimulus categories, from the same animal. (**d**) Raster plot of all SRNs, sorted per stimulus category in the same order as **c**. (**e**) Combined SRN neuron centroids colored by local cell density for each stimulus category; density units are normalized to the average highest density of each stimulus for comparison. (**f**) Number of SRNs per animal per stimulus category, with consecutive bars reflecting different time points; no significant difference is observed after FDR correction, except inter-stimulus differences. (**g**) Anatomical heatmaps of variation in the number (left) or fraction (right) of SRNs per brain region between 4 and 8 dpf; black outlines indicate statistical significance (*P <* 0.05, Mann-Whitney U tests, FDR-corrected). (**h**) Correlation between the values from the previous plots and the variation in the total number of segmented neurons per brain region. (**i**) Morphologies of 54 RGCs from mapZebrain, mirrored to the left eye, with main retinorecipient (RR) brain regions annotated on the right. (**j**) Regional changes in fraction of SRNs between 4 and 8 dpf in RR and other brain regions; *P* = 0.028, permutation test, see Methods. (**k**) Left, directed projectome edges between brain regions; right, discrete projectome layers, with 1 indicating source RR regions, and 4 indicating regions that are 3 edges away from source nodes. (**l**) Regional changes in fraction of SRNs between 4 and 8 dpf, sorted per projectome layer; Spearman correlation coefficients is indicated on the plot; *P* = 0.0105 for distribution changes, K-W test. (**m**) Similar to **k**, with projectome edges randomized according to an EDR model, yielding null projectome layers on the right. (**n**) Null distribution of Spearman correlations between Δ*F*_SRN_ and null projectome layers (100,000 surrogates); a vertical red line indicates the empirical correlation, which significantly exceeds the null distribution (empirical *P* = 0.0224). A, anterior; P, posterior. Scale bars: 100 *µ*m.

Pooling SRNs across ages revealed stimulus-specific response maps that recapitulated established visuomotor pathways in zebrafish ^68,67^ (Fig. 6e). Responses to changes in illumination were enriched in the left habenula, consistent with its known functional asymmetry ^69^, whereas dark flashes evoked widespread bilateral activity. Phototactic stimuli predominantly recruited the hemisphere contralateral to the stimulated eye. Forward and backward optomotor response (OMR) stimuli produced similar bilateral patterns, although forward motion recruited more neurons overall, whereas directional OMR stimuli elicited strongly lateralized responses in the pretectum, optic tectum and anterior hindbrain. Beyond these dominant response domains, sparse SRNs were detected throughout the brain, including in the dorsal hindbrain. Occasional responses in regions without an established visual role, such as the olfactory epithelium, likely represent the expected background of false-positive detections.

Remarkably, the total number of SRNs remained stable from 4 to 8 dpf, with no stimulus category showing a significant developmental change (Fig. 6f; all FDR-corrected *P >* 0.05). We therefore asked whether this numerically stable population was redistributed across brain regions. When responses were pooled across stimuli, several regions exhibited significant changes in either SRN number or fraction between 4 and 8 dpf (Fig. 6g, 23*/*64 and 7*/*64 regions with FDR- corrected *P <* 0.05 for number and fraction, Mann-Whitney U test). The optic tectum, pallium, and torus semicircularis showed the largest increases in SRN number. These regions also underwent substantial anatomical growth, and regional changes in SRN number were positively correlated with changes in the total number of segmented neurons, indicating a potential sampling bias (Pearson *r* = 0.503; Fig. 6h, top). Despite this number increase, the fraction of visually responsive neurons within these regions decreased, and overall, regional changes in SRN fraction correlated weakly to changes in the number of segmented neurons (Pearson *r* = −0.061; Fig. 6h, bottom). Several diencephalic regions, including posterior tuberculum, dorsal and ventral thalamus, exhibited large decreases in both SRN number and fraction, whereas caudal hindbrain regions showed convergent increases. Thus, visual responsiveness is redistributed across brain regions during development, increasing in some regions while declining in others. Because absolute SRN numbers are influenced by region size, we focused our next analyses on SRN fractions calculated from pooled neuronal counts within each age group.

The reduction of diencephalic and mesencephalic visual responses raised the possibility that visual activity progressively shifts from primary retinorecipient structures toward downstream circuits. To test this hypothesis, we used retinal ganglion cell (RGC) morphologies from mapZebrain ^42^ to identify the principal retinorecipient (RR) regions. High counts of RGC terminals identified the tectal neuropil and periventricular layer, dorsal thalamus and pretectum as principal RR domains, consistent with established visual anatomy ^70^ (Fig. 6i). Developmental changes in SRN fraction were significantly lower in RR regions than elsewhere in the brain (Fig. 6j, *P* = 0.028, permutation test, Methods), supporting a relative redistribution away from the initial targets of retinal input. We next asked whether this redistribution followed the anatomical progression of visual pathways. We constructed a directed mesoscale projectome from 4,327 single-neuron morphologies ^42^, and assigned to each region a projectome-depth index corresponding to its shortest directed path from an RR node (Fig. 6k). Developmental changes in SRN fraction increased steadily and significantly with discrete projectome layers (Fig. 6l; Spearman *r* = 0.449; *P* = 0.0105, K-W test), indicating a gradual shift toward regions positioned further downstream from retinal input. To determine whether this gradient specifically followed the organization of the empirical projectome, we generated surrogate networks from an empirically matched exponential distance rule ^71,57^ while preserving network density (Fig. 6m; Methods). The gradient in SRN fraction was stronger than expected from the spatially constrained null model (Fig. 6n; *P* = 0.0224, 100,000 networks), suggesting that the gradient is shaped by the specific organization of the empirical projectome. Comparable gradients were recovered across other age comparisons, although weaker in adjacent time points, indicating that the effect is not specific to the 4-versus-8-dpf comparison (Supp. Fig. S12a-h).

To complement the cell-count-based analyses, we quantified visual responsiveness directly from regional activity. We generated regressors approximating the calcium responses evoked by each visual stimulus (Supp. Fig. S12i) and used multilinear regression to predict the activity of each of the 278 mesoscale nodes in every animal (Supp. Fig. S12j; Methods). Averaged across hemispheres, regression coefficients increased significantly with age in 22*/*139 brain regions and decreased in 17*/*139 regions, revealing gradual and bidirectional changes in visually evoked activity (Supp. Fig. S12k; FDR-corrected *P <* 0.05; Methods). Consistent with the count-based results, the variance explained by visual stimuli decreased moderately across the diencephalon and mesencephalon but increased markedly in the caudal hindbrain, where visually evoked activity was more prominent at 8 dpf, particularly during OMR stimuli (Supp. Fig. S12j,k).

Finally, we examined whether an analogous redistribution emerges through the optimization of artificial neural networks. We trained convolutional neural networks (CNNs) to classify MNIST digits ^72^ using a simplified, biologically inspired architecture comprising a single convolutional layer, representing retinal processing, four fully connected layers representing successive processing stages, and a global activity regularization mimicking metabolic constraints (Supp. Fig. S12l; Methods). Between early and late training, activation progressively shifted from the input toward deeper layers, producing a gradient similar to the one observed along the zebrafish projectome (Supp. Fig. S12m,n). Early networks already performed well above chance (48.7 ± 6.5% accuracy compared with a 10% chance level), indicating that this shift reflects network optimization rather than trivial random initialization.

Together, these findings reveal a developmental redistribution of visual responses from primary retinorecipient structures toward downstream brain regions. The emergence of a qualitatively similar shift during artificial network optimization suggests that this reorganization reflects a general computational principle of visual maturation.

### Global reduction of motor-related activity

To determine whether developmental changes extended from sensory to motor systems, we next examined the maturation of motor behavior and its neural correlates. Using camera recordings acquired during brain imaging, we tracked tail movements and detected swimming events with a temporal convolutional network ^73^ trained on manually annotated traces (Fig. 7a,b; Methods). Across 79 animals, we identified 6,409 swimming events. Swim-bout frequency varied considerably across individuals but declined moderately with age, indicating reduced locomotor output in older larvae (Supp. Fig. S13a, *R*^2^ = 0.125). Bout amplitude, velocity and duration showed similar weakly decreasing and variable developmental trends (Supp. Fig. S13b-f). Swimming was slightly more frequent during the first 15 minutes of imaging, and locomotor output remained strongly correlated between the two halves of the experiment, revealing stable individual differences in motor drive (Supp. Fig. S13g-i; Spearman *r* = 0.864). Relative to a 5-minute pre-stimulus baseline, only dark flashes significantly increased swim-bout frequency during stimulus presentation (Supp. Fig. S13j; FDR-corrected *P <* 0.05), potentially reflecting rapid habituation or suppression of motor responses to the subsequent stimuli under head-restrained conditions. Overall, motor output decreased moderately across development but remained highly variable among individuals.

**Figure 7:**
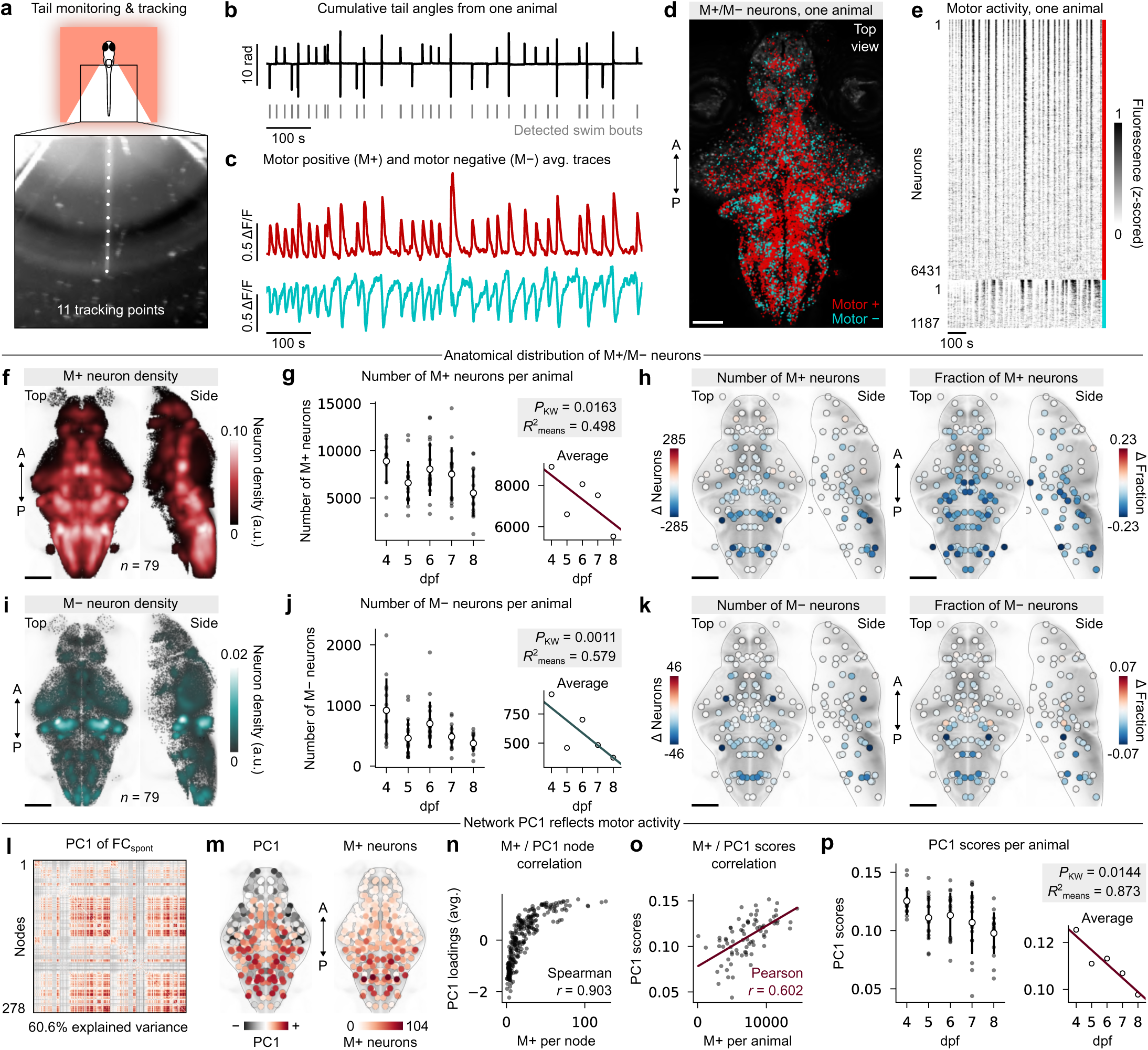
Global reduction of motor-related activity. (**a**) Example behavioral camera frame and tail tracking points. (**b**) Cumulative tail angles from one representative animal; upward and downward deflections correspond to left and right swimming, respectively. (**c**) Average fluorescence traces of M+ and M*−* neuronal populations associated with the trace above. (**d**) Anatomical visualization of the two previous neuronal populations. (**e**) Raster plot of the two previous populations, sorted by *P* -value to accentuate the most strongly-correlated neurons. (**f**) M+ neuron centroids from all animals in atlas coordinates, colored by local cell density. **g** Left, distributions of M+ neuron counts per age group, with error bars indicating standard deviation; right, linear regression of distribution averages; *P* -value for K-W test and linear regression coefficient are indicated above. (**h**) Anatomical heatmaps of variation in the number (left) or fraction (right) of M+ neurons per brain region between 4 and 8 dpf; black outlines indicate statistical significance (*P <* 0.05, Mann-Whitney U tests, FDR-corrected). (**i**-**k**) Similar to **f** -**h**, but for M*−* neurons. (**l**) First principal component of FCspont, with arbitrary positive and negative values matched with the following plot; the component’s % of explained variance is indicated below. (**m**) Left, node-averaged PC1 loadings; right, average number of M+ neurons per node. (**n**) Spearman correlation between the two previous quantities; correlation coefficient is indicated on the plot. (**o**) Correlation between the number of M+ neurons and PC1 scores per animal; correlation coefficient is indicated on the plot. (**p**) Similar to **g** and **j** but for individual PC1 scores per age group.

We next asked whether brain-wide motor representations remained stable in the absence of overt behavioral changes. Using the same event-triggered fluorescence approach applied to visual responses, we identified neurons exhibiting significant increases (motor-positive, M+) or decreases (motor-negative, M-) in activity during swimming (Fig. 7c-e; FDR-corrected *P <* 0.05, see Methods for the animal 29 exception due to decreased swimming). M+ neurons were broadly distributed throughout the brain and particularly abundant in the hindbrain, but were comparatively sparse in the optic tectum (Fig. 7f). Their numbers declined substantially with age, from 8897 *±* 2432 neurons at 4 dpf to 5525 *±* 2490 neurons at 8 dpf 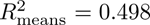. This reduction extended across most brain regions and was especially pronounced in the hindbrain (Fig. 7h). M-neurons were less abundant and formed discrete populations in the hindbrain, cerebellum, optic tectum, habenula and pretectum (Fig. 7i). Their numbers declined as well, from 919 *±* 527 neurons at 4 dpf to 344 *±* 373 neurons at 8 dpf 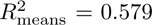, with most regions showing reductions in both their number and fraction (Fig. 7k). Importantly, the abundance of M+ and M-neurons was only weakly associated with swimming frequency, amplitude, or other behavioral measures (Supp. Fig. S13k; maximum explained variance, 20.2%). Thus, both positively and negatively motor-related populations become progressively less prominent during larval development, beyond what can be explained by differences in measured locomotor output.

Because neuron counts do not directly capture how motor-related activity contributes to brain-wide dynamics, we next examined its expression in spontaneous FC. PCA of individual FC matrices identified a dominant component accounting for 60.6% of inter-individual variance (Fig. 7l). Its node-averaged loadings accentuated the hindbrain and closely matched the spatial distribution of M+ neurons (Fig. 7m,n; Spearman *r* = 0.903). Moreover, individual PC1 scores, which quantify the component’s expression in each animal, correlated positively with M+ neuron abundance (Fig. 7o; Pearson *r* = 0.602). The dominant axis of FC variation therefore captures a widespread motor-related connectivity signature. Consistent with the cellular decrease, PC1 scores decreased significantly with age and followed a less variable developmental trajectory than M+ or M*−* neuron counts (Fig. 7p; *P* = 0.0144, K-W test; 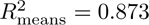). This reduction was largely independent of differences in locomotor output (Supp. Fig. S13k).

Together, these findings reveal a global weakening of motor-related activity throughout the brain as the larval nervous system matures, in contrast to the complex redistribution of visual representations observed over the same developmental

### Intrinsic forebrain fluctuations recruit an expanding neuronal population

Despite widespread maturation throughout the brain, several of our analyses converged on the forebrain as a major locus of functional reorganization. The teleost forebrain is of particular interest because it comprises pallial and subpallial territories broadly homologous to mammalian cortical and subcortical systems and plays a central role in higher-order behavioral control ^74,75,76,77^.

To identify the dynamics underlying this reorganization, we applied spatial independent component analysis ^78^ (ICA) to coarse-grained activity signals from all 79 animals. ICA decomposes brain activity into statistically independent spatial patterns and their associated time courses (Fig. 8a). Across parameter choices, ICA consistently recovered a component encompassing the forebrain, whose activity fluctuated spontaneously throughout each recording (Fig. 8b,c; 10 components; Supp. Fig. S14a). Other components captured anatomically compact domains, including unilateral and bilateral motor and tectal circuits (Fig. 8b,c). To assess their robustness across animals, we repeated ICA in individual recordings and correlated the resulting components to the group-level solution. The forebrain component emerged as the most reproducible feature of brain activity, exhibiting the highest correspondence in 63/79 animals during spontaneous activity and 66/79 during visual stimulation (Fig. 8d; both FDR-corrected *P <* 0.0001, paired *t*-test). Visual stimulation increased the reproducibility of several other components, consistent with stimulus-locked activity, but none surpassed the forebrain component (Fig. 8d; Supp. Fig. S14b,c).

**Figure 8:**
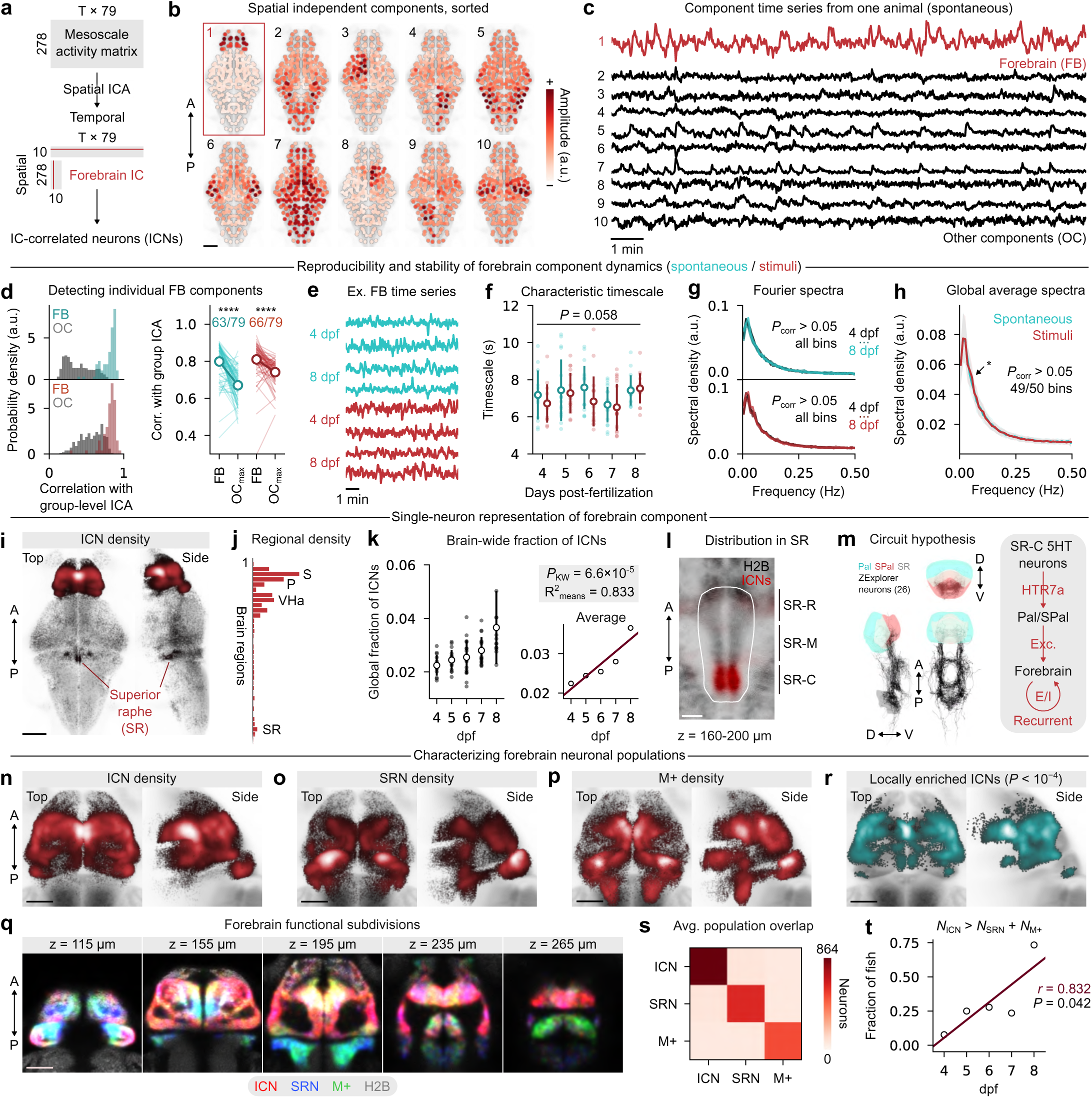
Intrinsic forebrain fluctuations recruit an expanding neuronal population (**a**) Schematization of the section’s main analysis workflow. (**b**) 10 group-level spatial components uncovered through spatial ICA; the forebrain (FB) component is accentuated in red. (**c**) Time courses of the 10 previous components; the FB signal is accentuated in red. (**d**) Left, distributions of correlation between individual and group-level components; colored distributions (blue, spontaneous; red, stimuli) correspond to FB correlations, while gray corresponds to other components (OC); right, comparison between FB and the highest OC correlation; FB correlates significantly more with group-level components (FDR- corrected *P* = 10*^−^*^13^ and *P* = 10*^−^*^9^ for spontaneous and stimuli, paired t-test); fractions of animals where FB is the highest correlation are indicated above. (**e**) Example FB component time series from two animals per age group and per recording period. (**f**) Distributions of FB signal characteristic timescales per age group and recording period; no significant difference is observed (*P* = 0.058, K-W test). (**g**) Average Fourier power spectra of the FB time series for each age group during spontaneous and stimulus epochs; no significant distribution shifts are observed across all spectral bins (*P >* 0.05, K-W tests with FDR-correction). (**h**) Average spectra for the spontaneous and stimuli periods; only one spectral bin differs significantly in amplitude, indicated by an arrow (FDR-corrected *P* = 0.048, Wilcoxon signed-rank test). (**i**) ICN centroids from all animals in atlas coordinates, colored by local cell density; the superior raphe (SR) is annotated. (**j**) Average density values per brain region, with key brain regions annotated; the plot is truncated due to negligible densities in hindbrain regions. (**k**) Left, distributions of global fractions of ICNs per age group, with error bars indicating standard deviation; right, linear regression of distribution averages; *P* -value for K-W test and linear regression coefficient are indicated above. (**l**) Filtered spatial density of ICNs (red) overlaid on the H2B template (gray) in the SR region; white outlines correspond to the mapZebrain region mask, and the region’s three main subdivisions are roughly annotated on the right; scale bar, 20 *µ*m. (**m**) Left, anatomical visualization of ZExplorer neurons with somata located in SR and axon terminals located in the pallium or subpallium regions; right, minimal SR-forebrain circuit hypothesis; image generated using the ZExplorer website’s rendering interface. (**n**-**p**) ICN, SRN, and M+ neuron centroids colored by local cell density within the forebrain in atlas coordinates. (**q**) Filtered spatial densities from the three previous populations overlaid on anatomy at various depths within the forebrain; colors are indicated below. (**r**) ICN centroids, colored by local density, that exceed a local enrichment threshold (*P <* 0.0001), accentuating different subregions of the forebrain. (**s**) Average cellular overlap between the three functional populations (that is, the number of cells belonging to two categories simultaneously); diagonal values correspond to the average population sizes. (**t**) Fraction of animals per age group whose number of ICNs exceeds the two other populations combined; Pearson correlation coefficient and permutation test *P* -value are indicated on the plot. A, anterior; P, posterior. Scale bars: 100 *µ*m for whole-brain, 50*µ*m for forebrain.

The forebrain component exhibited remarkably stable temporal dynamics across development and experimental conditions (Fig. 8e). Its integrated autocorrelation time averaged 7.11 ± 1.24 seconds and showed no significant developmental trend or difference between spontaneous and visually stimulated epochs (Fig. 8f; *P* = 0.058, Kruskal–Wallis test, Methods). Similarly, Fourier spectra were largely similar across ages and recording conditions. No frequency bin differed significantly between age groups (Fig. 8g; all FDR-corrected *P >* 0.05, K-W tests), whereas 1*/*50 frequency bins differed between spontaneous and visually stimulated periods (Fig. 8h; FDR-corrected *P* = 0.048, Wilcoxon signed-rank test). The small increase in spectral power at 0.05 Hz likely reflects residual stimulus-locked activity (20-s interstimulus interval), as spatial ICA does not enforce temporal independence between components. Thus, the most reproducible component of larval zebrafish brain activity is an intrinsic forebrain fluctuation whose temporal properties remain largely invariant across development and are minimally affected by ongoing visual input.

We next asked whether this component contributed to the developmental increase of functional connectivity. Regressing its time course from each node before computing FC markedly reduced the number of convergent developmental changes in FC_spont_ and FC_stim_ between 4 and 8 dpf (Supp. Fig. S14d) and nearly abolished the increase in connectivity within the forebrain module (Supp. Fig. S14e). The recovered component therefore accounts for a substantial fraction of emerging forebrain functional connectivity. Because its temporal structure remains stable, this increase likely involves the progressive recruitment of additional neurons into a shared dynamical process.

To identify the cellular substrate of this process, we correlated the activity of every neuron with the forebrain component time course and assessed significance using temporal permutations (Methods; 400 permutations per neuron; FDR-corrected *P <* 0.05). Significant neurons, hereafter termed independent-component neurons (ICNs), were distributed throughout much of the forebrain, with their highest density in the subpallium, and were also unexpectedly present in the superior (dorsal) raphe and two bilateral clusters in the neighboring medulla (Fig. 8i,j). The fraction of ICNs increased steadily with age (Fig. 8k; *P* = 6.6 10*^−^*^5^, K-W test; *R*^2^ = 0.833 across group averages), providing a simple cellular explanation for increasing within-forebrain connectivity. Inspection of their 3D spatial distribution further revealed their localization within the caudal part of the SR (Fig. 8l), spatially consistent with a recently identified serotonergic subpopulation of caudal raphe neurons (SR-C neurons ^79^). Examination of the ZExplorer atlas ^52^ further identified superior raphe neurons with dense projections to the forebrain, terminating predominantly in the subpallium (Fig. 8m, left). These observations suggest a candidate circuit in which caudal serotonergic raphe neurons modulate the forebrain fluctuation through excitatory projections to pallial or subpallial circuits, from which activity may propagate to other forebrain regions ^80^ (Fig. 8m).

The heterogeneous distribution of ICNs further suggested that they occupy specialized domains within the forebrain. We therefore compared their spatial organization with that of the sensory-responsive (SRNs) and motor-correlated (M+) neurons identified previously (Fig. 8n–p). Although the three populations occupied partially overlapping domains (Fig. 8q), nearest-neighbor enrichment analysis identified ICN-dense regions relative to both sensory and motor populations (Fig. 8r; FDR-corrected *P <* 0.0001, one-sided Gaussian-tail tests based on 100 label permutations; Methods). These regions included the hippocampal-like dorsolateral pallium ^81,82^ and the medial subpallium, pointing to a specialized anatomical substrate for intrinsic forebrain dynamics. Moreover, fewer than 10% of ICNs were also sensory- or motor-correlated at the single-cell level, indicating that they constitute a largely distinct functional class (Fig. 8s). As this population expanded, it progressively outnumbered the sensory and motor populations, exceeding their combined numbers in 11/15 larvae at 8 dpf (Fig. 8t; Pearson *r* = 0.832, *P* = 0.042, permutation test).

Together, these findings identify a neuronal population underlying the emergence of intrinsic forebrain functional connectivity and nominate ascending serotonergic inputs as a candidate driving mechanism.

## Discussion

Early development requires the coordinated growth and functional reorganization of the brain to support an increasing behavioral repertoire. Here, by integrating brain-wide anatomical and functional measurements, we identify coordinated cellular and network-level signatures of vertebrate brain maturation over the first days of life.

Neuroanatomical growth of the larval zebrafish brain is highly heterogeneous. Several forebrain and midbrain structures expand disproportionately, whereas hindbrain regions show comparatively limited and, in some cases, negative anatomical growth. Regions with the highest neurogenesis exhibit the largest increases in spontaneous correlations, mostly at short distances. This association may reflect several mechanisms, such as the greater functional similarities between clonally related neurons ^83^ or neurons born within the same time period ^84,85^, or the presence of instructive, developmentally transient activity patterns within these regions ^9^. Although newborn neurons are initially less active than mature neurons, our measurements integrate activity across several days and likely capture many neurons after their functional integration and maturation ^58,49^. Despite reduced growth, hindbrain structures also undergo notable functional changes, including increases in long-range correlations, sensory responsiveness and stimulus-driven functional connectivity. This indicates that substantial circuit maturation also occurs within pre-existing neuronal populations. Indeed, the preferential expansion of neuropil relative to cell body regions suggests that early larval brain maturation primarily favors the elaboration of dendritic and axonal arbors, most likely accompanied by synaptic remodeling. Combining birthdating ^49^ with developmental synaptic measurements ^86^ will help establish how the contributions of neurogenesis and neuritic growth evolve throughout development.

Brain-wide functional connectivity undergoes extensive reorganization, with distinct subnetworks strengthening or weakening across development. A prominent feature of network reorganization is the emergence of long-range interactions, particularly in forebrain and caudal hindbrain regions. Given the established relationship between structural and functional connectivity in zebrafish ^87,38^ and other model organisms ^8,88^, these changes likely reflect the progressive establishment and strengthening of long-range axonal projections. Axons extend over several days in zebrafish ^89^ and mice ^90^, while forming synaptic contacts before reaching their final targets, providing a straightforward mechanism for the spatial expansion of correlated activity. This relationship may also be bidirectional: emerging projections allow activity to propagate across distant regions, and the resulting patterns of coordinated activity can in turn regulate axonal growth and circuit organization ^91,92^. Concurrent with long-range strengthening, functional networks become increasingly modular, in line with observations from structural and functional connectivity across species ^61,7,24^. Because modules are anatomically compact, strengthening long-range interactions across their boundaries might be expected to undermine modular segregation. However, cellular analyses reconcile these seemingly opposing trends: neuronal ensembles become more broadly distributed across modules, while individual modules become increasingly differentiated by the particular mixtures of functional ensembles they contain. Thus, the expansion and broader anatomical distribution of neuronal ensembles can support functional specialization by generating increasingly distinct cellular compositions across modules. This suggests a simple mechanism through which long-range integration and modular segregation can emerge concurrently ^93,94^. Because gene expression partially predicts single-neuron ^95^ and region-level ^38^ functional interactions in zebrafish, an important next step will be to determine how transient transcriptional programs and genetically defined cell populations contribute to the establishment of functional ensembles and large-scale functional organization.

Visual processing extends beyond primary sensory structures and relies on coordinated activity across anatomically distributed regions. Although previous zebrafish studies have characterized maturation of the retinotectal pathway and visual tuning in tectal neurons ^33,96,58,51^, less is known about how visual representations reorganize throughout an entire developing brain. We observed a striking redistribution of visually responsive neurons, with responsiveness decreasing in several retinorecipient regions while moderately increasing in forebrain and hindbrain structures. This reorganization followed the architecture of retinal projection pathways, suggesting that visual activity is progressively redistributed along pre-established circuits. These findings parallel the increased developmental recruitment of higher-order visual and association cortices observed in humans, suggesting that maturation progressively extends stimulus representations along the visual hierarchy ^97,98,99^. Comparable developmental expansions of visually driven activity have also been observed across vertebrate animal models ^100,101,102^. In the C. elegans connectome, the proportion of synapses contributing to feedforward communication from sensory to motor neurons also increases with age ^7^. Together, our findings place these cross-species observations within a brain-wide connectomic framework, revealing a graded developmental redistribution of visual responsiveness from direct retinal targets through successive layers of the connectome. We observed a similar redistribution in biologically inspired CNNs trained under metabolic constraints, suggesting that this organizational principle may arise from general computational pressures favoring the transformation of visual inputs into increasingly abstract or behaviorally relevant representations. Consistent with this idea, recent work in zebrafish has shown that sensory representations outside primary visual regions, particularly in anterior hindbrain rhombomeres, show enhanced alignment with the statistics of behavioral outputs, a feature also reproduced in feedforward CNNs ^103^. The developmental increase in visual responsiveness within caudal regions may therefore reflect the progressive transformation of sensory information into motor-decodable signals to generate appropriate actions. Of note, our immobilized larvae exhibited reduced stimulus-evoked movements, potentially because agarose embedding and passive stimulus presentation strongly gate motor output ^104^. Future studies using closed-loop virtual reality in head-restrained animals ^105^ or tracking microscopy in freely swimming larvae ^106^ will be required to determine how the maturation of visual processing relates to real-time changes in behavioral output.

In parallel with the redistribution of sensory representations, brain-wide motor-related activity progressively declined. The precise identity of these declining motor-correlated populations remains uncertain under our current paradigm. A subset of motor-positive neurons likely comprises spinal-projecting neurons (SPNs) that directly drive locomotion ^107^, but many others may represent premotor interneurons, neurons receiving efference copies or sensory feedback from self-generated movements, and, in the case of negative correlations, inhibitory feedback and circuits involved in movement preparation or suppression ^104,108^. These functional categories can be more readily distinguished in millisecond-timescale electrophysiological or voltage-imaging recordings, where activation timing relative to movement onset can separate the early recruitment of putative SPNs or premotor neurons from late activation consistent with efference copies or sensory feedback ^109,110^. By contrast, our slow calcium recordings blur these temporal offsets into apparent coactivation. We speculate that this decline reflects the refinement of broadly distributed reafferent and efference-copy signals into sparser, more selective motor representations ^111^. It is unlikely to involve a comparable loss of core locomotor neurons, as descending motor circuits develop through the sequential addition of neuronal populations: early-born circuits supporting rapid, vigorous movements are retained, while later-born populations enable slower and more refined locomotion ^112,113^. Development may therefore expand core locomotor circuits while progressively restricting the propagation of motor-related signals throughout the brain. Despite the global decrease observed here, widespread motor representations remain a defining feature of the mature mammalian brain, permeating much of the neocortex and strongly shaping both spontaneous and task-related activity ^114,115,116^. Declining motor-related activity is therefore unlikely to continue indefinitely and may instead represent a transient rebalancing of brain-wide activity during early development, presumably through synaptic pruning mechanisms or increased inhibitory signaling.

Among all brain regions, the forebrain emerges as a major locus of developmental reorganization. As the evolutionary precursor of numerous mammalian cortical and subcortical structures ^74,75^, the teleost telencephalon supports diverse cognitive functions that mature throughout development^76,77^. Perhaps reflecting increasing demands, the forebrain undergoes pronounced anatomical expansion, establishes stronger long-range interactions with the rest of the brain, and exhibits the largest increase in intrinsic functional connectivity. We also identify a distributed population of forebrain neurons displaying prominent spontaneous fluctuations that are largely independent of sensory and motor activity. This population expands disproportionately relative to the total forebrain neuronal population and becomes a dominant source of spontaneous activity, suggesting that intrinsic dynamics are an important signature of the developing telencephalon. Despite previous studies that have also noted reproducible patterns of high spontaneous activity within the larval forebrain ^31,117^, the mechanisms underlying these activity fluctuations remain unclear. Their association with dorsal raphe neurons, together with the extensive serotonergic innervation and modulation of the teleost forebrain ^79,118^, raises the possibility that they are driven by ascending serotonergic inputs, although establishing causality will require targeted perturbations. One explanation is that these fluctuations reflect elevated vigilance induced by experimental manipulation, as serotonergic raphe inputs were recently shown to drive large-amplitude synchronous activity in the adult zebrafish pallium during vigilance states ^119^. However, this evoked activity required a strong olfactory cue and was confined to the central pallium, suggesting that the neuronal population described here expresses a distinct form of spontaneous dynamics. An intriguing possibility is that this activity constitutes an evolutionary precursor of the mammalian default mode network (DMN), a distributed system characterized by coherent spontaneous activity during rest ^120,121^. Although direct homologies between the teleost and mammalian forebrain remain unresolved, this interpretation is broadly consistent with the population’s early emergence ^122^, its ubiquitous spontaneous activity, and its enrichment within the dorsolateral pallium, which shares genetic, anatomical and functional features with the mammalian hippocampal formation ^123,74,75^, a major component of the DMN’s medial temporal subsystem ^124^. The preferential involvement of caudal raphe neurons, which are minimally recruited during sensorimotor behaviors but respond to food-related signals ^79^, further raises the possibility that these fluctuations reflect ongoing internal or homeostatic states. Whether this activity represents a proto-DMN or, more conservatively, a developmentally transient pattern of activity, analogous to the coherent spontaneous activity observed in the immature mammalian hippocampus ^125^, remains unresolved. Regardless, its expansion indicates that spontaneous forebrain dynamics, potentially driven by serotonergic inputs, are an important feature of early forebrain maturation.

Our study presents several limitations that should be considered. First, calcium imaging provides an indirect measure of neuronal activity and is susceptible to developmental changes in optical signal quality as the brain grows. Although signal properties explained only a moderate fraction of the observed changes, residual imaging biases cannot be excluded, and developmental variation in fluorescent sensor expression may further influence measurements over extended timescales. Brain-wide electrophysiological surveys circumvent some of these limitations but remain experimentally demanding, and current high-density probes are poorly suited to the larval zebrafish brain ^116^. Emerging voltage-imaging approaches offer a promising middle ground by providing a more direct measure of neuronal activity while retaining brain-wide optical access ^126,110^. Second, several findings imply changes in synaptic connectivity that cannot be resolved directly by functional imaging. The strengthening of long-range interactions, increasing network modularity, and redistribution of sensory representations all suggest substantial remodeling of structural circuits. Combining functional imaging with developmental connectomics ^7,127^, made increasingly feasible by automated electron-microscopy reconstruction pipelines ^53^, will provide a critical opportunity to link synaptic remodeling directly to the emergence of large-scale brain dynamics. Third, our study covers a narrow developmental window from 4 to 8 days post-fertilization. Although many behavioral, anatomical, and functional changes followed approximately linear trajectories during this period, neurodevelopment does not proceed linearly across the lifespan. Studies in other vertebrates have identified periods of rapid behavioral or circuit reorganization interspersed with more stable phases ^18,14,128^, suggesting that many trends described here will eventually plateau or undergo abrupt transitions. Non-monotonic trajectories may also be observed within individual brain structures, such as changes in spontaneous burst properties within the optic tectum ^34^, reflecting localized maturation processes that differ from global trends. Extending these analyses into juvenile and adult stages remains technically challenging because the zebrafish brain rapidly outgrows the capabilities of conventional microscopes. The smaller, transparent teleost Danionella cerebrum is a promising model for investigating network maturation and emerging brain functions over substantially longer timescales ^129,130^. Finally, we examined stereotypical development under controlled laboratory conditions. Experience, sensory environment, and other components of the exposome are major influences on brain development. Applying this framework across different environmental conditions will help disentangle genetically programmed developmental trajectories from environment-dependent circuit refinement or alteration ^131^.

## Methods

### Animal care

All experiments were conducted on zebrafish larvae aged 3–9 days post-fertilization (dpf). Larvae were maintained in embryo medium (13.7 mM NaCl, 0.54 mM KCl, 1.0 mM MgSO_4_, 1.3 mM CaCl_2_, 0.025 mM Na_2_HPO_4_, 0.044 mM KH_2_PO_4_, and 4.2 mM NaHCO_3_; pH 7.2, adjusted with 12 N HCl) at a density of one larva per 2 mL in 50 mL Petri dishes. Dishes were maintained in an incubator at 28*^◦^*C under a 14:10 h light/dark cycle. From 4 dpf onward, the medium was replaced daily and larvae were fed live *Tetrahymena thermophila* CU428.2 (Cornell University). *T. thermophila* were cultured at room temperature in Petri dishes containing sterile SPP medium (2% proteose peptone, 0.1% yeast extract, 0.2% glucose, 33*µ*M FeCl_3_, 250 *µ*g/mL penicillin/streptomycin, and 0.25 *µ*g/mL amphotericin B) and harvested during the stationary phase. They were washed three times by centrifugation (2 min at 0.8g per wash), with the pellet resuspended in embryo medium after each centrifugation. To promote consistent growth across clutches, food delivery was standardized by calibrating the optical density of *T. thermophila* suspensions using a spectrophotometer. The administered volume was adjusted to yield approximately 15,000 *T. thermophila* per larva, corresponding to a final concentration of 7,500 cells/mL in the housing medium. All procedures were approved by the Université Laval Animal Care Committee (CPAUL protocol #2022-1012).

### Behavioral experiments and analysis

Zebrafish larvae aged 3–9 dpf from two lines, Casper and Tg(*elavl3* :H2B-GCaMP6s) (also in Casper background), were individually placed in 96-well plates containing approximately 0.3 ml of embryo medium per well. Plates were secured in a ZebraBox monitoring system (ViewPoint Behavior Technology, France), and locomotor activity was recorded daily during a 1-h multisensory stimulation protocol at 30 fps. Larvae were first exposed to 30 min of white illumination delivered from beneath the plate (4,700 lux), followed by five acoustic startle stimuli (400 Hz vibration, 100 ms duration, 30 s interstimulus interval) under white illumination. The larvae then underwent three 5-min dark periods interleaved with two 5-min light periods. Finally, five additional acoustic startle stimuli with the same parameters were delivered in darkness. Locomotion was quantified in 1 s bins using the ViewPoint tracking algorithm. Tracked displacement was converted into arbitrary locomotor units and averaged within the relevant stimulus epochs to obtain the behavioral measures shown in Supp. Fig. S1. Light-period locomotion was quantified during the initial 30-min illumination period, whereas dark-period locomotion was averaged across the three dark blocks, excluding transitions. Responses to light-to-dark and dark-to-light transitions were quantified over the 5 s following each transition, and acoustic startle responses were quantified from the 1-s bin containing stimulus delivery. Response probabilities were calculated by binarizing locomotion based on the zero values naturally produced by the tracking software and averaging across repeated trials within each animal (three light-to-dark transitions, two dark-to-light transitions, and five acoustic stimuli under each illumination condition) before population averaging. Linear regressions excluded the 3 dpf time point because locomotor activity at this age was markedly lower than all subsequent ages. Data from the two lines were pooled as both exhibited equivalent developmental trends. After each recording session, larvae were returned to standard housing and feeding conditions before undergoing the same experiment on the following day. The Casper population was divided into two groups that were monitored on alternating days, such that each animal was tested every 2 days, to confirm that daily testing did not influence development.

### Two-photon imaging

Larvae were embedded in 2% low-melting-point agarose (Invitrogen, 16520100) in 30-mm glass-bottom Petri dishes (Mat-Tek, P50G-1.5-30-F) and oriented upright for brain imaging using a toothbrush bristle, without direct contact with the animal. Agarose was carefully removed from below the swim bladder and around the tail using a scalpel (Fisher, 35-205), allowing the tail to move freely. The preparation was then submerged in embryo medium and transferred to the microscope for imaging. Calcium imaging was performed using a Scientifica SliceScope resonant-scanning two-photon microscope controlled with SciScan LabVIEW software and equipped with a Spectra-Physics Insight X3 tunable laser. Fluorescence was collected through a piezo-driven 16 × water-dipping objective (Nikon, 0.8 NA), reflected by a T585lpxr dichroic mirror, filtered using a 505/119-nm bandpass filter, and detected with a GaAsP photomultiplier tube. Whole-brain calcium imaging was performed at an excitation wavelength of 920 nm and an average power of 25 mW, measured after the objective. Brain activity was recorded across 22 imaging planes (512 × 720 pixels; zoom, 1.375×; volume rate, ~ 0.986 Hz) separated by approximately 12 *µ*m. The first plane was excluded because of piezo recoil artifacts associated with the sawtooth scanning pattern. To define the imaging volume consistently across animals, the uppermost plane was positioned at the dorsal surface of the brain, and the lowermost plane was positioned slightly above the ventral boundary of the pan-neuronal expression pattern. The galvo voltage controlling the vertical scanning amplitude was slightly increased in older larvae to accommodate their greater brain length. Although this adjustment slightly compressed images along the vertical axis, cellular resolution was preserved and remained comparable across developmental stages (Supp. Fig. S8a–e). Most animals underwent a single continuous 30-min recording. In some animals, axial drift occurred during the first half of the experiment but stabilized completely during the second half. For these animals, an additional 15-min recording was acquired to replace the missing spontaneous activity segment. Any experiments where notable z-drift was observed (e.g. cells moving out of focus) were immediately discarded.

High-resolution anatomical stacks were acquired at 860 nm, near the isosbestic point of GCaMP6s, using the following parameters: 512 720 pixels, 200 planes, 2-*µ*m axial spacing, 32× frame averaging, and 1.25× zoom. Before each imaging session, fluorescent beads were imaged at 11 positions distributed across the horizontal field of view to characterize spatial distortions introduced by resonant scanning. These measurements were subsequently used to correct the anatomical stacks (see Brain unwarping). A subset of experiments, including morphological validation and imaging of the Tg(*vglut2a*:DsRed)^nns14Tg^and TgBAC(*gad1b*:DsRed)^nns26Tg^ reporter lines, was performed using a similar Scientifica SliceScope two-photon microscope equipped with a galvo–galvo scanning configuration. To image excitatory and inhibitory neuronal populations, the *vglut2a* and *gad1b* reporter lines were crossed with Tg(*elavl3* :H2B-GCaMP6s) animals. Green and red fluorescence were subsequently filtered using 514/44 nm (FF01-514/44-25, Semrock) and 650/150 nm (FF01- 650/150-25, Semrock) bandpass filters, respectively, and detected simultaneously using two GaAsP photomultiplier tubes The green channel, corresponding to pan-neuronal nuclear GCaMP6s expression, was used for brain registration.

### Visual stimulation

Visual stimuli were projected underneath the imaging chamber using a red-light DLP projector (AAXA Technologies, KP-750-00) equipped with a Kodak Wratten 2 dark-red #29 filter and directed toward the animal using a dichroic mirror (Thorlabs). Standard white printing paper secured against the glass bottom of the imaging chamber served as a diffuse projection screen. The screen spanned the larva’s entire visual field, except for a small cut-out region for tail monitoring. Before each experiment, a calibration cross video was used to center the projected frames relative to the microscope objective and the larva’s head, ensuring consistent spatial positioning of the stimuli, particularly for phototaxis. Tail movements were monitored under infrared illumination to avoid interference with visual stimulation. During the first 15 min of brain imaging, larvae were maintained under constant, spatially uniform illumination. This was followed by a 15-min visual-stimulation period. Dark-flash stimuli consisted of 7-s periods of darkness followed by 13 s of uniform illumination, repeated six times. Optomotor response (OMR) stimuli consisted of binary gratings with a bar width of 0.25 cm moving at 0.5 cm/s in each of four orthogonal directions. Forward and backward motion were interleaved within one block, and leftward and rightward motion within a second block, using the same 7-s stimulation and 13-s interstimulus timing. Phototaxis stimuli were generated by projecting dark 1.5 mm to either the left or right of the larva, with the two sides presented alternately. Each stimulus type was presented six times. The ’light’ stimulus was not programmed as a separate stimulus but occurred naturally during the intervals following dark flashes. Stimuli were rendered as .avi videos at 30 frames/s using Python. Video playback was triggered by microscope acquisition using an Arduino and a custom-written Bonsai workflow ^132^.

### Behavioral monitoring and tail tracking

Tail movements were recorded from below the imaging chamber at approximately 400 frames/s using a high-speed camera (FLIR Blackfly S USB3, BFS-U3-28S5M-C) equipped with an adjustable macro lens (Navitar Zoom 7000). The larva was illuminated from above at an angle of approximately 45*^◦^* using 850-nm LED arrays. Infrared light passed through a dichroic mirror before being reflected toward the camera and isolated using a bandpass filter (Thorlabs; center wavelength, 852 nm; bandwidth, 10 nm). A 4× binning factor was applied to increase the signal-to-noise ratio, yielding 160 × 160-pixel images with sufficient spatial resolution for accurate tail tracking. Camera acquisition was triggered by microscope scanning using an Arduino and a custom-written Bonsai workflow ^132^.

### Calcium imaging data processing

Raw 16-bit microscopy data were separated into 21 .tif imaging planes. Lateral motion was corrected independently within each plane using the piecewise non-rigid registration algorithm NoRMCorre ^133^, implemented in Python through the CaImAn package ^134^. Registration was performed using large 120 × 120-pixel patches with a 60-pixel overlap to minimize local registration artifacts. Fluorescent nuclei were segmented from the temporal mean images of the motion-corrected recordings using a local-maxima detection algorithm described previously ^38^. For each detected nucleus, the fluorescence signal *F* (*t*) was extracted from a circular region of interest with a 3-pixel radius centered on the corresponding maximum. This radius represented a conservative lower bound on the expected nuclear radius and reduced signal contamination from neighboring cells. When regions of interest overlapped, pixels within the overlapping area were excluded from both regions. Raw fluorescence traces were corrected for slow baseline drift and converted to normalized fluorescence changes (Δ*F/F*_0_). To estimate the time-varying baseline *F*_0_(*t*) for each neuron, the fluorescence trace was first Gaussian-filtered (*σ* = 3 s), after which a 60-s minimum filter was applied. The resulting baseline was further smoothed using a broader Gaussian filter (*σ* = 10 s). Normalized fluorescence traces were then calculated as

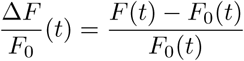

### Image unwarping

To correct geometric distortion along the horizontal scanning axis caused by the sinusoidal trajectory of the resonant mirror, we adapted a coordinate-mapping approach developed previously ^45^. Let *l* denote the index of a uniformly timed intensity sample acquired during one forward sweep of the resonant scan, and *l^′^* its corresponding position on an ideal linear scan. When samples span the complete forward sweep, the mapping is

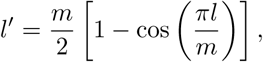

where *m* = 512 is the number of pixels along the scanning axis. In our acquisition scheme, PMT detection was restricted to the central portion of the mirror trajectory, excluding the most nonlinear regions near the mirror turning points. We therefore introduced a symmetric padding parameter *p*, representing the width of the excluded portion from each end of the complete sweep, and mapped the acquired coordinate *l* onto the corresponding full-sweep coordinate *l̃*:

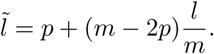

Substituting *l̃* for *l* in the first equation yields a non-integer projected coordinate for each of the acquired pixels. These coordinates were used as a one-dimensional lookup table to remap each scan line onto a uniform spatial grid using linear interpolation. Because *p* was not known a priori, we calibrated it separately for each imaging session by imaging 15-micron fluorescent beads at 11 predetermined positions along the horizontal field of view, then stacking individual bead images into a single calibration image. For each candidate value *p* ∈ [30, 200] (500 uniform samples), we unwarped the calibration image and measured the width of each bead from an intensity profile passing through their center, using an empirically validated intensity threshold. We retained the value of *p* that minimized the standard deviation of bead width across the 11 positions. Finally, because the unwarping approach altered the effective pixel size along the scanning axis, corrected images were resampled using linear interpolation to obtain a uniform pixel size of 1 × 1 *µ*m^2^. To validate the morphology of unwarped brains, we re-imaged a subset of animals using either a two-photon microscope equipped with a linear scanner or an epifluorescence microscope. We then used the length-width ratios of corresponding brain images as a simple morphological quantity for error evaluation.

### Brain atlas registration

To map calcium imaging data into a common three-dimensional coordinate space, we used a two-stage registration procedure described previously ^38^. At each stage, Advanced Normalization Tools (ANTs ^135^) was used to estimate a sequence of three transformations: an initial rigid transformation, followed by an affine transformation and, finally, a non-rigid transformation using the symmetric normalization (SyN) algorithm. We used parameters previously optimized for the registration of live zebrafish brain images ^136^:

`antsRegistration -d 3 --float 1 --output [,$filename aligned] --interpolation WelchWindowedSinc --use-histogram-matching 0 -r [$filename fixed,$filename moving,1] -t rigid[0.1]`

`-m MI[$filename fixed,$filename moving,1,32,Regular,0.25]`

`-c [200x200x200x0,1e-8,10] --shrink-factors 12x8x4x2 --smoothing-sigmas 4x3x2x1vox`

`-t Affine[0.1] -m MI[$filename fixed,$filename moving,1,32,Regular,0.25]`

`--convergence [200x200x200x0,1e-8,10] --shrink-factors 12x8x4x2 --smoothing-sigmas 4x3x2x1vox`

`-t SyN[0.2,6,0] -m CC[$filename fixed,$filename moving,1,2]`

`-c [200x200x200x200x10,1e-7,10] --shrink-factors 12x8x4x2x1 --smoothing-sigmas 4x3x2x1x0vox`

Mutual information was used as the similarity metric for the rigid and affine transformations, whereas cross-correlation was used for the non-rigid SyN transformation. $filename_fixed denotes the reference image stack, saved in .nrrd format with voxel dimensions specified in its header, and $filename_moving denotes the image stack to be transformed. In the first registration stage, each piezo-acquired calcium imaging volume was aligned with its corresponding anatomical brain scan. In the second stage, the same anatomical scan was aligned with the 6 dpf mapZebrain template generated from the same nuclear GCaMP transgenic line. After each stage, the transformed volume was overlaid on its corresponding fixed volume and visually inspected for gross registration errors. In some older animals, the elongated forebrain could not be adequately matched to the more compact anatomy of the 6 dpf template using the default parameters. In these cases, the SyN parameters were adjusted to accommodate larger deformations.

Following volumetric registration, the three-dimensional coordinates of neurons detected in the original calcium imaging planes were exported to a *.csv* file in physical units (*µ*m). The estimated transformations were then applied to these coordinates using antsApplyTransformsToPoints, first to map them from functional imaging space into anatomical scan space and subsequently from anatomical scan space into atlas space. The transformed coordinates were converted into atlas voxel coordinates, and each neuron was assigned to one of 64 anatomical brain regions by indexing the corresponding three-dimensional binary region masks (Supp. Table 1).

To generate an anatomical brain template for each developmental time point, individual anatomical scans were converted to .nii.gz format with the appropriate voxel dimensions and spatial metadata specified in the image headers. We then used the ANTs antsMultivariateTemplateConstruction2.sh script ^46^ to perform unbiased groupwise template construction. Four template-construction iterations were used to achieve clear delineation of fine anatomical structures and low-contrast boundaries.

### Quantification of brain morphology

To quantify global brain morphology, we first generated two-dimensional projections of each anatomical brain scan and manually measured three linear dimensions. Brain length was measured from the olfactory bulb to the caudal boundary of the brain at the level of the area postrema and the beginning of the spinal cord. Brain width was measured between the lateral extremities of the optic tectum, whereas brain thickness was measured from the torus longitudinalis to the ventralmost hypothalamic neurons. To estimate total brain volume, we manually segmented a three-dimensional brain mask on the 6 dpf atlas template and filtered the resulting mask to smooth and clean its boundaries. This mask was then projected onto each individual anatomical scan using the antsApplyTransforms command and the inverse transformations obtained during registration. Nearest-neighbor interpolation was used to preserve the binary nature of the transformed mask. The voxels contained within each transformed mask were then summed and converted into physical units using the corresponding voxel dimensions to obtain total brain volume. Regional brain volumes were quantified using the same procedure, with individual atlas region masks warped onto each animal’s anatomical scan.

### Comparison with adult brain

To contextualize larval brain growth and morphological changes, we compared our measurements with the Adult Zebrafish Brain Atlas (AZBA) ^47^. Adult brain volume was estimated by summing the volumes of all individual atlas regions. We then used this value as the endpoint of a simple exponential growth model, *v* = *α^t^v*_0_ where *v*_0_ was the mean brain volume at 4 dpf, *α* was the empirically measured daily volume growth factor (1.051), and *t* was the elapsed time in days. We solved for the time required for the larval brain to reach the estimated adult volume. To derive a 95% confidence interval, we repeatedly subsampled the 4 and 8 dpf populations (10 random animals per time point) and calculated the corresponding daily growth factor. The 2.5th and 97.5th percentiles of this distribution were then used as lower and upper growth-rate estimates, respectively. We also compared regional volumes between the larval and adult atlases. Because anatomical nomenclature differed between atlases, we manually inspected brain regions and constructed a correspondence table, which was subsequently used to match and correlate larval and adult regional volumes

### Cellular birthdating and confocal imaging

To visualize neurons born between 4 and 8 dpf, we sequentially labeled pan-neuronal nuclear HaloTags in Tg(*elavl3* :H2B- HaloTag) larvae using JF552 and JF635 dyes ^49^. At 4 dpf, groups of ~10 larvae were incubated for 2 h in 4 ml of embryo medium containing 0.3 *µ*M JF552, prepared from a 1 mM stock solution in DMSO. Larvae were then thoroughly rinsed with fresh embryo medium and returned to standard housing conditions. Four days later, the same larvae were incubated with JF635 for 2 h, washed, and embedded in agarose for dual-color confocal imaging. This sequential labeling strategy distinguished neurons already present at 4 dpf, which contained both fluorophores, from neurons born during the intervening period, which were labeled only by JF635. Imaging was completed within approximately 3 h of the second dye application, as longer delays allowed newly generated, unlabeled neurons to accumulate within the brain and appear as dark regions.

For confocal imaging, 8 dpf larvae were fully embedded upright in 2% low-melting-point agarose in glass-bottom Petri dishes. Images were acquired using a Zeiss LSM 880 upright confocal microscope equipped with a 20*×* apochromatic water-dipping objective (NA 1.0). JF552 and JF635 were simultaneously excited using 561 nm (DPSS 561-10) and 633 nm (HeNe633) lasers, respectively. Fluorescence emission was collected using two PMT detectors equipped with 515–620 nm and 650–754 nm band-pass filters for JF552 and JF635, respectively. Images were acquired at a pixel size of 1.004 *µ*m, with a pixel dwell time of 1.304 *µ*s and eightfold frame averaging. The confocal pinhole was adjusted to produce an optical section thickness of 5 *µ*m, and volumetric stacks were acquired using a 4 *µ*m axial step size. Two partially overlapping stacks of approximately 70 planes each were acquired per animal to cover the anterior and posterior brain. Whole-brain volumes were then reconstructed by stitching the two stacks using linear blending (Pairwise stitching plugin from Fiji ^137^).

### Segmentation of microscopy images

To segment fluorescently labeled nuclei in birthdating experiments and distinguish cell-body and neuropil compartments in fluorescent transgenic lines, we developed an iterative sparse-annotation workflow for training 2D U-Net models. Separate binary models were trained for each segmentation task (e.g., labeled nuclei, cell bodies, or neuropil versus background). For each model, we manually annotated several hundred 3D coordinates per animal using Fiji ^137^. Image stacks were visualized in dual-channel mode to simultaneously inspect the relevant fluorescence channels. Annotated coordinates were distributed approximately uniformly throughout the brain, with additional sampling in regions exhibiting weak fluorescence or low contrast between channels to ensure adequate representation of difficult cases. Before crop extraction, each optical section was lightly filtered using a Gaussian filter (*σ* = 0.5 pixels), and local contrast was enhanced using contrast-limited adaptive histogram equalization (CLAHE). A dataset of multichannel, 2D image crops (2 129 129 pixels) centered on the annotated coordinates was then assembled for each segmentation task (5191 total crops for Halotag confocal images, 7200 for neuropil and cell bodies, and 6092 for glutamatergic/GABAergic neurons). Crops from all animals were pooled and randomly divided into training and validation sets using a 90:10 ratio. We trained compact U-Net models with 32 filters in the initial convolutional layer using the Adam optimizer, a learning rate of *λ* = 5 10*^−^*^4^, a single-pixel binary cross-entropy loss, and 100 training epochs. Training data were augmented through random rotations and reflections. Each crop was additionally reduced to the model input size (64 × 64) using a randomly displaced sub-crop, such that the annotated pixel was not systematically positioned at its center. The binary cross-entropy loss was evaluated only at the location of the annotated pixel, allowing the network to learn from sparse categorical annotations without requiring manually segmented masks. These random displacements of the annotated pixel within the input field reduced center-position bias and encouraged the model to generate spatially coherent predictions. Models were trained iteratively: after each training round, output segmentations were visually inspected, additional coordinates were annotated in regions where the model failed, and the expanded dataset was used for the subsequent training round. Final models achieved F1 scores of 0.98–0.99 on the training set and 0.96–0.97 on the validation set. Following training, each optical section was segmented independently using overlapping tiled inference, with a tile size of 64×64 pixels and an overlap of 24 pixels between adjacent tiles. Predictions from overlapping tiles were recombined using smooth spatial weighting to minimize discontinuities at tile boundaries. The resulting combined model outputs were thresholded at 0.5 to obtain binary segmentation maps, which were subsequently assembled into 3D volumes. These volumes were transformed into a common anatomical reference space using antsApplyTransforms (with nearest neighbor interpolation) and the transformations previously estimated from the corresponding GCaMP channel. Co-registered binary segmentations were then averaged across animals and lightly spatially filtered to generate population-level probability maps, in which each voxel represented the fraction of animals exhibiting a segmented feature at that location. All models were implemented and trained using PyTorch on a GPU with 8 GB of memory, using a batch size of 64.

### Measuring functional connectivity

To obtain a mesoscale representation of brain activity that could be statistically compared across animals, we assigned individual neurons to a common set of atlas-based anatomical nodes. We initially distributed 400 nodes (200 per hemisphere) across 64 brain regions in proportion to their volumes, such that larger structures like the optic tectum and the pallium received a greater number of nodes. Each regional three-dimensional mask was then subdivided into its allocated number of nodes using an iterative *K*-means clustering procedure. Specifically, the voxel coordinates within each regional mask were independently clustered 25 times, and the centroids obtained across all runs were subsequently pooled and re-clustered to identify a stable set of consensus centroids. Each voxel was then assigned to its nearest consensus centroid, effectively partitioning the regional mask into three-dimensional Voronoi cells.

Individual neurons were assigned to the resulting subregional masks according to their registered coordinates. We retained only nodes containing at least two neurons in every one of the 79 experiments in both hemispheres. This stringent sampling criterion restricted the analysis to a common, bilaterally symmetrical brain volume that was reliably imaged across all animals. It also excluded regions susceptible to inconsistent sampling, particularly the most ventral brain regions and the caudal extremity of the brain, which was occasionally truncated in larger animals or in larvae positioned at a slight downward angle in the agarose. After applying this criterion, 278 nodes (139 per hemisphere) were retained for subsequent analyses.

Following anatomical assignment, the calcium activity of all neurons within each node was averaged to obtain a representative mesoscale time series. Pairwise Pearson correlation coefficients were then calculated between all node time series, yielding matrices that were interpreted as functional connectivity. Spontaneous functional connectivity, FC_spont_, was calculated from the initial 15-min spontaneous recording period after excluding the first minute to minimize imaging onset effects. Stimulus-evoked functional connectivity, FC_stim_, was calculated from the subsequent 15-min visual-stimulation period and therefore captured the combined correlation structure induced by the full set of presented visual stimuli.

### Measuring signal properties

In Supp. Fig. S8, we characterized several imaging and signal properties across development. To estimate the apparent spatial footprint of individual neurons, we extracted raw fluorescence values surrounding each detected neuronal centroid and averaged these image patches across all neurons. Gaussian functions were then fitted to the horizontal and vertical intensity profiles of the average fluorescence footprint, and their full widths at half maximum (FWHMs) were used to quantify the apparent neuronal diameter along each axis. Vertical FWHM values decreased slightly with age, consistent with the larger scanning amplitude required to image the larger brains of older animals.

Noise levels in the calcium-imaging time series were quantified following the frame-rate-normalized definition introduced by Rupprecht *et al.* ^138^:

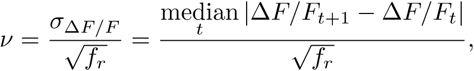

where *σ*_Δ*F/F*_ represents the magnitude of frame-to-frame fluorescence fluctuations and *f_r_* is the imaging frame rate. This normalization accounts for the dependence of shot noise on sampling rate and allows noise levels to be compared across recordings acquired at different frame rates, although all our recordings were made using identical parameters. Signal amplitude was defined as the difference between the 95th and 5th percentiles of each Δ*F/F* time series. The signal-to-noise ratio was then calculated by dividing this signal amplitude by the corresponding noise level, *ν*.

### Global and motor signal regression

For specific analyses, we regressed either the global or motor signal from each node time series. The global signal was defined as the activity averaged across all network nodes, whereas the motor signal was defined as the mean activity of all *M* + neurons. For each node independently, the selected signal was fitted to its activity using linear regression with an intercept, and the fitted component was subtracted to obtain a residual time series. Pairwise Pearson correlation coefficients were then calculated between the residual time series to generate global-signal-regressed or motor-signal-regressed FC matrices.

### Connectivity-distance analysis

To identify the individual edges driving developmental changes in the relationship between functional connectivity and anatomical distance, we decomposed the Pearson correlation coefficient into edge-wise contributions. For a set of *M* undirected edges, the correlation between functional connectivity *W* and Euclidean distance *D* can be written as

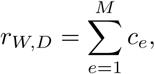

where the contribution of edge *e* is

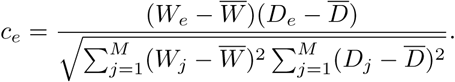

The sum of these edge-wise contributions therefore exactly recovers the Pearson correlation coefficient. Positive contributions indicate edges for which FC and distance deviate from their respective means in the same direction, whereas negative contributions indicate deviations in opposite directions. We calculated *c_e_* separately from the group-averaged FC matrices at 4 and 8 dpf and defined the developmental change in contribution as 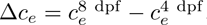 . Positive values of Δ*c_e_* indicate edges that shifted the FC–distance correlation in the positive direction over development, whereas negative values shifted it in the negative direction. For simplicity, we inverted the contribution signs in the main text, such that “positive” contributions refer to those that strengthened the negative FC–distance relationship by increasing its absolute magnitude.

We analyzed positive and negative changes separately and retained edges within the upper 10% of absolute contribution changes. Within both signs, the selected edges formed distinct, non-overlapping short- and long-distance distributions, allowing them to be classified into four categories: short-range positive, long-range positive, short-range negative, and long-range negative contributions to the developmental change in the FC–distance relationship. For visualization and subsequent analyses, these categories were displayed using their observed FC changes between 4 and 8 dpf, 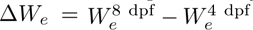, rather than their normalized edge-wise contribution values. Note that Spearman rank correlations were used in Fig. 4b,c to accommodate the nonlinear relationship between functional connectivity and anatomical distance, while edge-wise contributions were subsequently derived from Pearson correlations because their additive decomposition permits a direct quantitative interpretation of each edge’s contribution to the overall correlation. This methodological distinction did not affect our conclusions as both measures produced highly similar developmental trends.

### Detection of network modules

To identify functional modules across multiple spatial scales, we applied a hierarchical consensus-clustering procedure based on the Louvain community-detection algorithm ^63^, following an approach used in our previous study ^38^. For each group-averaged global-signal-regressed (GSR) FC matrix, clipped in the [0, 1] range, we generated an ensemble of 1000 partitions by maximizing undirected, unsigned modularity across resolution parameters *γ* ranging from 0.25 to 10.0. This range produced partitions containing between approximately 2 and 100 modules. We then constructed a consensus matrix *C*, whose elements *c_ij_* represented the fraction of partitions in which nodes *i* and *j* were assigned to the same module:

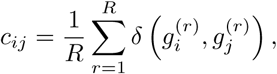

where *R* = 1000, 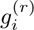 denotes the module assignment of node *i* in partition *r*, and *δ* is the Kronecker delta. We transformed co-assignment probabilities into distances using *D^C^* = 1 *C* and applied agglomerative hierarchical clustering with Ward’s linkage criterion. Partitions containing different numbers of modules were obtained by cutting the resulting dendrogram at successive levels using the maxclust criterion. For each hierarchical partition, we quantified modularity as ^139^

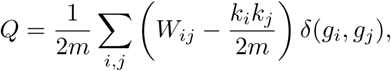

where *W_ij_*is the FC weight between nodes *i* and *j*, *k_i_* = *_j_ W_ij_* is the weighted degree of node *i*, 2*m* = ∑*_i,j_ W_ij_* is the total network weight, and *g_i_*denotes the module assignment of node *i*. Modularity was calculated using the Brain Connectivity Toolbox ^140^ implementation in Python (bctpy).

To determine whether the observed modular organization could be explained solely by spatial proximity, we generated deterministic geometric surrogate matrices in which connectivity decayed exponentially with Euclidean distance:

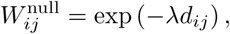

where *d_ij_* is the Euclidean distance between nodes *i* and *j*, and *λ* is the distance-decay parameter. For each empirical group-averaged FC matrix, including each leave-one-out estimate, *λ* was selected to maximize the Pearson correlation between the empirical and geometric connectivity matrices. Hierarchical consensus modules were then identified from the geometric surrogate matrices using the same procedure applied to the empirical data.

### Functional clustering algorithm

To identify functionally coherent neuronal ensembles without imposing a predetermined final number of clusters, we developed an iterative clustering procedure inspired by a previous zebrafish study ^67^.

Before clustering, neuronal activity traces were lightly temporally filtered (gaussian *σ* = 1) and denoised using PCA by reconstructing the time series matrix using its first 200 temporal components, thereby retaining shared population-level dynamics while suppressing neuron-specific fluctuations, such as noise, not captured by the low-dimensional representation. Each reconstructed neuronal time series was subsequently z-scored across time before clustering. After preprocessing, neurons were partitioned into *K* = 1000 initial clusters using *K*-means clustering with Euclidean distance. The initial clusters were then refined through three iterations of the following procedure.

First, neurons whose activity correlated with their cluster centroid below a threshold *r*_1_ were removed and placed in an unassigned pool. Clusters whose mean pairwise correlation between constituent neurons was below a second threshold, *r*_2_, were subsequently discarded. To merge redundant clusters, we constructed an undirected graph in which clusters were connected when the correlation between their centroids exceeded a third threshold, *r*_3_. All clusters belonging to the same connected component were then merged, allowing transitive merging when clusters were linked through one or more intermediate clusters. This transitive criterion allowed smoothly varying manifestations of a shared activity pattern, such as motor-related signals, to be consolidated into a single cluster without requiring every pair of constituent centroids to exceed the threshold. Merging was performed simultaneously using centroids calculated before the merging step rather than through sequential pairwise updates. Following merging, each neuron in the unassigned pool was reassigned to the cluster with the most highly correlated centroid, provided that this correlation exceeded *r*_1_. Clusters containing fewer than five neurons were then discarded. Cluster centroids were recalculated following changes in cluster membership and during each subsequent refinement iteration.

After three iterations, the procedure yielded a data-driven number of functionally coherent clusters, together with a pool of unassigned neurons whose activity did not reliably match any retained cluster and generally exhibited low signal quality. We used empirically validated thresholds (*r*_1_*, r*_2_*, r*_3_) = (0.2, 0.1, 0.75), selected to accommodate the comparatively low signal-to-noise ratio of single-neuron calcium traces acquired during brain-wide two-photon imaging. The high merge threshold *r*_3_ constrained transitive merging to limit chaining between weakly related activity patterns.

### Identifying event-correlated neurons

To identify stimulus-responsive neurons (SRNs), we quantified the change in Δ*F/F* between pre-stimulus and stimulus-response windows for each neuron and for each stimulus type. Fluorescence was averaged over five time points within each window, and the difference between windows was calculated separately for each of the six stimulus repetitions. The resulting fluorescence differences were then averaged across repetitions to obtain a single response statistic for each neuron. A null distribution was generated by repeating this procedure using randomly selected onset times. Empirical *P* -values were calculated from the percentile of the observed response statistic within its corresponding null distribution. Multiple comparisons were corrected independently within each animal using the Benjamini–Hochberg procedure, and neurons with an FDR-corrected *P <* 0.05 were classified as SRNs.

Motor-correlated neurons (*M* + neurons) were identified using an analogous procedure, with fluorescence differences calculated between pre- and post-movement windows surrounding detected swimming events. Unlike the visual stimuli, the number of available motor events varied across animals. Animal 29 exhibited an exceptionally small number of swimming events, resulting in insufficient statistical power to identify *M* + neurons after FDR correction. We therefore used the uncorrected *P* -values for this animal.

### Projectome gradient analysis

To contextualize the anatomical reorganization of SRNs, we used the single-neuron morphology dataset from mapZebrain^42^ to identify RR regions and their downstream projection pathways. Regions containing at least 10 retinal ganglion cell (RGC) neurite terminals were classified as retinorecipient. This criterion identified three principal RR domains—the optic tectum, pretectum, and dorsal thalamus—with the principal retinal targets described in the zebrafish literature.

We next constructed a directed binary projectome from 4,327 reconstructed neuronal morphologies ^42^. A directed connection from region *i* to region *j* was retained when neurons originating in region *i* contributed at least five neurite terminals to region *j*, thereby excluding weak projections. The position of each region within the visual projection hierarchy was then defined as its shortest directed path length from any RR region. This procedure organized the projectome into four successive layers, consistent with recent estimates obtained from an independent neuronal morphology dataset ^52^.

To evaluate whether the observed relationship between SRN organization and projectome hierarchy exceeded that expected from spatial embedding and random distance-dependent connectivity, we generated surrogate projectomes using an empirically supported exponential distance rule ^71,57^. In these surrogates, directed binary edges were sampled independently according to

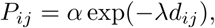

where *d_ij_* is the Euclidean distance between regions *i* and *j*, *α* controls overall connection density, and *λ* controls the rate at which connection probability decays with distance. The model parameters were fitted sequentially. First, *λ* was selected to minimize the Kolmogorov–Smirnov distance between the empirical and model connection-length distributions, sampled over multiple surrogate graphs. With *λ* fixed, *α* was then adjusted such that the expected density of the surrogate projectomes matched the empirical projectome density. This procedure yielded *λ*_opt_ = 0.00637 and *α*_opt_ = 0.624.

### Stimulus regression analysis

To complement neuron-count-based analyses of stimulus responsiveness, we quantified the extent to which visual stimuli explained mesoscale activity. For each of the seven stimulus categories, we generated a binary time series sampled at the volumetric imaging rate and convolved it with an exponential kernel, *h*(*t*) = *e^−t/τ^*for *t* 0. We used *τ* = 3.5 s, consistent with previous work ^38^, to approximate the slow fluorescence response of nuclear-localized H2B-GCaMP6s. For every animal and network node, activity was fitted using multiple linear regression with the seven stimulus regressors and an intercept. Model fit was quantified using the coefficient of determination, *R*^2^, and values from corresponding nodes in the two hemispheres were averaged, yielding 139 bilateral anatomical locations. Developmental changes in stimulus- related explanatory power were then assessed at each location by calculating the Pearson correlation between *R*^2^ and age across all 79 animals. Two-sided *P* -values were calculated from the beta null distribution as implemented in scipy, and FDR-corrected across the 139 anatomical locations.

### CNN training

To examine whether visual representations reorganize across successive processing stages during learning, we trained a simple feedforward CNN, an architecture commonly used as a hierarchical model of biological visual processing ^141^. The network received single-channel 28×28 images as input. Its first layer, representing the retina, consisted of 16 convolutional filters with a kernel size of 5 × 5, a padding of 2 pixels, and no bias term. Convolutional outputs were passed through a rectified linear unit (ReLU) activation function and downsampled using 2 × 2 average pooling, yielding 16 feature maps of 14 14 pixels. These features were flattened and propagated through four fully connected layers containing 128 units each, with a ReLU activation applied after every layer. A final linear layer containing 10 output units produced the logits used for digit classification.

The network was trained to classify handwritten digits from the MNIST dataset using the Adam optimizer, a learning rate of 10*^−^*^3^, a batch size of 128, and a cross-entropy classification loss. To impose a global metabolic constraint on network activity, we supplemented the classification loss with a mean-squared activation penalty calculated across all units in the convolutional and fully connected hidden layers:

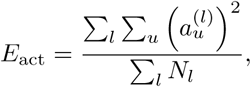

where 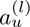 is the activation of unit *u* in layer *l* and *N_l_* training objective was is the number of activation values in that layer. The complete

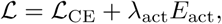

with an activity regularization coefficient of *λ*_act_ = 20. To compare activity across learning stages, we extracted network activations after 20 iterations (weight updates), when test accuracy was approximately 50%, and after 400 iterations, when test accuracy reached approximately 98%. At each checkpoint, activations were evaluated over the complete MNIST training set and averaged across both samples and units separately for each of the four fully connected layers. Because ReLU activations are non-negative, these averages provided a direct measure of activity magnitude without cancellation between positive and negative unit responses. Early and late activations were always compared within the same training trajectory. The complete procedure was repeated across 10 independently initialized models, and activation trajectories were subsequently averaged across models. Models were implemented and trained using PyTorch on a GPU with 8 GB of memory.

### Tail tracking and swim bout detection

To reconstruct tail posture in each behavioral frame, we used an intensity-based tracking algorithm described previously ^142,38^. Briefly, the static background was removed by subtracting the temporal intensity mode of each pixel, after which recordings were Gaussian-filtered and locally contrast-enhanced using CLAHE. Filter parameters were adjusted for each animal to accommodate differences in recording quality and tail length. A tail base point was manually positioned near the swim bladder, and the tail was represented by 10 segments of equal length. Each successive segment was traced by searching a 120*^◦^* angular range in 6*^◦^*increments and selecting the direction with the highest line-profile intensity. Tracking coordinates were refined using the local intensity center of mass within a 5-pixel window, providing subpixel resolution.

Segment angles were calculated relative to the main body axis, producing 10 angular time series per recording. Angles were unwrapped to eliminate discontinuities at *π* and *π* and mean-centered such that the resting tail position corresponded to zero. Sparse tracking errors were removed by applying two consecutive three-frame median filters, corresponding to approximately 7.5 ms at the camera sampling rate, followed by Gaussian smoothing with *σ* = 1 frame.

Swimming events were segmented from the 10 tail-angle time series using a temporal convolutional network (TCN) comprising residual blocks with exponentially increasing dilation factors ^73^. Frame-wise binary swim annotations were generated using a custom GUI and included 801 manually annotated swimming events from several animals. Training examples consisted of 2,000-frame windows (~ 5 s), sampled with equal probability from periods containing at least one swimming event or no swimming events. To augment the dataset and ensure invariance to swimming direction, each training example was duplicated after inversion of all tail angles while retaining the same binary target sequence. This procedure produced 40,000 training examples.

The network first projected the 10 input traces onto 64 hidden channels using a 1 × 1 convolution. These representations were passed through five residual blocks with dilation factors of 1, 2, 4, 8, and 16. Each block contained two one-dimensional convolutions with a kernel size of 5, each followed by batch normalization, a ReLU activation, and dropout with a probability of 0.1. A final 1 × 1 convolution produced one swim-detection logit per camera frame. The model was trained for one epoch using binary cross-entropy with logits and the Adam optimizer, with a learning rate of 10*^−^*^3^, a weight decay of 10*^−^*^4^, a batch size of 64, and gradient clipping at a maximum norm of 1.0.

For inference, complete recordings were divided into overlapping 16,384-frame windows with an overlap of 1024 frames. Predictions within overlapping regions were averaged, and the resulting logits were transformed into frame-wise swimming probabilities using a sigmoid function. Probabilities above 0.5 were classified as swimming. Model predictions were visually inspected across all recordings using the annotation interface to confirm appropriate detection of swimming events.

### Forebrain analysis

To isolate forebrain dynamics, we concatenated the mesoscale activity matrices from all animals along the temporal dimension and applied ICA separately to the spontaneous and visual-stimulation periods. ICA was performed using 10 components, from which we identified a component with spatial loadings concentrated in the forebrain. This component consistently emerged whenever at least three components were extracted (Supp. Fig. S14a). The concatenated component time course was subsequently divided to recover an individual time series for each of the 79 animals. To characterize the temporal persistence of this signal, we calculated its integrated autocorrelation time ^143^ independently in each animal:

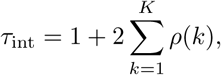

where *ρ*(*k*) is the normalized autocorrelation at lag *k* and *K* is the first lag at which the autocorrelation crossed zero. The resulting values were expressed in seconds using the volumetric sampling rate. Frequency content was next quantified using the fast Fourier transform (FFT). For each animal, we retained the positive-frequency coefficients, calculated their amplitudes, and normalized them by their sum. Normalized spectra were then aggregated into 50 frequency bins for comparison. Developmental differences were tested separately within each bin using K-W tests across age groups, followed by FDR correction across bins (Fig. 8g). Differences between the spontaneous and visual-stimulation periods were evaluated within each frequency bin using paired Wilcoxon signed-rank tests, with FDR correction applied across bins (Fig. 8h). To identify independent-component neurons (ICNs), we calculated the Pearson correlation between the corresponding ICA time course and the calcium activity of every neuron within each animal. Statistical significance was assessed using circular time-shift surrogates. Circular permutations preserved the internal temporal structure and amplitude spectrum of the ICA signal while disrupting its alignment with neuronal activity. The 10 smallest shifts in either temporal direction were excluded because they retained substantial alignment with the original signal owing to its autocorrelation, leaving 400 surrogate shifts per neuron. One-sided empirical *P* -values were calculated as the proportion of surrogate correlations equal to or greater than the observed correlation, followed by standard FDR correction across neurons. To quantify the local spatial enrichment of ICNs, we calculated neuronal-class composition (either SRN, M+ or ICN) within the 100 nearest spatial neighbors of every neuron, based on their registered three-dimensional coordinates. For neuron *i*, the local dominance of ICNs was defined as

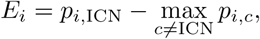

where *p_i,c_* is the proportion of neighboring neurons belonging to class *c*. Positive values therefore indicated that ICNs were more locally prevalent than any other neuronal class. To generate a null distribution, neuronal class labels were randomly permuted across coordinates 100 times, thereby preserving the total number of neurons in each class while eliminating their spatial organization. Local compositions and ICN-dominance scores were recalculated after each permutation and compared with the empirical values.

### Statistics

Standard statistical tests are specified in the main text and figure legends. Unless otherwise stated, all tests were two-sided and statistical significance was defined as *P <* 0.05. For families of related comparisons, *P* -values were adjusted using the Benjamini–Hochberg procedure to control the false discovery rate at *q* = 0.05. These results are denoted as “FDR-corrected” throughout the manuscript. Specialized statistical procedures are described below.

In Fig. 3e, developmental changes in functional connectivity were assessed using the network-based statistic (NBS) ^54^. Two-sided independent-samples *t*-tests were first performed for every network edge, and edges passing an initial threshold of *P <* 0.05 were grouped into connected components. Statistical significance was then evaluated at the component level by permuting age-group assignments 1000 times and comparing the number of edges in each empirical component with the maximum component size obtained under permutation. For both spontaneous and stimulus-evoked FC, the observed component was larger than those obtained in all 1000 permutations (*P <* 0.001).

In Fig. 3m, the significance of correlations between regional brain maps was assessed using BrainSMASH ^144^. BrainSMASH generates surrogate spatial maps whose variograms approximately match that of the empirical map, thereby approximately preserving its spatial autocorrelation while randomizing its regional organization. Empirical correlations exceeded those of 1000 surrogates (*P <* 0.001).

In Fig. 6j, retinorecipient (RR) labels were randomly permuted across regions 100,000 times while preserving the number of regions assigned to each group. For each permutation, we recalculated the difference between the group means to generate a null distribution. The null distribution was approximated as Gaussian using its empirical mean, *µ*_0_, and sample standard deviation, *σ*_0_, and the observed statistic, *T*_obs_, was standardized as 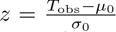 . A one-sided lower-tail *P* -value was calculated as *P* = Φ(*z*), where Φ is the standard normal cumulative distribution function. A similar procedure was used in Fig. 8r to identify neurons located within areas of significant local ICN enrichment. Neuronal class labels were randomly permuted across spatial coordinates 100 times, and the local ICN-dominance score was recalculated for every neuron after each permutation. For each neuron, the observed dominance score was compared with a Gaussian approximation of its shuffled-label null distribution using a one-sided upper-tail test. The resulting *P* -values were FDR-corrected across neurons.

## Data availability

The processed data used in this study will be made publicly available in the Université Laval collection on Borealis.

## Code availability

Data were analyzed using custom Python code available at https://github.com/PDKlab/neurodevelopment.

## Acknowledgements

We are grateful to Minoru Koyama for sharing the Tg(*elavl3* :H2B-HaloTag) line and for stimulating discussions in the early stages of this project. We are grateful to Sandra Migneault for assistance with animal care and to Pauline Marc-Tudor for contributing to the analysis of a pilot dataset during the summer of 2023. We also thank Vincent Breton-Provencher, Daniel Barabasi, Florian Engert, Anh-Tuan Trinh and Emre Yaksi for insightful discussions. A.L. was supported by PhD scholarships from the Natural Sciences and Engineering Research Council of Canada (NSERC), Fonds de recherche du Quebec–Nature et technologies (FRQNT), and Unifying Neuroscience and Artificial Intelligence–Quebec (UNIQUE). The zebrafish work was funded by the NSERC (RGPIN-2019-06887 and RGPIN-2024-06492 to P.D. and RGPIN-2023-05980 to P.D.K.), the Sentinel North program of Universite Laval (Canada First Research Excellence Fund), the Northern Contaminants Program of Canada, and the Next Generation Networks for Neuroscience (Neuronex)/FRQS (295823).

## Author contributions

A.L., P.D. and P.D.K. conceived the study. A.L. performed the calcium imaging experiments. M.Leb. conducted free-swimming behavioral experiments. A.L., M.A.E.B and H.P. performed additional anatomical imaging experiments. A.L. developed software and analyzed the data. P.D. and P.D.K. supervised the project. A.L. wrote the first draft. All authors revised the manuscript and approved the final version.

## Supplementary Material

### Supplementary fgures

**Figure S1:**
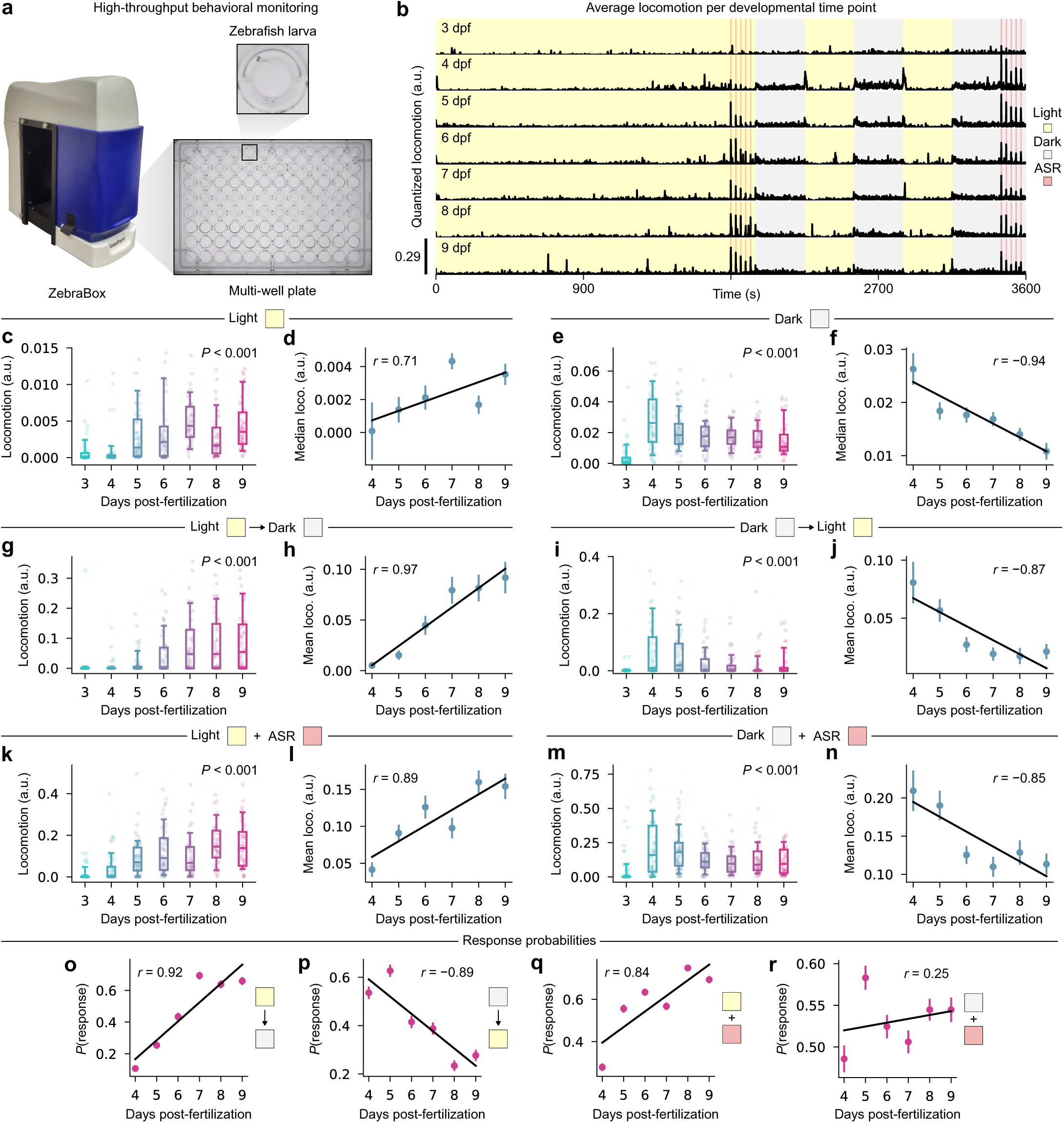
Supplementary analysis of free-swimming behavior. (**a**) Schematization of the behavioral monitoring system. (**b**) Average locomotion over time for 6 different age groups annotated in the top left corner; shaded regions indicate different sensory stimuli. (**c**) Distributions of locomotion averaged over the main light period; boxplot whiskers indicate 5th and 95th percentiles, *P <* 0.001, K-W test. (**d**) Linear regression of distribution averages from the previous plot; error bars indicate SEM, Pearson’s *r* = 0.71. (**e**) Distributions of locomotion averaged over the main dark periods; *P <* 0.001, K-W test. (**f**) Linear regression of distribution averages from the previous plot; Pearson’s *r* = *−*0.94. (**g**) Distributions of locomotion averaged after light to dark transitions; *P <* 0.001, K-W test. (**h**) Linear regression of distribution averages from the previous plot; Pearson’s *r* = 0.97. (**i**) Distributions of locomotion averaged after dark to light transitions; *P <* 0.001, K-W test. (**j**) Linear regression of distribution averages from the previous plot; Pearson’s *r* = *−*0.87. (**k**) Distributions of locomotion averaged during acoustic stimuli in ambient light; *P <* 0.001, K-W test. (**l**) Linear regression of distribution averages from the previous plot; Pearson’s *r* = 0.89. (**m**) Distributions of locomotion averaged during acoustic stimuli in darkness; *P <* 0.001, K-W test. (**n**) Linear regression of distribution averages from the previous plot; Pearson’s *r* = *−*0.85. (**o**-**r**) Probability of swimming (that is, locomotion greater than zero) in response to light-to-dark (**o**), dark-to-light (**p**), acoustic startles in ambient light (**q**) and acoustic startles in darkness (**r**), averaged over trials; linear regression coefficients are indicated on each plot.

**Figure S2:**
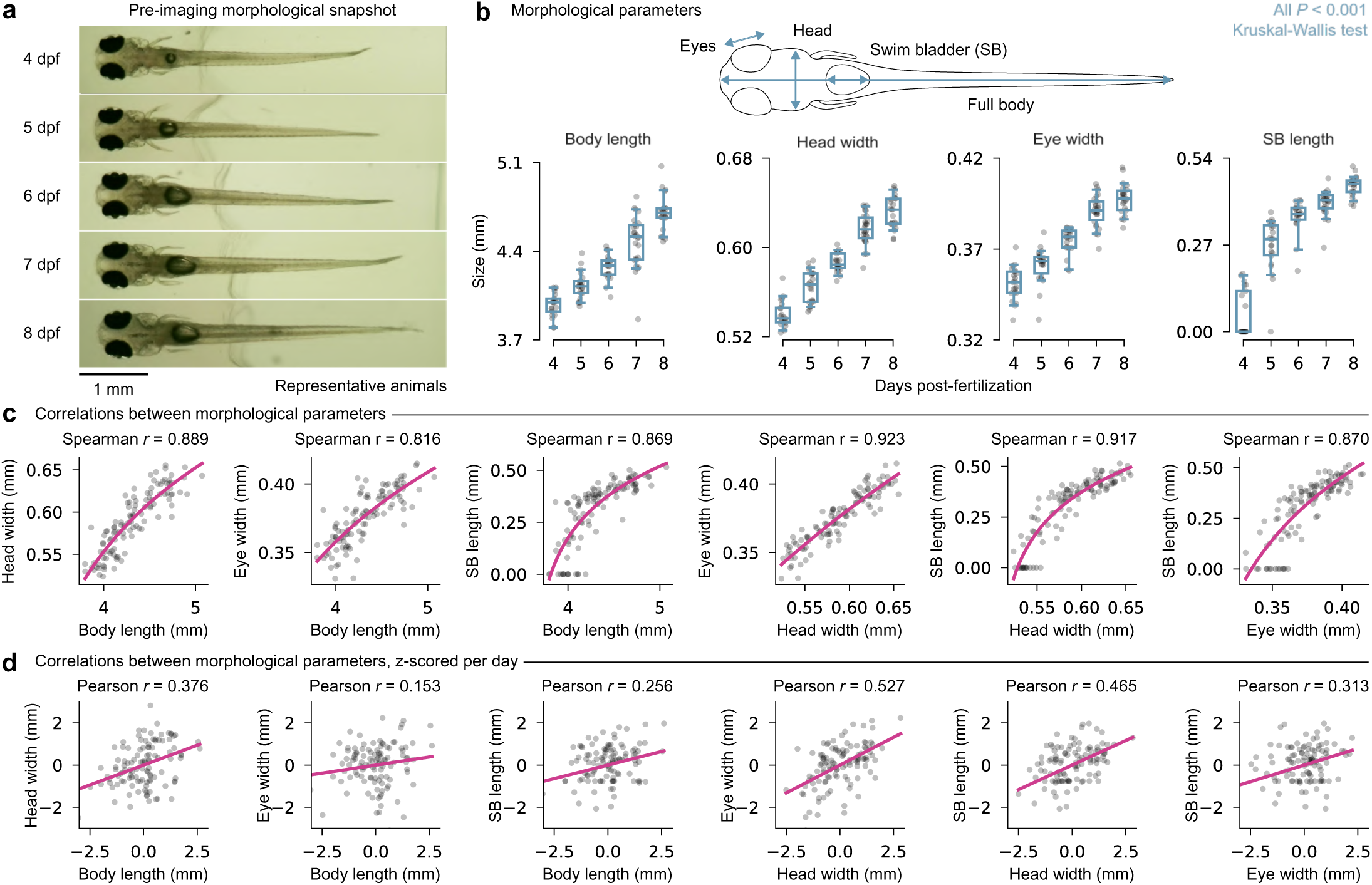
Larval growth curves. (**a**) Representative photographs of larvae at each developmental stage used for subsequent morphological analyses. (**b**) Top, schematic of the morphological measurements; bottom, body length, head width, eye width, and swim bladder (SB) length across development; Boxplot whiskers indicate the 5th and 95th percentiles; All measures increased significantly with age (K-W test, all *P <* 0.01); one larva at 7 dpf exhibited a markedly smaller body length than the rest of the cohort but showed no other obvious abnormalities and was therefore retained in all analyses. (**c**) Pairwise Spearman correlations between morphological features. Nonlinear curve fits are shown to highlight trends. Head width emerged as the strongest predictor of the other morphological measures. (**d**) Pairwise correlations after z-scoring each feature within time points to assess whether relationships persisted independently of developmental stage. All correlations stayed positive after normalization, with head width remaining the feature most strongly correlated with the others.

**Figure S3:**
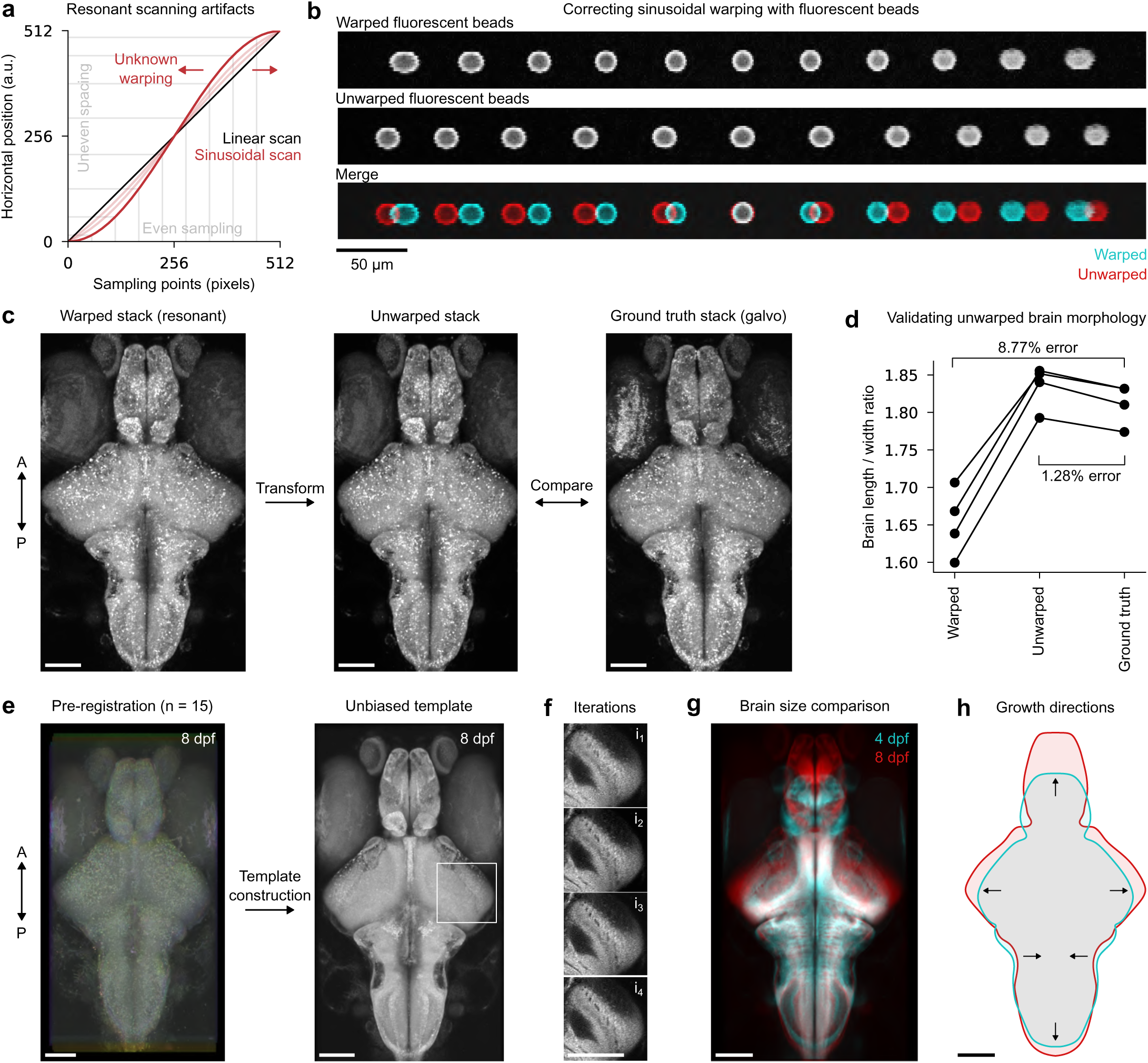
Brain unwarping approach and anatomical template building. (**a**) Due to the sinusoidal scanning pattern of the resonant mirror, uniformly spaced pixels within the horizontal axis of the image (*x* axis of the plot) are not uniformly distributed in space (*y* axis); this nonlinear projection can be inverted to retrieve the unwarped images, as long as the sinusoidal warping is measured using objects of known morphology (see Methods for details). (**b**) Top, horizontal warping of a single fluorescent bead positioned at different horizontal coordinates within the field of view; middle, numerically unwarped beads; bottom, color merge of the two previous images; note that the central bead is slightly wider after unwarping, which is later corrected through rescaling. (**c**) Maximum intensity projections of anatomical brain scans; once the warping is learned from fluorescent beads, the raw microscopy stacks (left) are unwarped (middle), which slightly narrows the brain; to validate this approach, we compared with images of the same brain acquired on a different microscope equipped with linear scanning (right). (**d**) Quantification of the morphological error introduced by sinusoidal warping or by the numerical unwarping procedure; prior to unwarping, brain width is exagerated, which lowers its length/width ratio; the true ratio is nearly recovered following the unwarping (*n* = 4 larvae, ground-truth two-photon or epifluorescence images). (**e**) Left, pseudocolor merge of 15 anatomical brain scans prior to alignment; right, the same brain scans after iterative registration to produce a template brain. (**f**) Example crop near the ventral optic tectum/torus semicircularis highlighting small morphological holes (vasculature) that become progressively refined through each template refinement iteration; notice little improvement between iterations 3 and 4. (**g**) Color merge of mean intensity projections from 4 and 8 dpf template brains, aligned to the midbrain-hindbrain boundary. (**h**) Color brain outlines drawn manually from the previous panel, with arrows indicating various growth directions; notice the shrinking of the anterior part of the hindbrain. A, anterior; P, posterior. Scale bars: 100 *µ*m.

**Figure S4:**
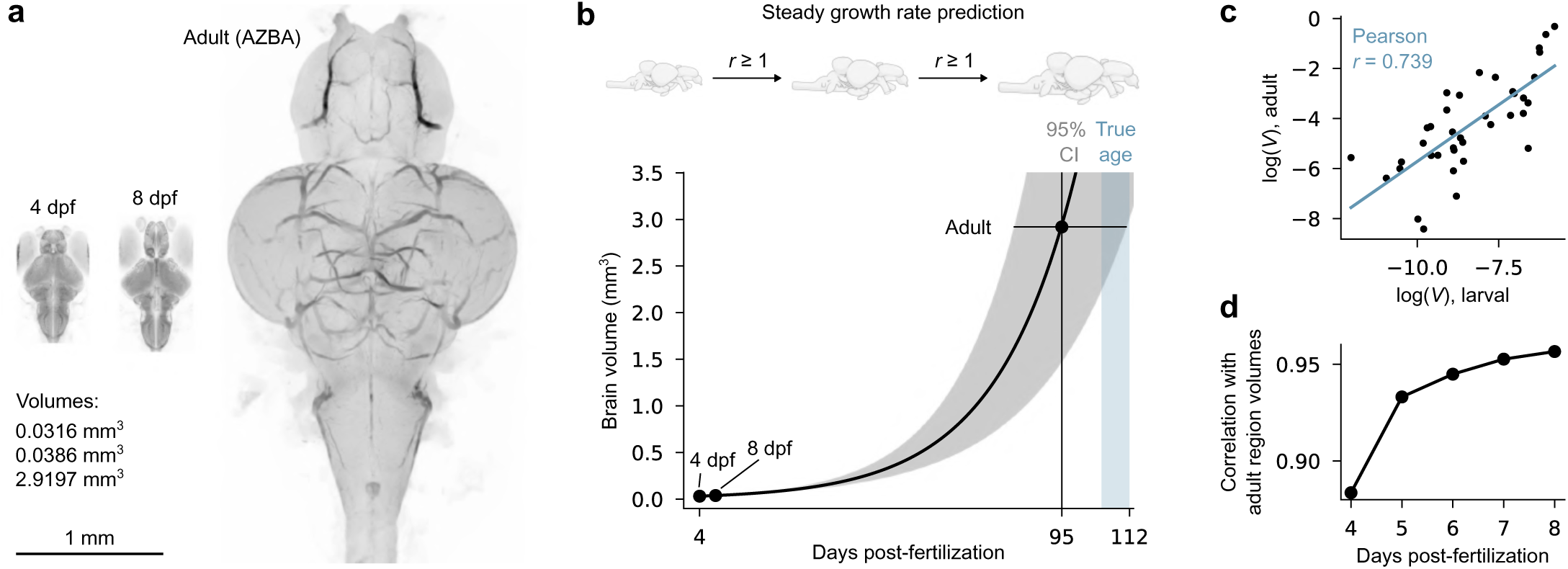
Comparison between larval and adult brains. (**a**) Side-by-side maximum-intensity projections of 4 dpf, 8 dpf, and adult zebrafish brains; brain volumes are indicated in the lower-left panel. Note that the adult brain image, obtained from the Adult Zebrafish Brain Atlas (AZBA), was generated using intrinsic autofluorescence and therefore exhibits different contrast than the larval images. (**b**) Extrapolation of larval brain growth to predict the time required to reach adult brain volume. The shaded region indicates the 95% confidence interval (CI) derived from the variability of brain volumes measured at 4 and 8 dpf, yielding a range of growth-rate estimates. The green shaded region denotes the ground-truth age range (105-112 dpf) of the adult brains used in AZBA. Predicted adult brain volume is reached after approximately 80-110 days, overlapping with the observed age range and suggesting that early larval growth rates are approximately maintained during the first months of life. (**c**) Correlation between log-transformed regional brain volumes at 6 dpf and adulthood (Pearson *r* = 0.739). Homologous regions between the larval and adult atlases were identified based on anatomical location and nomenclature (38*/*64 larval regions matched to adult regions). Similar correlations were observed for other larval time points (not shown). (**d**) Correlations between adult regional volumes and the 10 largest brain regions at each larval developmental stage. Larval-adult similarity increased steadily over development, indicating that relative brain-region proportions progressively converge toward adult proportions.

**Figure S5:**
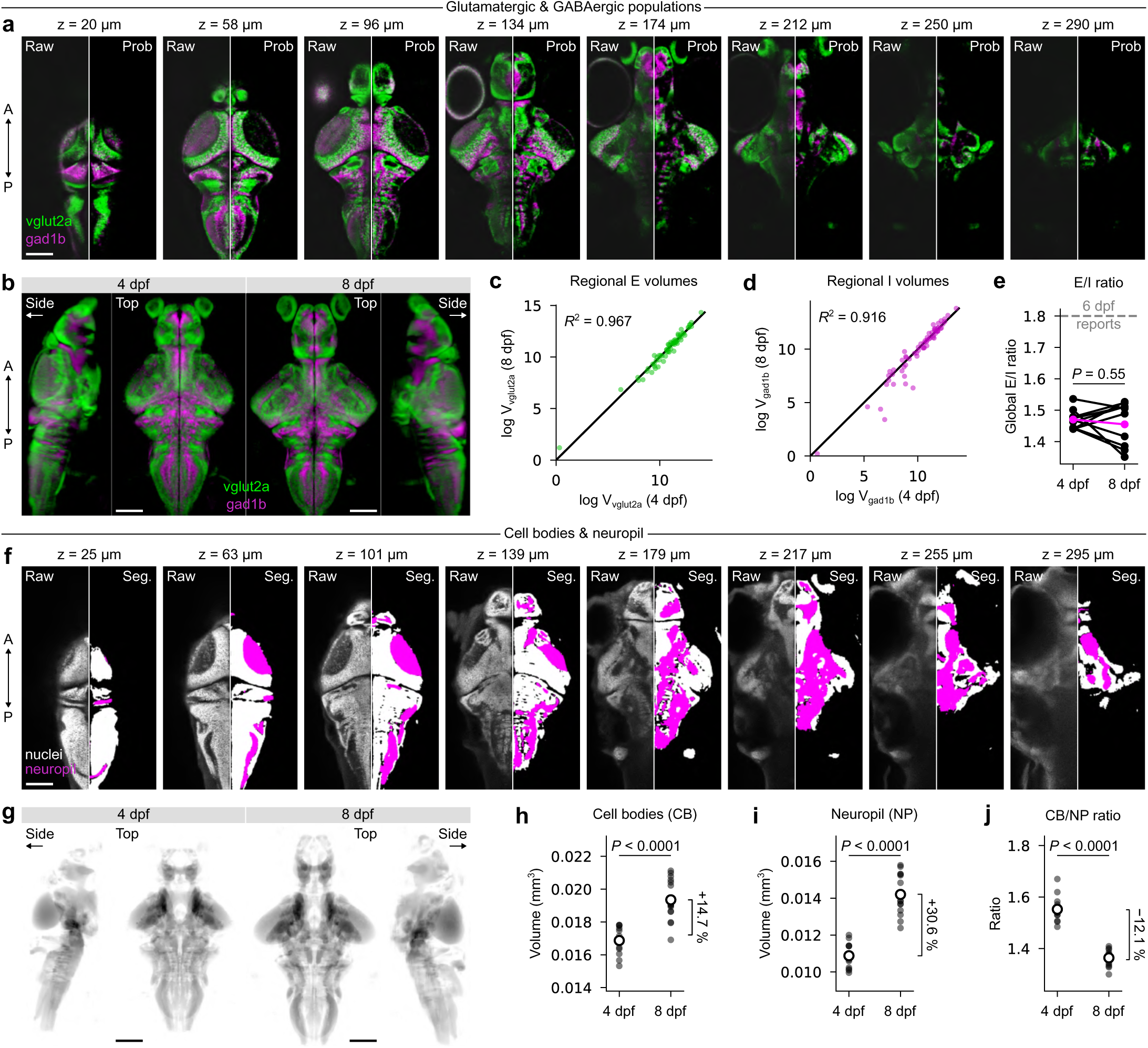
Supplementary analysis of neurotransmitters, neuropil, and cell bodies. (**a**) Averaged vglut2a (green) and gad1b (magenta) fluorescence microscopy images from 4 co-registered 8 dpf animals per channel (left), with corresponding probability densities inferred from individual binary segmentations (right). Depth relative to brain surface is indicated above each image. Notice how probability maps eliminate unspecific signal, including neuropil and eyes. (**b**) Maximum intensity projections of vglut2a and gad1b probability densities at 4 and 8 dpf, with pixelwise color contrast to accentuate specific neurotransmitter domains. Images are scaled arbitrarily in the vertical axis to facilitate comparison. (**c**-**d**) Comparison between log-transformed regional volumes of E and I neurons at 4 and 8 dpf. The black line corresponds to the identity line, and regression coefficients are indicated in each plot’s top left corner. (**e**) Global excitatory/inhibitory cell ratios estimated by summing probability maps across all brain regions. Different paired lines correspond to different train-test splits of the dataset, with each model trained on both 4 and 8 dpf data simultaneously to minimize bias. The colored line corresponds to a model trained on the full dataset. A dashed gray line indicates a recently reported ratio of 1.8 in 6 dpf larvae. *P* = 0.55, paired t-test. (**f**) Left, 6 dpf H2B template brain from mapZebrain; right, cell body (CB, white) and neuropil (NP, magenta) segmentations. Depth relative to brain surface is indicated above each image. (**g**) Mean intensity projections of neuropil regions from 4 and 8 dpf template brains. (**h**-**j**) Quantifications of CB and NP volumes, as well as their ratio in 4 and 8 dpf larvae. All quantities change significantly with age (*P <* 0.0001, Mann-Whitney U tests). Percentage change is indicated on the right of each plot. Scale bars: 100 *µ*m.

**Figure S6:**
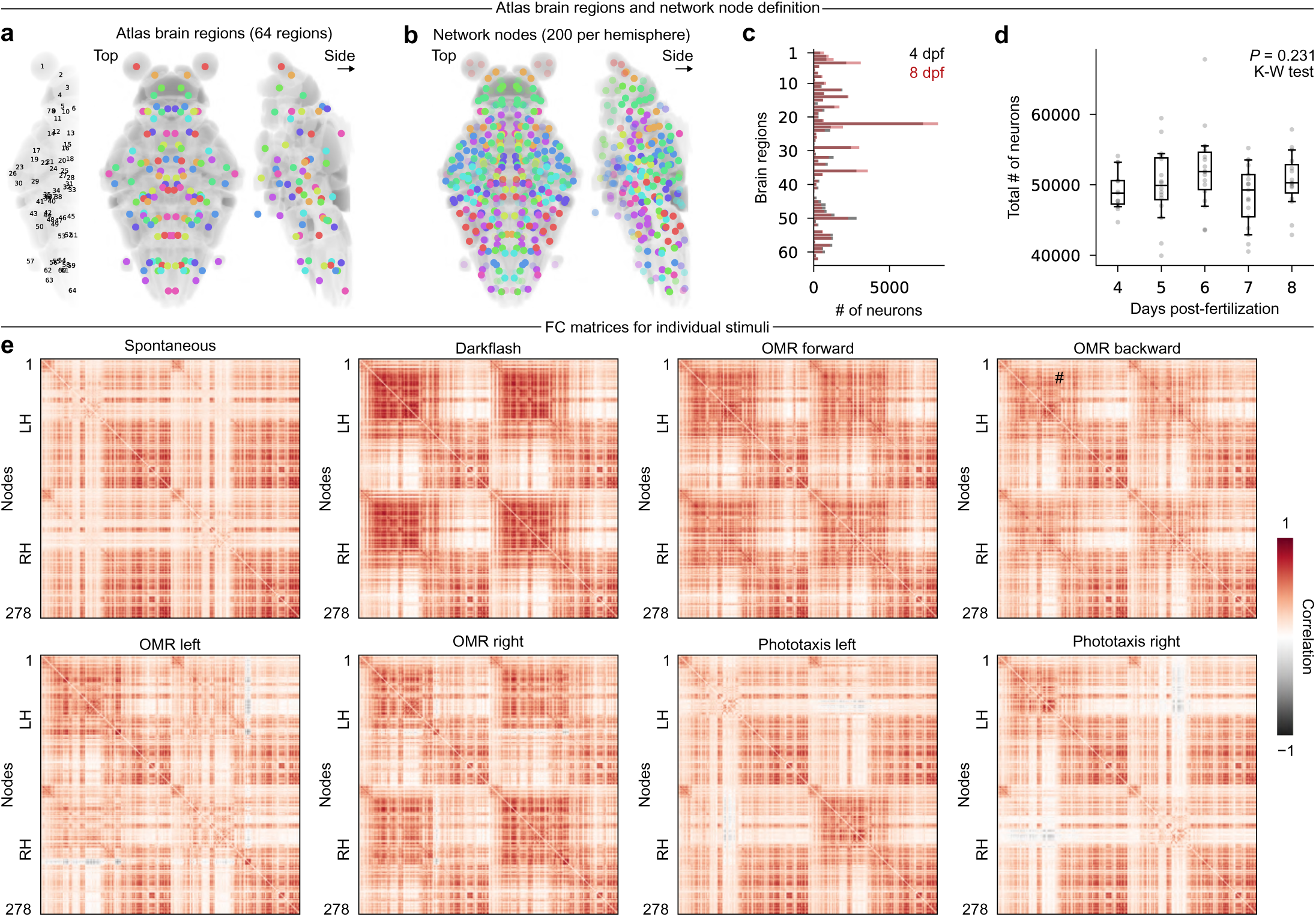
Network node definition and individual stimulus connectivity matrices. (**a**) Brain region centroids, sorted and numbered along the brain’s anteroposterior axis. (**b**) 200 network nodes per brain hemisphere with pseudocolors mapped to brain regions from the previous plot. Faded colors indicate nodes that have been excluded from FC analysis due to uneven sampling across animals. (**c**) Average number of neurons per brain region for 4 and 8 dpf animals. Notice generalized growth in anterior brain regions, and decrease in hindbrain regions. (**d**) Total number of segmented neurons per animal. Boxplot whiskers indicate 5th and 95th percentiles. No significant change in neuron count is observed (*P* = 0.231, K-W test). (**e**) Eight functional connectivity matrices calculated over different epochs of the 30-minute experiments. The first one corresponds to the first 15 minutes of spontaneous activity (provided as a visual reference), while the others correspond to individual visual stimuli.

**Figure S7:**
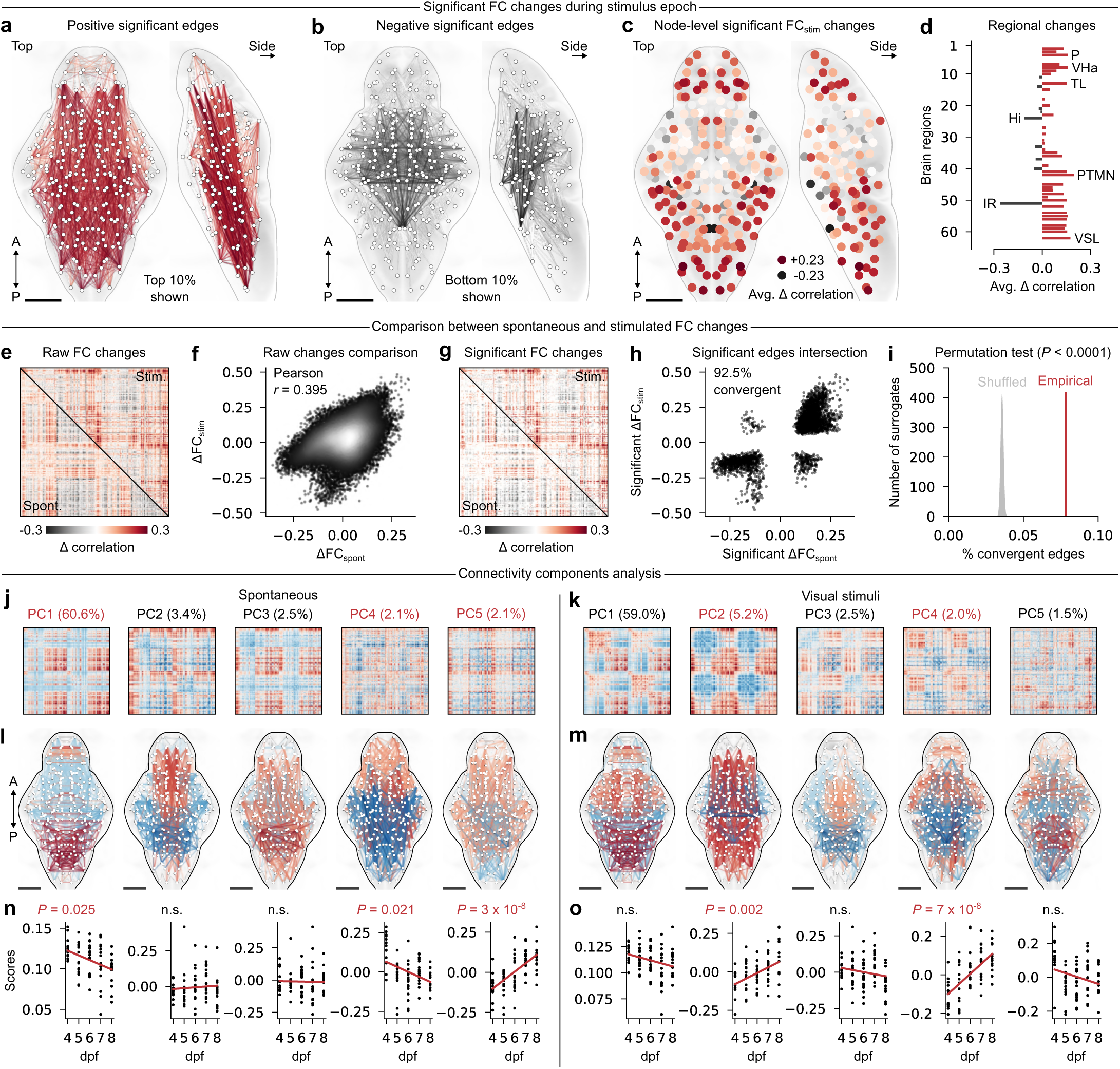
Supplementary analysis of functional connectivity. (**a**) Network visualization of the top 10% of positive changes for FC_stim_. (b) Similar visualization for the bottom 10% of negative changes. (**c**) Average significant FC_stim_ changes per network node. (**d**) Chart of average significant FC_stim_ changes per anatomical brain region. (**e**) Group-average FC difference (8 minus 4 dpf) for spontaneous (bottom) and visually-evoked (top) activity. (**f**) Comparison between the two previous matrix triangles. Pearson correlation is indicated in the upper left corner. (**g**) Statistically significant (*P <* 0.05, t-test) FC difference (8 minus 4 dpf) for spontaneous (bottom) and visually-evoked (top) activity. (**h**) Scatter plot of edge weights for edges that are significant in both spontaneous and stimulus epochs. 92.5% of edges share the same signs and are deemed convergent. (**i**) Percentage of convergent edges in empirical (red line) and shuffled (gray, 10000 permutations) FC matrices. The empirical score completely exceeds the null distribution, thus *P <* 0.0001. (**j**) Principal FCspont components identified using PCA. Red and blue colors indicate arbitrary positive and negative signs, respectively. The fraction of explained variance is indicated above each component, and those that vary developmentally are highlighted in red. (**k**) Similar plot for FC_stim_. (**l**) Anatomical network visualization of the above components. (**m**) Similar plot for FC_stim_. (**n**) Individual component scores, with linear regressions. Regression P-values that are significant after FDR correction are highlighted in red (correction over all 79 components). (**o**) Similar plot for FC_stim_. A, anterior; P, posterior; avg, average.

**Figure S8:**
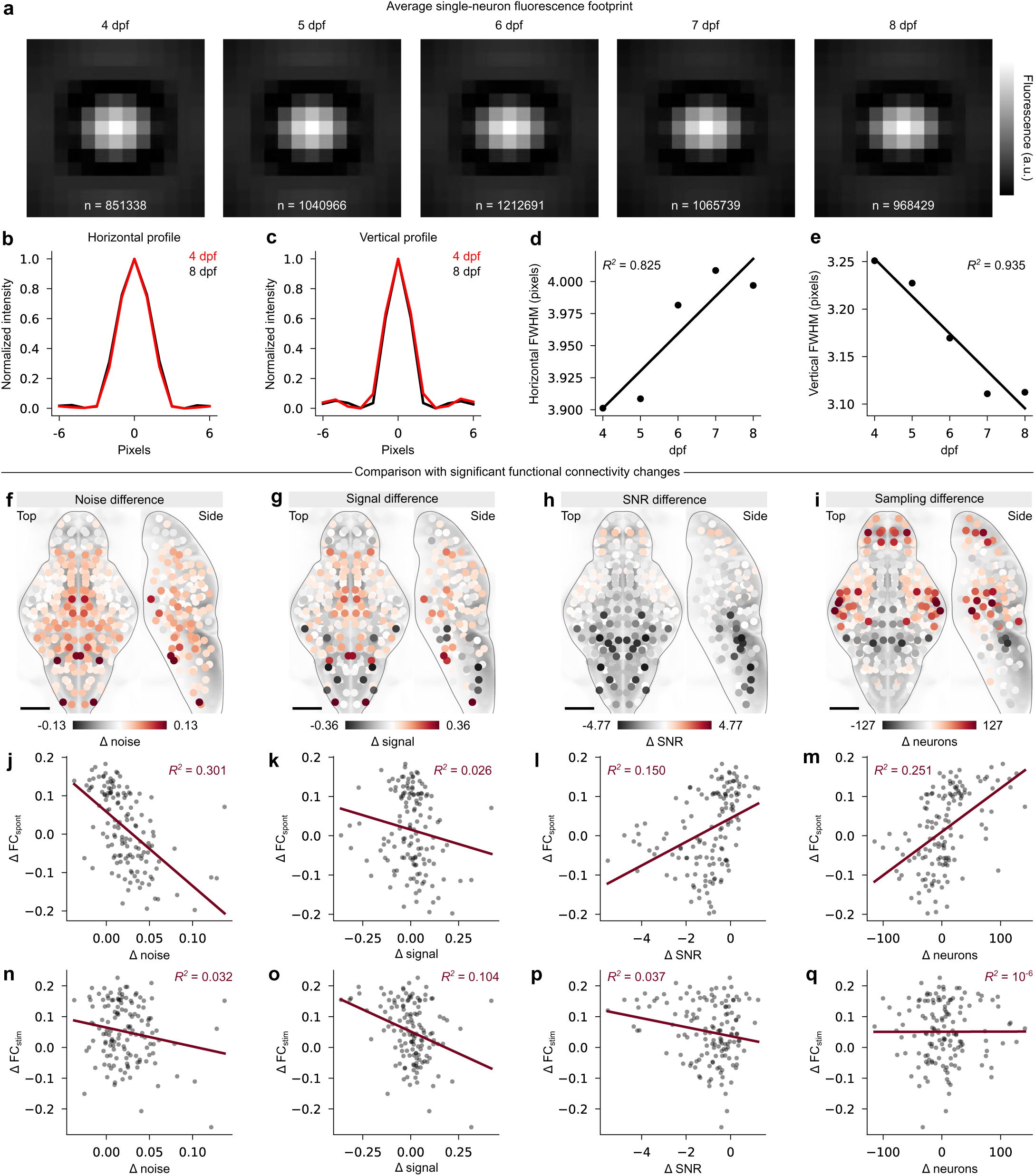
Signal properties. (**a**) Average single-neuron fluorescence footprints obtained by averaging in (13 *×* 13) crops around each detected local intensity maximum (fluorescent nuclei). The total number of neurons per age group is indicated on each image. Each neuron occupies a roughly 5 *×* 5 pixel footprint, providing ample subcellular resolution. (**b**) Average neuron intensity profiles along the horizontal axis at 4 (red) and 8 (black) dpf. (**c**) Similar plot for the vertical axis. Notice a slight vertical narrowing at 8 dpf. (**d**) Linear increase of the horizontal intensity profile. Because the pixel size does not change along this axis, this suggests a small developmental increase in nucleus size. (**e**) Linear decrease of the vertical intensity profile. In this case, this change reflects imaging parameters, as the vertical scanning amplitude was slightly increased in older animals (see Methods). (**f** -**i**) Node-level differences in various signal properties (noise level, signal intensity, signal-to-noise ratio (SNR), and sampling/neuron count) between 4 and 8 dpf. (**j**-**m**) Correlations between the four previous maps and node-averaged significant FCspont changes from Fig. 3i. Increased noise in mostly ventral regions explains no more than 30% variance in FC changes. (**n**-**q**) Similar plots for FC_stim_ changes. Explained variance is markedly lower for stimulus-evoked activity. Scale bars, 100 *µ*m.

**Figure S9:**
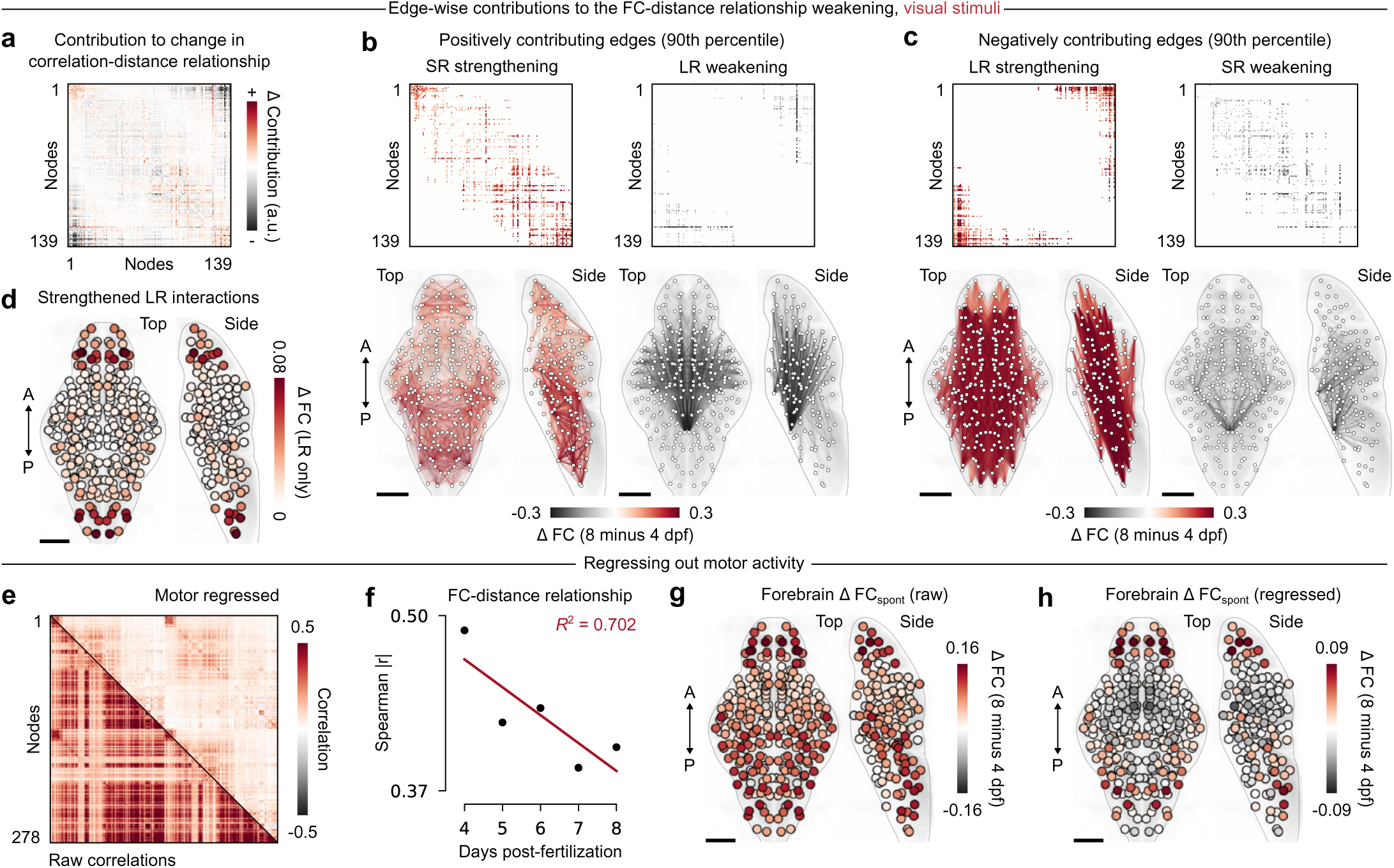
Supplementary analysis of long-range correlations. (**a**) Edge-wise contributions to the developmental change in the stimulus-evoked FC-distance relationship (8 minus 4 dpf). Positive or negative values denote edges that strengthen or weaken this relationship with development, respectively. (**b**) Top 10% positive edges from **a**, separated into short-range (SR; left) and long-range (LR; right) connections, (**c**) Bottom 10% negative edges from **a**, separated into LR (left) and SR (right) connections. (**d**) Node-level sum of FC changes associated with strengthened LR edges. In this case, mostly the pallium and caudal hindbrain are highlighted. (**e**) Group-averaged FC before (below) and after(above) regressing out a motor signal, defined as the average activity of M+ neurons, from the activity of each node. (**f**) Linear decay of the FCspont-distance relationship after motor regression. The decrease remains present after removing motor activity, despite a substantial decrease of correlations visible in **e**. (**g**) Node-averaged forebrain FCspont changes (8 minus 4 dpf), before motor regression. (**h**) Similar plot after motor regression. Positive correlation increases between the forebrain and several hindbrain nodes are preserved despite the removal of global motor activity. A, anterior; P, posterior. Scale bars, 100 *µ*m.

**Figure S10:**
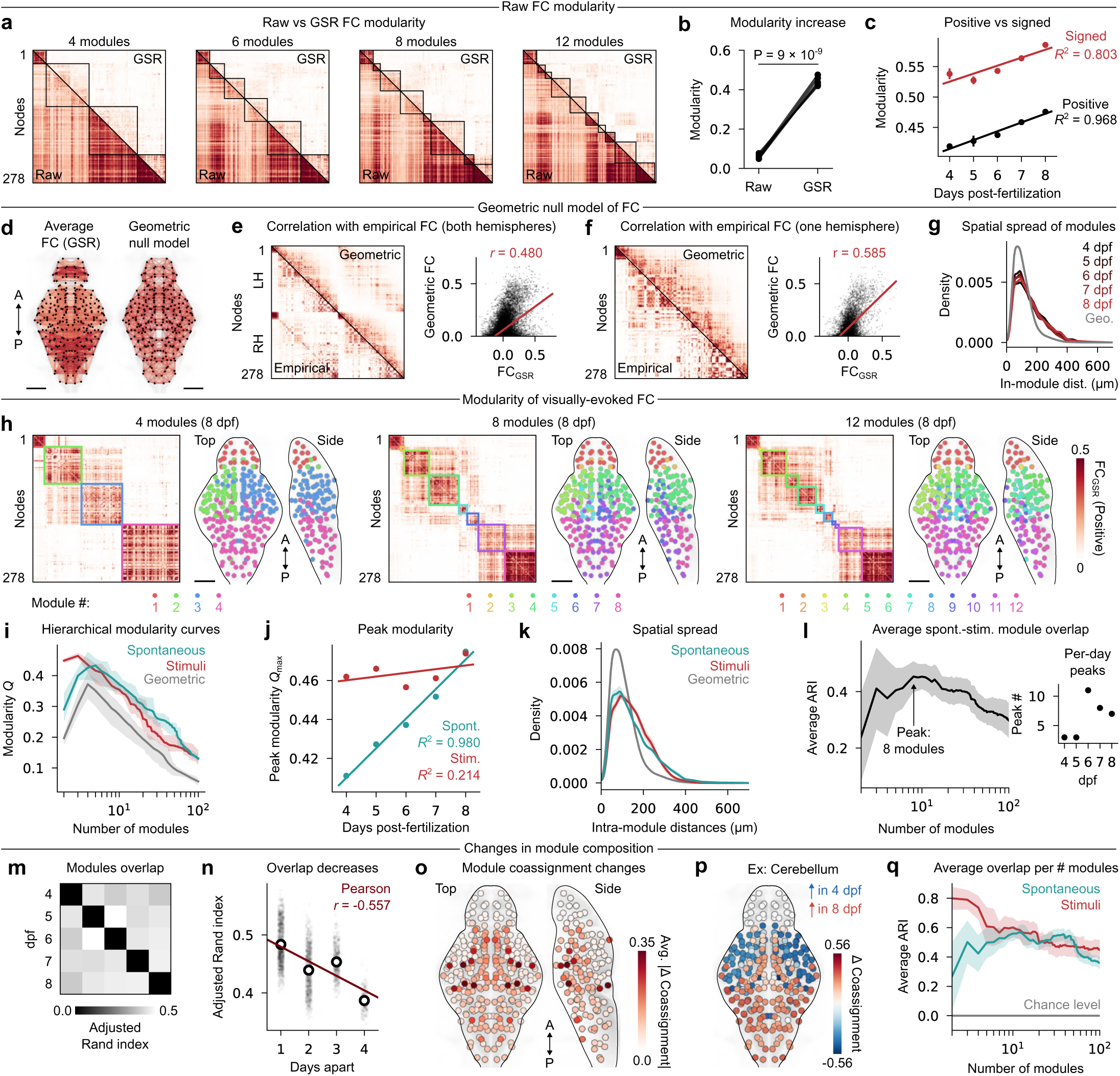
Supplementary modularity analyses. (**a**) FCspont matrices (bottom, raw correlations; top, global signal regression, GSR) sorted by 8 dpf modules. (**b**) Modularity evaluated using *γ*= 1 for raw and GSR group-averaged FC matrices. GSR significantly enhances modularity (*P* = 9 *×* 10*^−^*^9^, paired t-test). We refer to spontaneous FC with GSR as FCspont in the following panels for simplicity. (**c**) Modularity evaluated at *γ* = 1 on FCspont using either strictly positive (black) or signed (red) edge weights. Both modularity definitions exhibit a strong linear increase over development (regression coefficients annotated near each curve). (**d**) Network visualization of group-averaged FCspont (left) and geometric FC defined using exponentially decaying weights (right). (**e**) Left, comparison between group-averaged FCspont (bottom) and the average of all geometric FC models (top) used as modularity benchmarks. Right, correlation between both matrix triangles. (**f**) Similar to the previous panel, but FC matrices are restrained to a single hemisphere, thereby improving the fit. (**g**) Distributions of intra-module node distances, compiled across all partitions ranging from 2 to 100 modules. (**h**) Group-averaged 8 dpf FC_stim_ matrices (left) sorted by connectivity modules (right). (**i**) Average hierarchical modularity curves for spontaneous and stimuli periods, compared with modularity derived from geometric surrogates (gray line); the shaded regions indicate minimum and maximum values. (**j**) Linear increases of peak modularity values per age group for FC_spont_ (blue) and FCFC_stim_ (red). Linear regression coefficients are indicated on the plot. Stimulus-evoked modularity exhibits high variability and a negligible increase compared to spontaneous activity. (**k**) Distributions of intra-module node distances for both recording periods, averaged across age groups and compiled across all partitions ranging from 2 to 100 modules. (**l**) Left, Adjusted Rand index (ARI) curve quantifying the overlap between modules derived from spontaneous or stimulus-evoked activity. Overlap scores are averaged across all comparisons between developmental time points, peaking at 8 modules. Right, peak ARI values when comparing modules from the same time point only. Early time points show strong correspondence at coarse spatial scales, while later time points require further modular subdivisions to maximize correspondence between spontaneous and stimuli. (**m**) Anatomical overlap between modules from different developmental time points (spontaneous only), averaged across the full hierarchy. (**n**) Overlap scores from increasingly distant time points generally decrease (Pearson *r* = *−*0.557. Point distributions correspond to ARI values obtained from bootstrapped modules using the leave-one-out procedure. (**o**) Average absolute change in module coassignment probabilities per network node between 4 and 8 dpf, highlighting nodes that change module allegiance during development. (**p**) Example change in coassignment probabilties for the cerebellum, whose nodes are highlighted in the previous plot. (**q**) Average module overlap between all developmental time points for spontaneous (blue) and stimulus-evoked (red) activity. Visually-driven modules are similar at the coarsest scale (2-3 modules), whereas spontaneous modules show maximal correspondence at a deeper hierarchical level (10-15 modules). This suggests that spontaneous modules are comprised of similar building blocks that assemble, with developmentally varying strength, into the larger 4-5 modules where maximal modularity is observed. A, anterior; P, posterior. Scale bars, 100 *µ*m.

**Figure S11:**
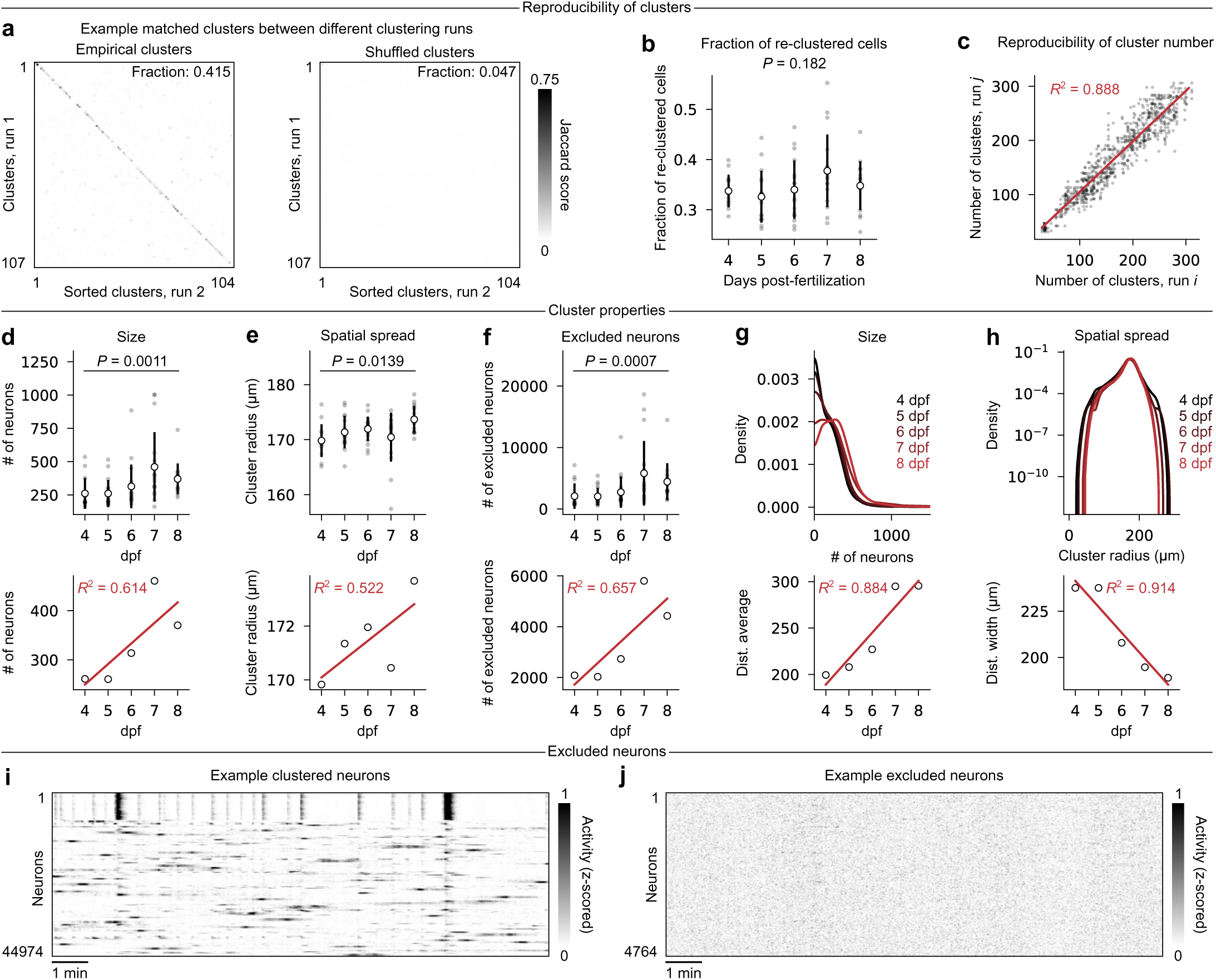
Properties of neuronal clusters. (**a**) Overlap between matched clusters (left, empirical; right, shuffled) from two repeated runs, quantified using Jaccard’s index. Clusters were optimally matched using a linear sum assignment optimization. The fraction of cells assigned to the same matched cluster on different runs is indicated in the top right corner. (**b**) Fraction of re-clustered cells (that is, cells belonging to matched clusters in different clustering runs) per animal. No significant developmental trend is observed (K-W test, *P* = 0.182), indicating comparable clustering reliability over the full dataset. (**c**) Reproducibility of the number of detected clusters per animal global *R*^2^ = 0.888 for all possible pairwise comparisons between the 5 clustering runs of each animal, indicating high reproducibility. (**d**-**f**) Top, average cluster size, spatial spread, and number of excluded neurons per animal (averaged over all clusters and five clustering runs). Distributions exhibit significant shifts over development (K-W test, P-values indicated on each plot after FDR correction). Bottom, linear regressions of the above distribution means, with regression coefficients indicated on each plot. (**g**) Top, probability densities of cluster size for each age group, combining all detected clusters over different clustering runs. Bottom, linear regression of distribution averages, which exhibit a monotonous developmental increase (*R*^2^ = 0.884). (**h**) Top, probability densities of cluster spatial spread for each age group, combining all detected clusters over different clustering runs. Bottom, linear regression of distribution widths (measured at a 10*^−^*^7^ threshold) which exhibit a monotonous developmental decrease (*R*^2^ = 0.914). (**i**) Raster plot of neuronal activity, sorted per functional clusters, from one 8 dpf animal. (**j**) Raster plot of excluded neurons from the same 8 dpf animal.

**Figure S12:**
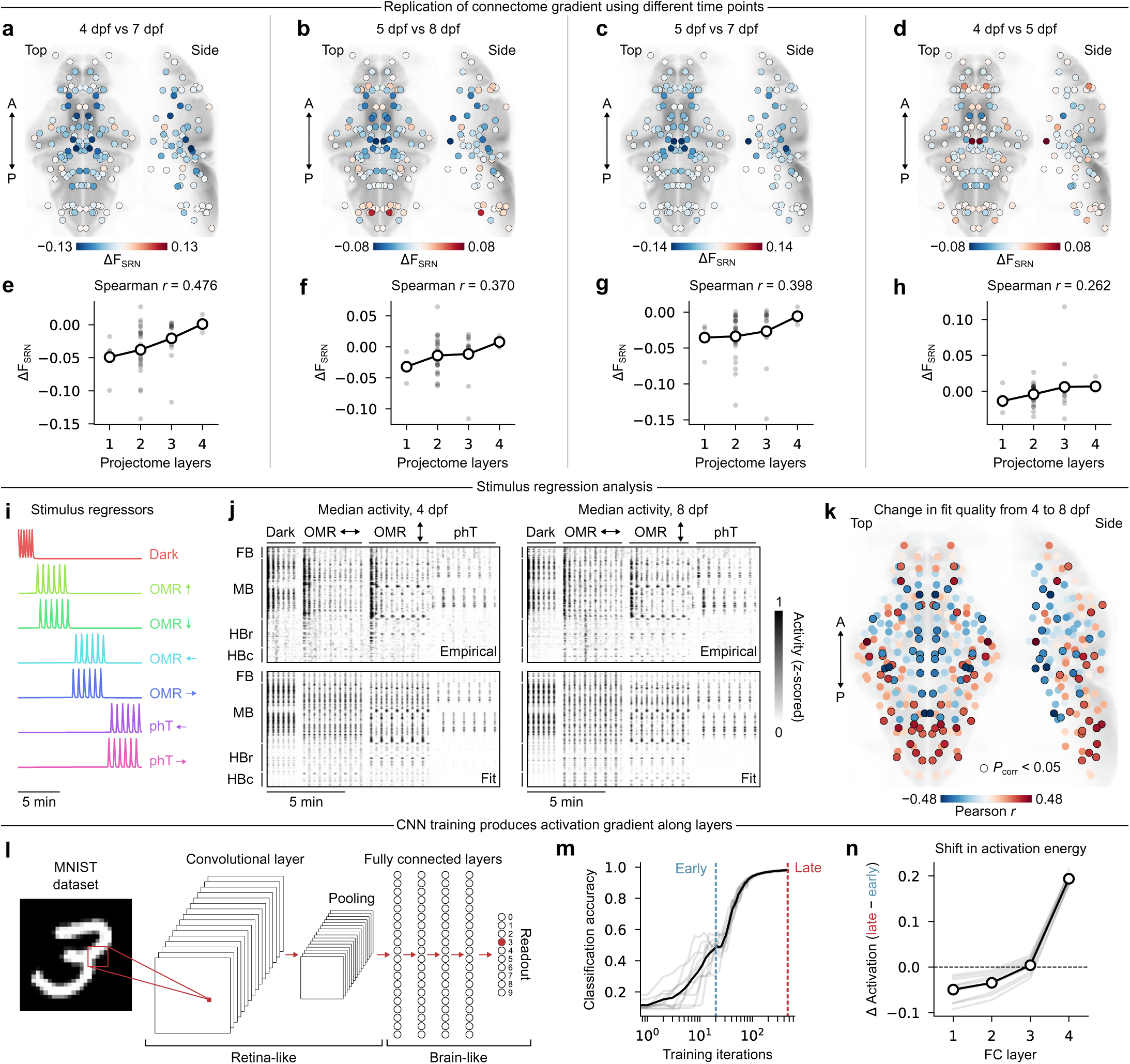
Supplementary analysis of visual stimuli. (**a**-**d**) Anatomical heatmaps of variation in the fraction of SRNs per brain region between different time points, indicated above each plot. (**e**-**h**) Change in the fraction of SRNs across the four projectome layers, associated with the plots above; Spearman correlation coefficients indicated above each plot. (**i**) Regressors (binary stimulus vectors convolved with an exponential kernel, *τ* = 3.5 s) for seven visual stimuli, excluding light, which drives very little activity at the mesoscale, compared to other stimuli. (**j**) Top row, median node-averaged activity during visual stimuli at 4 (left) and 8 (right) dpf; anatomical nodes are sorted by 4 major brain segments: forebrain (FB), midbrain (MB), rostral (HBr) and caudal (HBc) hindbrain; and stimulus blocks are annotated above; bottom, predicted activity obtained by fitting the regressors to the empirical data. (**k**) Linear regression of regressor explained variance against developmental stage, combining all 79 animals, and averaged across hemispheres; positive correlations indicate nodes whose regressor fits improve linearly over days, suggesting the strengthening of visually-induced activity, and vice-versa for negative correlations; nodes with significant correlations are highlighted with black outlines (*P <* 0.05, FDR-corrected). Note that this analyses evaluates linear changes across all time points, and avoids endpoint comparisons. (**l**) Schematization of the MNIST digit classification task and CNN architecture. (**m**) Classification accuracy over training iterations for 10 independently trained models (dark line indicates the average). Early (20 iterations) and late (400 iterations) training stages are indicated by dashed vertical lines. (**n**) Change in average unit activation in each of the fully connected (FC) layers, comparing early vs late training stages (dotted dark line indicates the average). Note a decrease in layers 1-2 and an increase in layers 3-4, mimicking the gradient observed along the zebrafish projectome.

**Figure S13:**
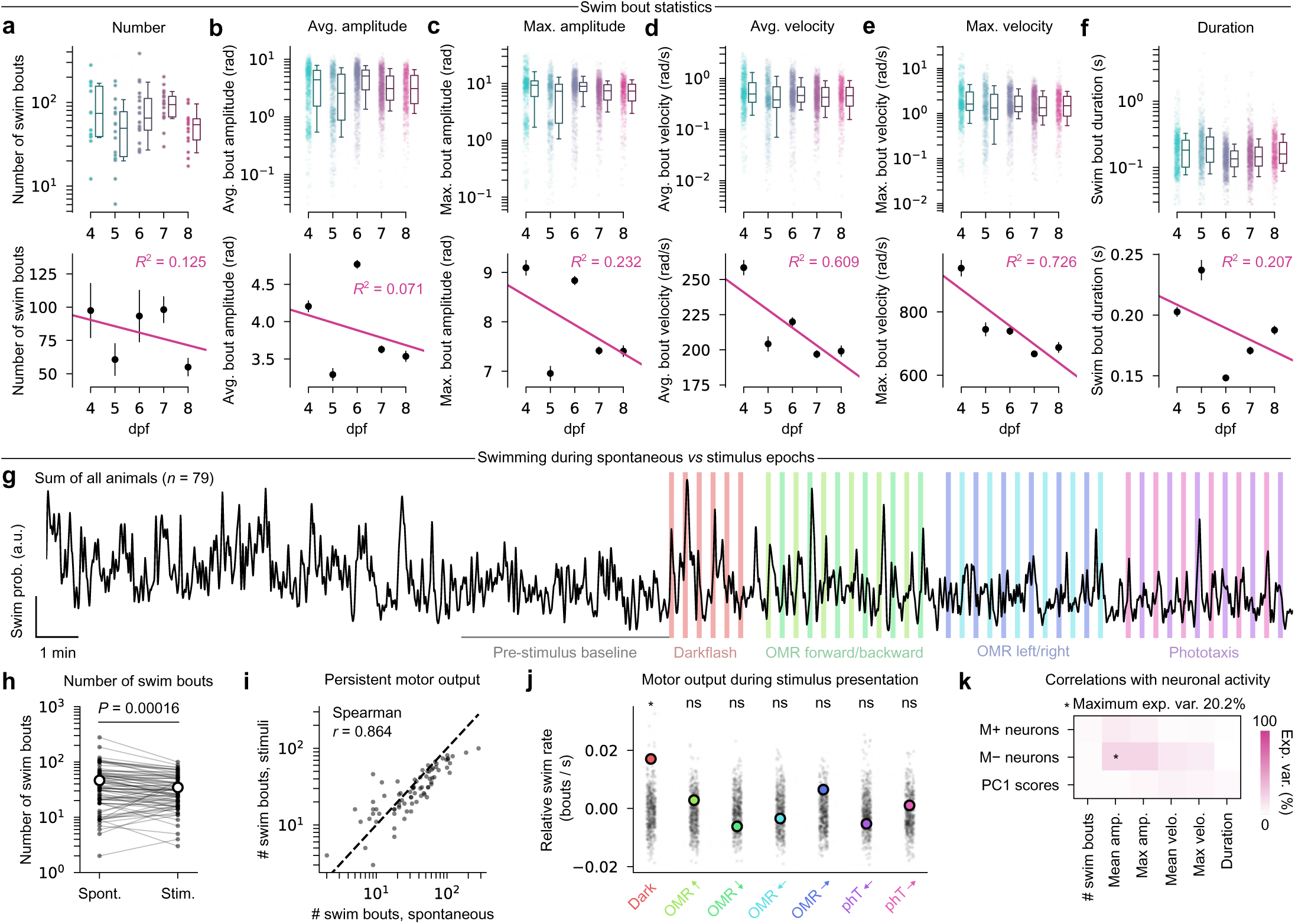
Supplementary analysis of head-restrained behavior. (**a**-**f**) Top row, distributions of various behavioral parameters (number, average amplitude, maximum amplitude, average velocity, maximum velocity, and duration of swim bouts) per developmental time point. Boxplot whiskers indicate 10th and 10th percentiles. Notice a bimodal distribution of swim amplitudes (**b**-**c**) at 4-5 dpf, indicating a high number of low-amplitude swimming events that is markedly reduced in older animals. Bottom row, linear regressions of distribution averages from the above plots. Although most regression coefficients are low, all swimming parameters exhibit a developmental decrease, indicating an overall trend of reduced locomotor output, both in number and intensity, in older animals. (**g**) Combined swim probability over the full experiment duration from all 79 animals. Shaded vertical bars indicate periods of stimulus presentation, and a gray line indicates a 5-minute pre-stimulus period used as a baseline for subsequent measures. (**h**) Comparison between the number of swim bouts during spontaneous and stimulus periods. Swimming is slightly but significantly reduced during the second half of the experiment (*P* = 0.00016, paired t-test). (**i**) Scatter plot of number of swim bouts during both halves of the experiment, with a dashed line corresponding to the identity line. Each animal’s spontaneous swimming is highly predictive of swimming during the stimulus period (Spearman’s *r* = 0.864), which suggests that the level of motor activity is preserved throughout the full experiment. (**j**) Average swim rates (in swim bouts per second), relative to the pre-stimulus baseline, during each visual stimulus. Point distributions correspond to null swim rates measured by randomly shuffling the stimulus presentation windows during the second half of the experiment. Only darkflash stimuli elicit swimming events more than their expected spontaneous levels (*P <* 0.05 after FDR correction). This can be perceived in panel **g** as large peaks during the first two presentations of the stimulus, with putative habituation rapidly taking over in subsequent repetitions and stimuli. (**k**) Correlations between neuronal-activity-derived quantities (number of M+ and M*−* neurons from Fig. 7g,j and individual PC1 scores from Fig. 7p) and session-averaged behavioral parameters from **a**-**f**. The maximum explained variance barely exceeds 20%, which suggests that the amount of swimming has little effect on neuronal representation. PC1 scores in particular are almost completely decorrelated from behavioral parameters.

**Figure S14:**
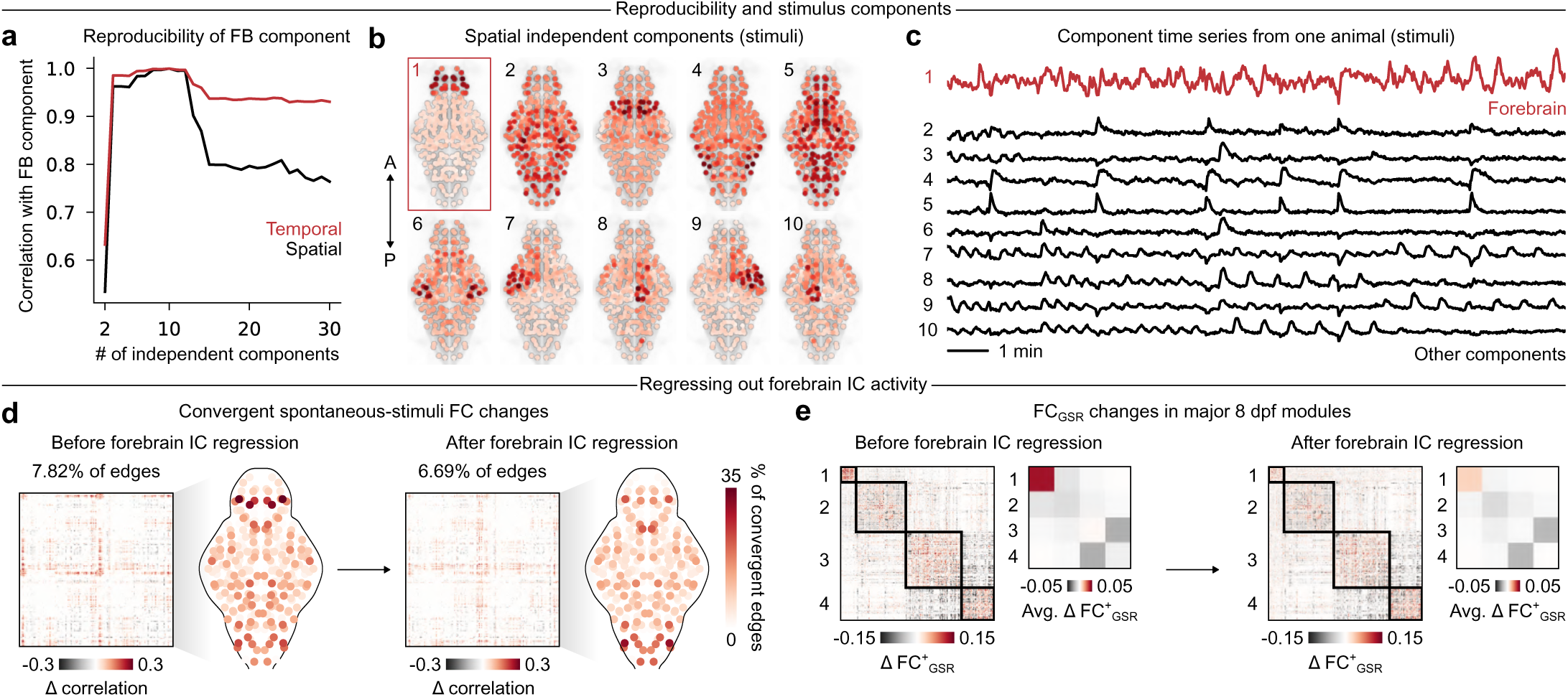
Supplementary analyis of the forebrain independent component. (**a**) Correlation between the forebrain component analyzed in Fig. 8 and homologous components identified using a different total number of component. The forebrain IC emerges beyond 3 components (indicated by an abrupt jump in correlation), and remains highly correlated until roughly 15 components are reached. Beyond this point, spatial correlations decrease, as the higher number of allowed components leads to further spatial subdivisions of the forebrain (not shown). The correlations between component time courses remain high, however, as the spontaneous activity pattern, which is present in all forebrain subdivisions, is recovered regardless of the exact spatial footprint. (**b**) 10 group-level spatial components uncovered through spatial ICA during the stimulus epoch; the forebrain (FB) component is accentuated in red. (**c**) Time courses of the 10 previous components; the FB signal is accentuated in red. Note the presence of periodic visually driven activity in several components, which is completely absent from the forebrain signal. (**d**) Number of convergent FC_spont_-FC_stim_ changes, before and after regressing out the forebrain IC time course from each node’s activity. The number of convergent changes is markedly reduced in forebrain nodes, indicating that convergent changes may represent the strengthening of the forebrain component during both epochs. (**e**) FCspont changes organized by main 8 dpf modules, before and after regressing out the forebrain signal. Connectivity strengthening in the forebrain module is nearly abolished following regression. Hence, the forebrain’s intrinsic functional connectivity can be attributed to a large degree to the forebrain IC. A, anterior; P, posterior.

### Supplementary tables

**Table 1:** List of 64 mutually exclusive and collectively exhaustive brain regions from mapZebrain used throughout this study. Regions are ordered from anterior to posterior according to the centroids of their 3D binary masks.

| ID | Name | Acronym | ID | Name | Acronym |
| --- | --- | --- | --- | --- | --- |
| 1 | Olfactory Epithelium | OE | 33 | Superior Raphe | SR |
| 2 | Olfactory Bulb | OB | 34 | Sup. Dorsal Med. Obl. Stripe 2&3 | SDMOS2-3 |
| 3 | Subpallium | S | 35 | Locus Coeruleus | LC |
| 4 | Pallium | P | 36 | Cerebellum | Cb |
| 5 | Dorsal Habenula | DHa | 37 | Sup. Ventral Med. Obl. (Remaining) | SVMOR |
| 6 | Epiphysis | E | 38 | Caudal Hypothalamus | Hc |
| 7 | Ventral Entopeduncular Nucleus | VEN | 39 | Sup. Dorsal Med. Obl. Stripe 4 | SDMOS4 |
| 8 | Ventral Habenula | VHa | 40 | Anterior (Dorsal) Trigeminal Motor Nucleus | ATMN |
| 9 | Eminentia Thalami (Remaining) | ETR | 41 | Sup. Dorsal Med. Obl. Stripe 5 | SDMOS5 |
| 10 | Preoptic Region | Po | 42 | Posterior (Ventral) Trigeminal Motor Nucleus | PTMN |
| 11 | Ventral Thalamus (Alar part) | VT | 43 | Glossopharyngeal Ganglion | GG |
| 12 | Dorsal Thalamus Proper | DT | 44 | Octaval Ganglion | OG |
| 13 | Torus Longitudinalis | TL | 45 | Int. Dorsal Med. Obl. Stripe 1 | IntDMOS1 |
| 14 | Pretectum (Alar part) | Pr | 46 | Int. Dorsal Med. Obl. Stripe 2&3 | IntDMOS2&3 |
| 15 | Posterior Tuberculum | PT | 47 | Int. Dorsal Med. Obl. Stripe 4 | IntDMOS4 |
| 16 | Rostral Hypothalamus | Hr | 48 | Int. Dorsal Med. Obl. Stripe 5 | IntDMOS5 |
| 17 | Tectal Neuropil | OTn | 49 | Int. Ventral Med. Obl. (Remaining) | IntVMOR |
| 18 | Medial Tegmentum (Remaining) | MTR | 50 | Medial Octavolateralis Nucleus | MON |
| 19 | Diffuse Nucleus of the Inferior Lobe | DNIL | 51 | Inferior Raphe | IR |
| 20 | Nucleus Med. Long. Fasc. | nMLF | 52 | Facial Motor Nucleus | FMN |
| 21 | Lateral Tegmentum | LT | 53 | Abducens Motor Nucleus | AMN |
| 22 | Periventricular Layer (Optic tectum) | OT | 54 | Inf. Ventral Med. Obl. (Remaining) | InfVMOR |
| 23 | Torus Semicircularis | TS | 55 | Inf. Dorsal Med. Obl. Stripe 4 | InfDMOS4 |
| 24 | Intermediate Hypothalamus (Remaining) | Hi | 56 | Inf. Dorsal Med. Obl. Stripe 5 | InfDMOS5 |
| 25 | Oculomotor Nucleus | ON | 57 | Posterior Lateral Line Ganglion | PLLG |
| 26 | Trigeminal Ganglion | TG | 58 | Inf. Dorsal Med. Obl. Stripe 2&3 | InfDMOS2-3 |
| 27 | Trochlear Motor Nucleus. | TMN | 59 | Inf. Dorsal Med. Obl. Stripe 1 | InfDMOS1 |
| 28 | Pituitary | Pit | 60 | Vagus Motor Nucleus | VMN |
| 29 | Nucleus Isthmi | NI | 61 | Inferior Olive | IO |
| 30 | Anterior Lateral Line Ganglion | ALLG | 62 | Vagal Sensory Lobe | VSL |
| 31 | Interpeduncular Nucleus | IPN | 63 | Lateral Reticular Nucleus | LRN |
| 32 | Sup. Dorsal Med. Obl. Stripe 1 (Remaining) | SDMOS1R | 64 | Area Postrema | AP |

